# Multivariate genomic analysis of 1.5 million people identifies genes related to addiction, antisocial behavior, and health

**DOI:** 10.1101/2020.10.16.342501

**Authors:** Richard Karlsson Linnér, Travis T. Mallard, Peter B. Barr, Sandra Sanchez-Roige, James W. Madole, Morgan N. Driver, Holly E. Poore, Andrew D. Grotzinger, Jorim J. Tielbeek, Emma C. Johnson, Mengzhen Liu, Hang Zhou, Rachel L. Kember, Joëlle A. Pasman, Karin J.H. Verweij, Dajiang J. Liu, Scott Vrieze, COGA Collaborators, Henry R. Kranzler, Joel Gelernter, Kathleen Mullan Harris, Elliot M. Tucker-Drob, Irwin Waldman, Abraham A. Palmer, K. Paige Harden, Philipp D. Koellinger, Danielle M. Dick

## Abstract

Behaviors and disorders related to self-regulation, such as substance use, antisocial conduct, and ADHD, are collectively referred to as *externalizing* and have a shared genetic liability. We applied a multivariate approach that leverages genetic correlations among externalizing traits for genome-wide association analyses. By pooling data from ~1.5 million people, our approach is statistically more powerful than single-trait analyses and identifies more than 500 genetic loci. The identified loci were enriched for genes expressed in the brain and related to nervous system development. A polygenic score constructed from our results captures variation in a broad range of behavioral and medical outcomes that were not part of our genome-wide analyses, including traits that until now lacked well-performing polygenic scores, such as opioid use disorder, suicide, HIV infections, criminal convictions, and unemployment. Our findings are consistent with the idea that persistent difficulties in self-regulation can be conceptualized as a neurodevelopmental condition.

## Main

Behaviors and disorders related to self-regulation, such as substance use disorders or antisocial behaviors, have far-reaching consequences for affected individuals, their families, communities, and society at large^1,2^. Collectively, this group of correlated traits are classified as *externalizing*^3^. Twin-family studies have demonstrated that externalizing liability is highly heritable (~80%)^4,5^, suggesting it will be as tractable to gene discovery as other complex traits or medical conditions^6^. To date, however, there have been no large-scale molecular genetic studies that utilize the extensive degree of genetic overlap among externalizing traits to aid gene discovery, as most studies have focused on individual disorders or diseases^7^. But for many high-cost, high-risk externalizing behaviors – opioid use disorder and suicide attempts being salient examples – there are too few cases available with genome-wide data to yield sufficient power for gene discovery^8,9^.

A complementary strategy to the single-disease approach is to study the shared genetic architecture across traits in multivariate analyses, which boosts statistical power by pooling data across genetically correlated traits^10^. Multivariate approaches can utilize summary statistics from genome-wide association studies (GWAS), which are now widely available, to allow for the discovery of connections between phenotypes not naturally studied together because they span different domains, fields of study, or life stages. Conveniently, by adjusting for sample overlap, novel statistical methods can attain an even greater effective sample size by efficiently utilizing observations from overlapping studies. Elucidating the shared genetic basis of externalizing liability has the potential to advance our understanding of the biological processes related to behavioral undercontrol, and enables mapping the pathways by which genetic risk and socio-environmental factors interact to contribute to the development of different externalizing outcomes.

Here, we applied genomic structural equation modeling (Genomic SEM) to summary statistics from GWAS on multiple forms of externalizing behavior for which large samples were available^10^. This approach was grounded in the existing literature showing shared genetic liability across numerous externalizing disorders and with non-psychiatric variation in externalizing behavior^5,11^. We posited that applying this multivariate approach would lead to the identification of genetic variants associated with a broad array of externalizing phenotypes, as well as related behavioral, social, and medical outcomes that were not directly included in our genome-wide association analysis.

## Results

### Multivariate analysis of seven externalizing phenotypes identifies numerous genetic associations with a general liability to externalizing

Following our preregistered analysis plan (https://doi.org/10.17605/OSF.IO/XKV36, Supplementary Information section 1), we collated GWAS summary statistics from externalizing-related disorders and behaviors, with our final analysis using data from seven externalizing phenotypes with sample sizes >50,000 (**Table 1**): (1) attention-deficit/hyperactivity disorder (ADHD), (2) problematic alcohol use (ALCP), (3) lifetime cannabis use (CANN), (4) age at first sexual intercourse (FSEX), (5) number of sexual partners (NSEX), (6) general risk tolerance (RISK), and (7) lifetime smoking initiation (SMOK). All samples were of European ancestry. The GWAS protocol is described in Supplementary Information section 2 (**Supplementary Tables 1–4**).

**Table 1.** Summary of seven externalizing-related disorders and behaviors with GWAS summary statistics (*N* > 50,000)

| Phenotype (abbreviation) | $N$ | $h^2$ ($SE$) | $\lambda_{GC}$ | Mean $\chi^2$ | Intercept | Ratio | Reference |
| --- | --- | --- | --- | --- | --- | --- | --- |
| Attention-deficit/hyperactivity disorder (ADHD) | 53,293 | .235 (.015) | 1.253 | 1.297 | 1.034 | .113 | <sup>12</sup> |
| Problematic alcohol use (ALCP) | 164,121 | .055 (.004) | 1.149 | 1.174 | 1.013 | .073 | <sup>13,14</sup> |
| Lifetime cannabis use (CANN) | 186,875 | .066 (.004) | 1.230 | 1.267 | 1.026 | .098 | <sup>15</sup> |
| Age at first sexual intercourse (FSEX) | 357,187 | .115 (.004) | 1.623 | 1.869 | 1.036 | .041 | <sup>16</sup> |
| Number of sexual partners (NSEX) | 336,121 | .097 (.004) | 1.492 | 1.682 | 1.027 | .041 | <sup>16</sup> |
| General risk tolerance (RISK) | 426,379 | .053 (.002) | 1.372 | 1.461 | 1.019 | .041 | <sup>16</sup> |
| Lifetime smoking initiation (SMOK) | 1,251,809 | .078 (.002) | 2.328 | 3.152 | 1.126 | .058 | <sup>17</sup> |

Notes: The statistics reported in this table were all estimated with LD Score regression<sup>18</sup>. Heritability ( $h^2$ ) is on the observed scale<sup>18</sup>. $\lambda_{GC}$ is the median $\chi^2$ statistic divided by the expected median of the $\chi^2$ distribution with 1 degree of freedom<sup>19</sup>. Mean $\chi^2$ is the average $\chi^2$ statistic. Intercept is the estimated LD Score regression intercept. Ratio measures stratification bias, defined as $(\text{Intercept} - 1) / (\text{Mean } \chi^2 - 1)$ <sup>18</sup>.

Consistent with twin studies^4,5^, the genetic correlations among the seven discovery phenotypes were moderate to high (**Figure 1A** and **Supplementary Table 5**). Using Genomic SEM^10^ (Supplementary Information section 3), which is unbiased by sample overlap and differences in sample sizes in the discovery phenotypes, we formally modeled the genetic covariances among the seven phenotypes and found that a common factor model fits the data best. This common factor, which we refer to as *EXT*, captures a shared genetic liability to the seven externalizing traits that we included in our analyses (**Figure 1B** and **Supplementary Table 7**).

**Figure 1.**
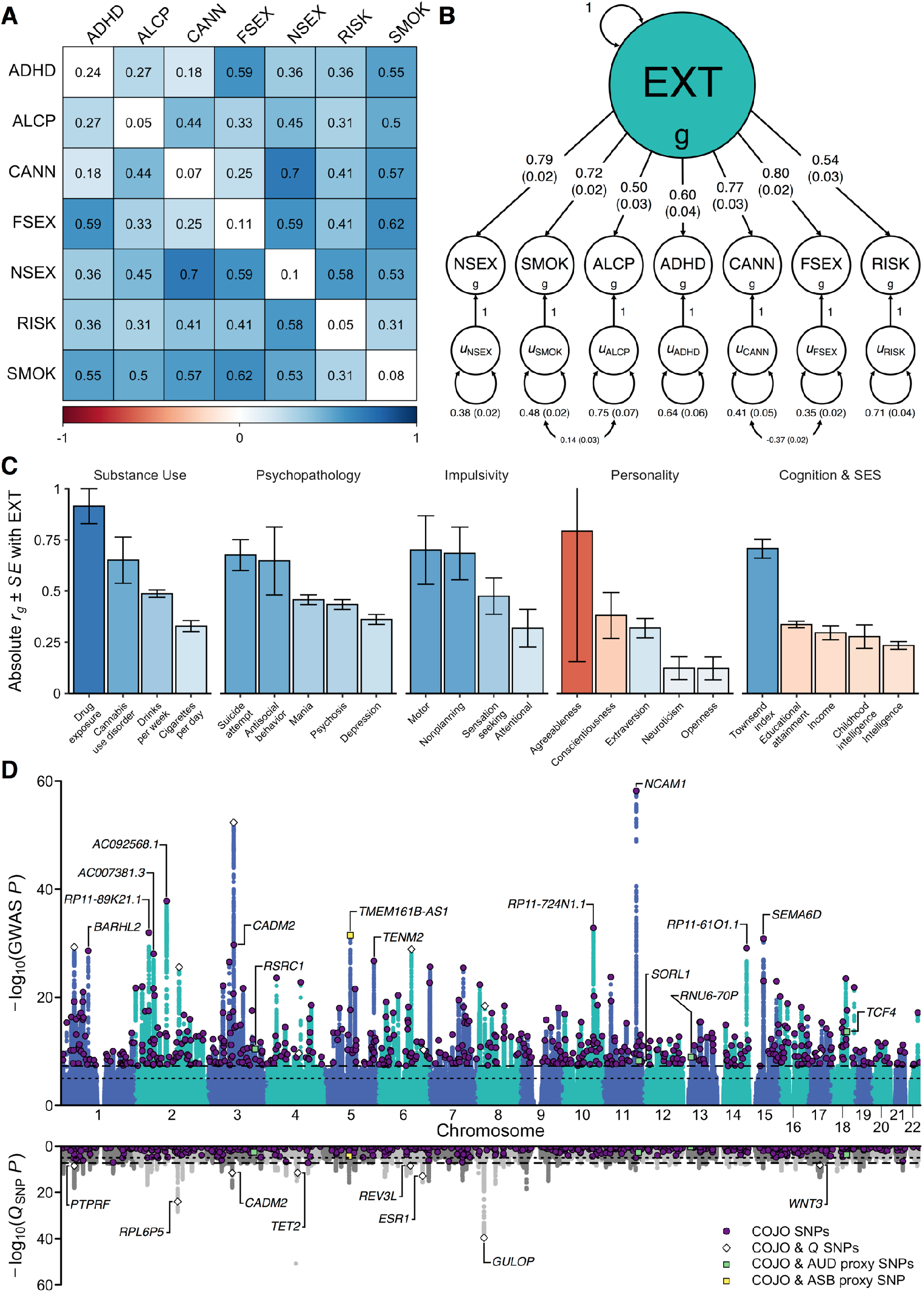
Multivariate genome-wide analyses with Genomic SEM. (**A**) Pair-wise genetic correlations (*r_g_*) among seven discovery phenotypes, with observed-scale SNP heritabilities (*h^2^*) on the diagonal. (**B**) Path diagram of a confirmatory factor model estimated with Genomic SEM. The parameter estimates were standardized, and standard errors are presented in parentheses. (**C**) Absolute value genetic correlations, |*r_g_*|, between the genetic externalizing factor (*EXT*) and phenotypes selected to establish convergent and discriminant validity, where blue and red bars represent positive and negative genetic correlations, respectively. Standard errors are presented as error bars. (**D**) GWAS associations (top panel) and *Q*_SNP_ tests of heterogeneity (bottom panel) for *EXT*. Purple dots represent 579 *EXT* lead SNPs that are conditionally and jointly associated (COJO) at genome-wide significance (two-sided test *P* < 5×10^-8^). White diamonds represent eight of the 579 SNPs that also show significant *Q*_SNP_ heterogeneity. Four green and one yellow squares represent five out of the 579 SNPs that also were Bonferroni-significant proxy-phenotype associations with alcohol use disorder (AUD) and antisocial behavior (ASB), respectively. ADHD is attention deficit hyperactivity disorder, ALCP is problematic alcohol use, CANN is lifetime cannabis use, EXT is externalizing, FSEX is age at first sex, NSEX is number of sexual partners, RISK is general risk tolerance, SMOK is lifetime smoking initiation.

We then extended Genomic SEM to estimate genetic correlations between *EXT* and 92 preregistered phenotypes with GWAS summary statistics that were not included among the seven discovery phenotypes (**Extended Data Fig. 1** and **Supplementary Table 8**). The genetic correlations indicate convergent and discriminant validity of the common *EXT* factor (**Figure 1C**): As anticipated, *EXT* showed strong genetic correlations with drug exposure (*r_g_* = .91), antisocial behavior (*r_g_* = .65), motor impulsivity (*r_g_* = .70), failures to plan (*r_g_* = .70), and (lack of) agreeableness (*r_g_* = −.79), a personality trait characterized by kindness and cooperativeness that has been found to be low in individuals displaying antisocial behavior. *EXT* was also strongly correlated with suicide attempts (*r_g_* = .68). *EXT* showed more modest inverse correlations with educational attainment (*r_g_* = −.32) and intelligence (*r_g_* = −.23), indicating that the latent factor is not simply reflecting genetic influences on cognitive ability. Finally, there was a strong genetic correlation with the Townsend index (*r_g_* = .71), a measure of neighborhood deprivation that reflects high concentrations of unemployment, household overcrowding, and low concentrations of home- and car-ownership^20^.

We next used Genomic SEM^10^ to perform a GWAS on the shared genetic liability *EXT* (**Figure 1D** and **Extended Data Fig. 2**) (Supplementary Information section 3.4). This analysis estimated single-nucleotide polymorphism (SNP) associations directly with the *EXT* factor, with an effective sample size of *N* = 1,492,085 individuals. These analyses are different in their approach and substantially increase sample size, statistical power, and the range of findings compared to previous work^21^ (Supplementary Information section 2.2.1). After applying conditional and joint multiple-SNP analysis (COJO) on a set of near-independent, genome-wide significant (two-sided test *P* < 5×10^-8^) lead SNPs^22^, we identified 579 conditionally and jointly associated SNPs (**Supplementary Table 9**), meaning they were significantly associated with *EXT* even after statistically adjusting for each other and other lead SNPs. Of the 579 *EXT* SNPs and their correlates within linkage disequilibrium (LD) regions (*r*^2^ > 0.1), 121 (21%) were new loci, not previously associated with any of the seven externalizing behaviors/disorders that went into the Genomic SEM model, and 41 (7%) can be classified as entirely novel, as they have not been reported previously for any trait in the GWAS literature.

Genomic SEM was used to perform SNP-level tests of heterogeneity (*Q*_SNP_; Supplementary Information section 3.5.1) that investigate whether each SNP had consistent, pleiotropic effects on the seven input phenotypes that effectively operate via the shared genetic liability *EXT* (**Extended Data Fig. 2**). Only 1% (8/579) of the 579 *EXT* SNPs were significant (one-sided *Q*_SNP_ *P* < 5×10^-8^) in *Q*_SNP_ tests (**Figure 1D; Supplementary Table 9**), providing further evidence that the genetic variants we identified primarily index a unitary dimension of genetic externalizing liability rather than representing an amalgamation of variants with divergent associations across the discovery phenotypes. The genome-wide *Q*_SNP_ analysis was adequately powered (mean χ^2^ = 1.864; **Extended Data Fig. 2**), and as expected, it identified heterogeneity in regions of the genome not associated with *EXT*. The strongest *Q*_SNP_ and most salient example of a trait-specific association is SNP rs1229984 (one-sided *Q*_SNP_ *P* = 1.67×10^-51^). This particular SNP, located in the gene *ADH1B*, is a known missense variant with a well-established role in alcohol metabolism^23^, and it was not associated with *EXT* (two-sided *P* = 0.022) but only with problematic alcohol use (two-sided *P* = 6.43 × 10^-57^).

Because the discovery stage effectively exhausted large study cohorts available for strict replication, we instead performed a series of preregistered quasi-replication analyses, which have previously been applied successfully in the GWAS setting^24,25^. Further below, we additionally perform holistic quasi-replication of the 579 *EXT* SNPs in polygenic score analyses (also in within-family models). For SNP-level quasi-replication analyses of the 579 SNPs (Supplementary Information section 4), a three-step holistic method tested their association with two independent, GWAS meta-analyses on externalizing phenotypes: (1) alcohol use disorder (*r_g_* with *EXT* = 0.52; *N* = 202,004), and (2) antisocial behavior (*r_g_* with *EXT* = 0.69; *N* = 32,574). First, we tested whether the 579 SNPs (or an LD proxy for missing SNPs, *r^2^* > 0.8) showed sign concordance, *i.e.*, the same direction of effect between *EXT* and alcohol use disorder or antisocial behavior: 75.4% of SNPs showed sign concordance with alcohol use disorder (two-sided test *P* = 6.84×10^-36^) and 66.9% with antisocial behavior (two-sided test *P* = 1.39×10^-15^) (**Extended Data Fig. 3**). For the second and third tests, we generated empirical null distributions for the two phenotypes by randomly selecting 250 near-independent (*r^2^* < 0.1) SNPs per each of the 579 SNPs, matched on allele frequency. In the second test, a greater proportion of the 579 SNPs were nominally associated (*P* < 0.05) with the two phenotypes compared to their empirical null distributions: 124 (21.4% vs. 6.6%) with alcohol use disorder (two-sided *P* = 1.87×10^-31^) and 58 (10.5% vs. 4.7%) with antisocial behavior (*P* = 1.64×10^-8^). In the third test, the 579 SNPs were jointly more strongly enriched for association with alcohol use disorder (one-sided Mann-Whitney test *P* = 5.89×10^-26^) and antisocial behavior (*P* = 1.10×10^-5^) compared to their empirical null distributions. Overall, the quasi-replications consistently suggested that the GWAS of *EXT* is not spurious overall, and that it is enriched for genetic signal with phenotypes of central importance to the literature on externalizing.

### Bioinformatic analyses highlight relevant neurodevelopmental and biological processes

We performed a series of bioinformatic analyses to explore the biological processes underlying externalizing liability (Supplementary Information section 6, **Supplementary Tables 9–10, and 21–29; Extended Data Figs. 5–8**). Consistent with the idea that persistent difficulties in self-regulation can be conceptualized as a neurodevelopmental condition^26,27^, MAGMA gene-property analyses suggested an abundance of enrichment in genes expressed in brain tissues, particularly during prenatal developmental stages (**Extended Data Fig. 7**), with the strongest enrichment seen in the cerebellum, followed by frontal cortex, limbic system tissues, and pituitary gland tissues (**Extended Data Fig. 6**). Furthermore, MAGMA gene-set analysis identified gene sets related to neurogenesis, nervous system development, and synaptic plasticity, among other gene-sets related to neuronal function and structure.

Because of the strong polygenic signal identified in the GWAS of *EXT*, four different gene-based analyses identified an abundance of implicated genes (>3,000): (1) functional annotation of the 579 SNPs to their nearest gene with FUMA^28^, which suggested 587 genes; (2) MAGMA gene-based association analysis^29^, which identified 928 Bonferroni-significant genes (one-sided test *P* < 2.74×10^-6^); (3) H-MAGMA^30^, a method that assigns non-coding SNPs to cognate genes based on chromatin interactions in adult brain tissue and which identified 2,033 Bonferroni-significant genes (one-sided test *P* < 9.84×10^-7^); and (4) S-PrediXcan^31^, which uses transcriptome-based analyses of predicted gene expression in 13 brain tissues and which identified 348 Bonferroni-significant gene-tissue pairs (two-sided test *P* < 2.73×10^-7^).

We found 34 genes that were consistently identified in all four methods, while 741 overlapped across two or more methods (**Supplementary Table 29; Extended Data Fig. 8**). Several of the 34 implicated genes are novel discoveries for the psychiatric/behavioral literature and have previously been identified only in relation to biomedical disease. Such discoveries include *ALMS1* (previously associated with kidney function and urinary metabolites^32^), and *ERAP2* (blood protein levels and autoimmune disease ^33,34^). Other genes among the 34 have previously been identified in GWAS of behavioral or psychiatric traits: Cell Adhesion Molecule 2 (*CADM2*, previously identified in GWAS related to self-regulation, including drug use and risk tolerance^16,35^), Zic Family Member 4 (*ZIC4*, associated with brain volume^36^), Gamma-Aminobutyric Acid Type A Receptor Subunit Alpha 2 (*GABRA2*; the site of action for alcohol and benzodiazepines, extensively studied in relation to alcohol dependence^37,38^, and proposed candidate gene for many psychiatric disorders^39,40^), *NEGR1* (neuronal growth regulator, associated with intelligence and educational attainment^25,41^), and Paired Basic Amino Acid Cleaving Enzyme (*FURIN*, associated with schizophrenia, risk tolerance, and trans-diagnostic vulnerability to psychiatric disorders^42,43^).

### Genetic risk scores explain substantial variation in behavioral, psychiatric, and social outcomes

We created a genome-wide polygenic score for *EXT*, adjusted for LD^44,45^, among subjects from two European-ancestry datasets selected for their detailed phenotypes related to externalizing outcomes (Supplementary Information section 5): (1) the National Longitudinal Study of Adolescent to Adult Health (Add Health; *N* = 5,107), a U.S.-based study of adolescents who were recruited from secondary schools in the mid-1990s; (2) the Collaborative Study on the Genetics of Alcoholism (COGA; *N* = 7,594), a U.S.-based study focused on understanding genetic contributions to alcohol use disorders.

To investigate the validity of *EXT*, in each sample, we fit a latent factor model to phenotypic data corresponding to the seven Genomic SEM phenotypes (**Extended Data Fig. 4** and **Supplementary Table 13**). Controlling for age, sex, and ten principal components of genetic ancestry, the *EXT* polygenic score was strongly associated with the latent phenotypic factor in both data sets (*β*_Add Health_ = 0.33, 95% CI, 0.30 to 0.36, Δ*R*^2^ = 10.5%; *β*_COGA_ = 0.30, 95% CI, 0.27 to 0.34, Δ*R*^2^ = 8.9%; **Figure 2A** and **Supplementary Table 14**). The variance explained by the *EXT* polygenic score (Δ*R*^2^ ~ 8.9-10.5%) is commensurate with many conventional variables used in social science research, including parental socioeconomic status, family income or structure, and neighborhood disadvantage/disorder^46–48^. Next, as further quasi-replication, in each sample we created a polygenic score using only the 579 *EXT* SNPs. This polygenic score was associated with the latent phenotypic externalizing factor in both samples, explaining ~3–4% of the variance (*β*_Add Health_ = 0.20, 95% CI, 0.17 to 0.23; *β*_COGA_ = 0.17, 95% CI, 0.13 to 0.20; **Supplementary Table 14**).

**Figure 2.**
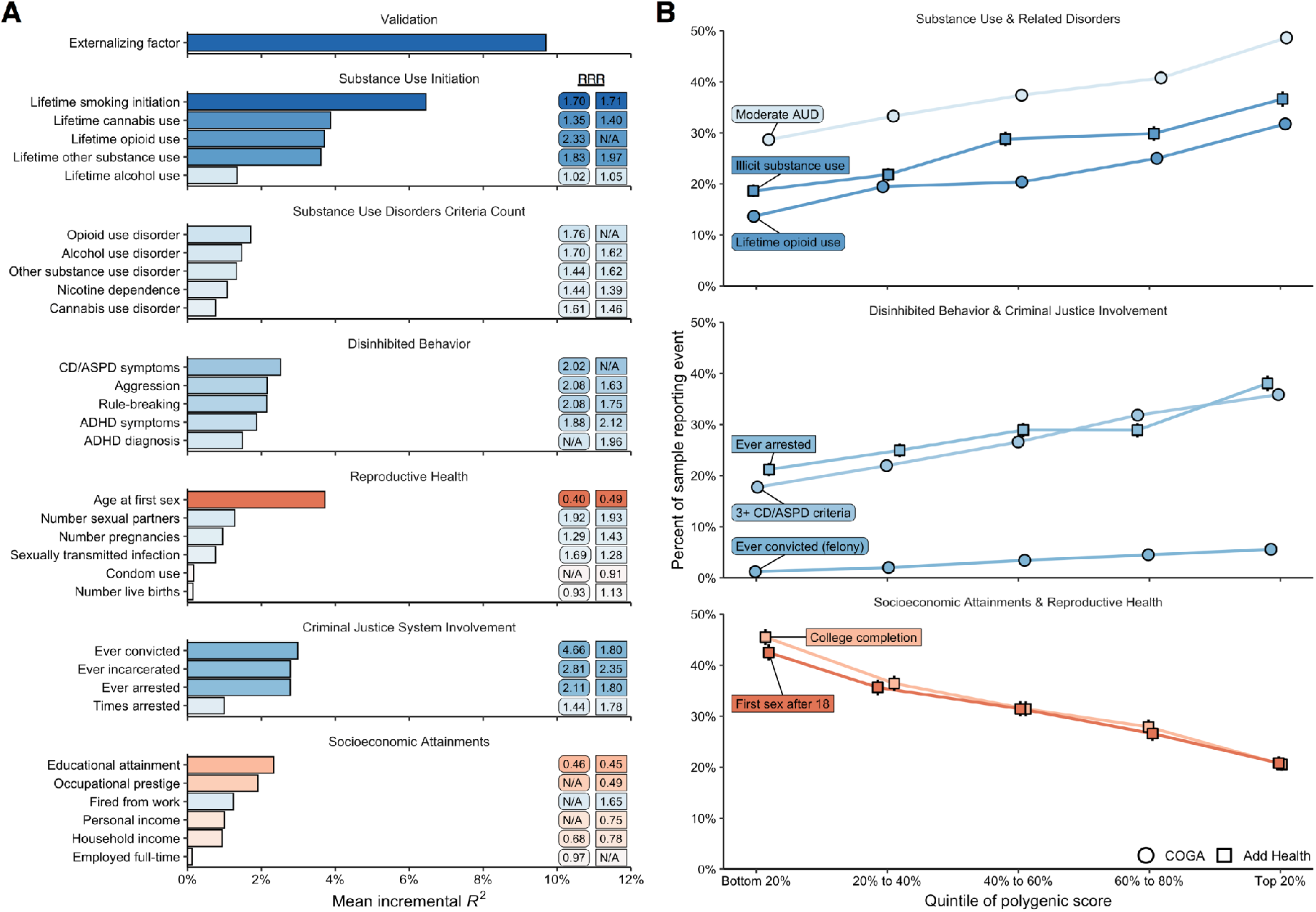
Polygenic score associations with behavioral, psychiatric, and social outcomes in the independent Add Health (*N* = 5,107) and COGA (*N* = 7,594) datasets. (**A**) Bar charts illustrating the mean proportion of variance (incremental *R*^2^, or Δ*R*^2^) explained by the polygenic score. Blue and red bars indicate positive and negative associations, respectively. Relative risk ratios (RRRs), comparing individuals in the lowest 20% to those in the highest 20% of the polygenic score distribution, are reported for Add Health and COGA in square and round boxes, respectively. (**B**) Line charts illustrating the relative risks across quintiles of the polygenic score for eight illustrative outcomes: (1) meeting 4 or more criteria for alcohol use disorder (AUD), (2) lifetime use of an illicit substance other than cannabis, (3) lifetime opioid use, (4) ever being arrested, (5) meeting 3 or more criteria for conduct disorder (CD) or antisocial personality disorder (ASPD), (6) ever being convicted of a felony, (7) completing college, and (8) first sexual intercourse at the age of 18 or older. 95% confidence intervals are presented with error bars for each quintile.

We next explored to what extent polygenic scores for *EXT* were associated with childhood externalizing disorders and a variety of specific phenotypes that reflect difficulty with self-regulation or its social consequences (**Figure 2B** and **Supplementary Tables 16– 19**). Polygenic scores for *EXT* explained significant variance (Δ*R*^2^) in criteria counts of ADHD (mean Δ*R*^2^ = 1.65%), conduct disorder (CD; mean Δ*R*^2^ = 3.1%), and oppositional defiant disorder (ODD; Δ*R*^2^ = 1.96%), as well as in phenotypes categorized as substance use initiation (mean Δ*R*^2^ = 1.3–6.5%), substance use disorders (mean Δ*R*^2^ = 0.8–1.7%), disinhibited behaviors (mean Δ*R*^2^ = 1.5–2.5%), criminal justice system involvement (mean Δ*R*^2^ = 1.0–3.0%), reproductive health (mean Δ*R*^2^ = 0.3–3.7%), and socioeconomic attainment (mean Δ*R*^2^ = 0.1–2.3%). Many of the phenotypes – such as opioid use disorder criteria count, conduct disorder and antisocial personality disorder criteria count, lifetime history of arrest or incarceration, and lifetime history of being fired from work, were not included in our Genomic SEM analyses; however, our *EXT* polygenic score is notable in capturing appreciable variance in phenotypes that are still lacking large GWAS samples (a striking example being opioid use disorder^8^). The associations between the *EXT* polygenic score and this broad range of phenotypes represents an affirmative test of the hypothesis that genetic variants associated with externalizing liability generalize to a wide variety of behavioral and social outcomes related to behavioral undercontrol.

To evaluate medical outcomes associated with genetic liability to externalizin*g*, we conducted a phenome-wide association study (PheWAS) in 66,915 genotyped individuals of European-ancestry in the BioVU biorepository, a U.S.-based biobank of electronic health records from the Vanderbilt University Medical Center, spanning 1990 to 2017^49,50^. A logistic regression was fit to 1,335 case/control disease phenotypes. Of these, 255 disease phenotypes were associated with the *EXT* polygenic score at a false discovery rate less than 0.05 (**Figure 3** and **Supplementary Table 20**). The most abundant associations were with mental and behavioral disorders, such as substance use, mood disorders, suicidal ideation, and attempted suicide. Individuals with higher *EXT* polygenic scores also showed worse health in nearly every bodily system. They were more likely to suffer, for example, from ischemic heart disease, viral hepatitis C and HIV infection, type 2 diabetes and obesity, cirrhosis of liver, sepsis, and lung cancer. Notably, many of these medical outcomes are mediated by behaviors related to self-regulation, e.g., smoking, drinking, drug use, condomless sex, and overeating.

**Figure 3.**
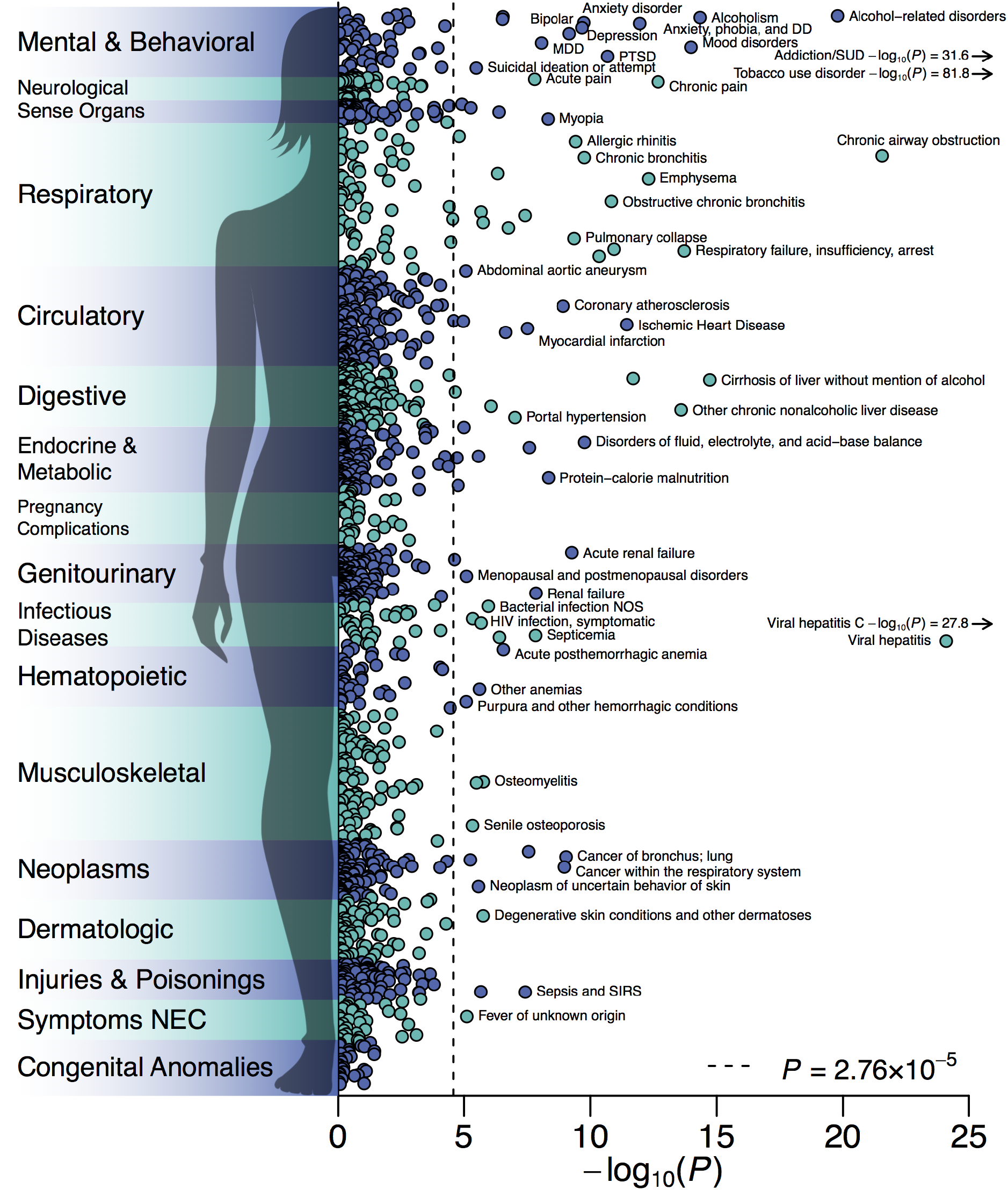
Phenome-wide association study in the BioVU biorepository. −log_10_ *P* values of two-sided test for association of polygenic score for *EXT* with 1,335 medical outcomes were derived with logistic regression in up to 66,915 patients, adjusted for sex, median age in the EHR data, and the first 10 genetic PCs. The dashed line is the Bonferroni-corrected significance threshold; adjusted for the number of tested medical conditions. 84 medical conditions were Bonferroni-significant, while 255 conditions were significant at a false discovery rate less than 0.05. The labels for some conditions were omitted. The full results, including case-control counts, effect sizes, and standard errors, are reported in **Supplementary Table 20**.

### Within-family analyses demonstrate that polygenic associations are robust to confounding

Genetic associations detected in GWAS can be due to direct genetic effects, but can also be confounded by uncontrolled population stratification, indirect genetic effects mediated through the parental environment, and assortative mating^51,52^. While reducing statistical power, sibling comparisons overcome these methodological challenges, because meiosis randomizes genotypes to siblings^51,53^. We therefore conducted within-family analyses of polygenic score associations in the sibling sub-samples of Add Health (*N* = 994 siblings from 492 families) and COGA (*N* = 1,353 siblings from 621 families), as well as a sample of sibling pairs from the UK Biobank (*N* = 39,640), which were held-out from the discovery stage (Supplementary Information section 2.3.2).

In Add Health and COGA, the phenotypic factor derived from observations corresponding to the seven discovery phenotypes (see above) was regressed on the *EXT* polygenic scores in a within-family model (**Supplementary Table 15**). Parameter estimates from the within-family models (β_Add Health_ = 0.12, 95% CI, 0.04 to 0.20; β_COGA_ = 0.14, 95% CI, 0.08 to 0.20) were slightly attenuated compared to OLS models without family-specific intercepts (β_Add Health_ = 0.20, 95% CI, 0.16 to 0.24; β_COGA_ = 0.16, 95% CI, 0.12 to 0.20), but remained strong (Add Health β / β_WF_ = 1.667; COGA β / β_WF_ = 1.142) and statistically significant (two-sided test *P* = 4.89×10^-3^ and 1.87×10^-6^, respectively). Additionally, the association of the quasi-replication polygenic score constructed with the 579 *EXT* SNPs did not attenuate in within-family models and remained significant (**Supplementary Table 15**).

In the UK Biobank sibling hold-out sample, we conducted polygenic score analyses of 33 phenotypes from the domains of risky behavior, reproductive health, cognitive ability, personality, and socioeconomic status (**Supplementary Table 19**). Similar to Add Health and COGA, within-family estimates were only modestly attenuated for risky behavior and reproductive health outcomes (mean β / β_WF_ = 1.079); however, effect-sizes in within-family models were substantially attenuated for cognitive ability and socioeconomic status outcomes (β / β_WF_ was 3.3 for educational attainment, 4.9 for household income, 2.1 for neighborhood deprivation). Overall, the *EXT* polygenic score remained significantly associated (two-sided test *P* < 0.05) with 21 outcomes, showing that our GWAS of externalizing captures direct genetic effects on behavioral health and is not solely a consequence of uncontrolled population stratification, indirect genetic effects, or other forms of environmental confounding.

## Discussion

Externalizing disorders and behaviors are a widely prevalent cause of human suffering, but understanding of the molecular genetic underpinnings of externalizing has lagged considerably behind progress made in other areas of medical and psychiatric genetics. For example, dozens of associated genetic loci have been discovered for schizophrenia (>100 loci)^54^, bipolar disorder (30 loci)^55^, and major depressive disorders (44 loci)^56^, whereas recent GWASs of antisocial behavior^57^, alcohol use disorders^58^, and opioid use disorders^8^ have identified only a very small number of significantly associated loci, if any at all. Here, we used multivariate genomic analyses to accelerate genetic discovery, identifying 579 genome-wide significant loci associated with a predisposition toward externalizing disorders and behaviors, 121 of which are entirely novel discoveries for any of the seven phenotypes analyzed. Our results demonstrate that moving beyond traditional disease classification categories can enhance gene discovery, improve polygenic scores, and provide information about the underlying pathways by which genetic variants impact clinical outcomes. GWAS efforts find almost ubiquitous genetic correlations across psychiatric disorders and diagnoses^59,60^; new analytic methods now allow us to capitalize on these genetic correlations. Pragmatically, non-disease phenotypes such as the ones we use here (*e.g.*, self-reported age at first sex) are often easier to measure in the general population than diagnostic status, making it easier to achieve large sample sizes. Expanding beyond individual diagnoses increases our ability to detect genes underlying human behavioral and medical outcomes of consequence.

Our results highlight again that there is no distinct line between the genetic study of biomedical conditions and the genetic study of social and behavioral traits^61^. Linking biology with socially-valued behavioral outcomes can be politically sensitive (**Box 1**)^62^. Polygenic scores created using our GWAS results were associated not just with psychiatric and substance use disorders, but also with correlated social outcomes, such as lower employment and greater criminal justice system involvement, as well as with biomedical conditions affecting nearly every system in the body. Considered together, our analyses demonstrate the far-reaching toll of human suffering borne by people with high genetic liabilities to externalizing.

### Box 1. Grappling with the Legacy of Eugenics

In 1912, Henry Goddard published what is now considered an infamous work of pseudoscience: *The Kallakak Family* traced several generations of a “feeble-minded” family to argue that not just intellectual ability, but also drunkenness, criminality, sexual promiscuity, and morality were hereditary ^63^. On the basis of these pedigrees, Goddard recommended that the “feeble-minded” should be institutionalized and prohibited from reproducing. Horrifically, these recommendations were put into practice: Involuntary sterilization programs and other forms of state-sponsored violence targeting the poor and ethnic/racial minorities persisted for decades^64,65^. Even now, the danger of eugenics is not safely in the past. Modern genetics research is routinely appropriated by white supremacist movements to argue that racialized disparities in health, employment, and criminal justice system involvement are due to the genetic inferiority of people of color rather than environmental and historical disadvantages^66–68^. At the same time, failing to understand how genetic differences contribute to vulnerability to externalizing can increase stigma and blame for these behaviors^69,70^. Given the horrific legacy of eugenics, the ongoing reality of racism in the medical and criminal justice systems, and the importance of combatting stigma in psychiatric disorders, the scientific results we report here, which are, for technical reasons, limited to European individuals, must be interpreted with the utmost care. Please see our supporting materials at www.externalizing.org for more information.

Our polygenic score for externalizing has one of the largest effect sizes of any polygenic score in psychiatric and behavioral genetics, accounting for 10% of the variance in externalizing factor scores, and meaningful variance in outcomes as varied as opioid use, age at first sex, being fired from work, and being convicted of a crime. These effect sizes rival the associations observed with “traditional” covariates used in social science research. But, these effect sizes remain far below twin estimates of heritability for externalizing^5^ and far below what is necessary to predict these outcomes for any individual^71,72^. Furthermore, while effect sizes were only modestly attenuated in within-family models of risky behavior and reproductive behavior, they were substantially attenuated in analyses of socioeconomic outcomes, indicating that substantial work remains to be done to clarify the association between externalizing genetics and socioeconomic inequality^51^. Additionally, application of these genetic discoveries to improve research and intervention will be limited as long as the samples available for genomics research fail to reflect the world’s genetic diversity^73^.

Finally, these results are *not* evidence that some people are genetically determined to experience certain life outcomes or are “innately” antisocial. Genetic differences are probabilistically associated with psychiatric, medical, and social outcomes, in part via environmental mechanisms that might differ across historical, political, and economic contexts^74^. For example, a policy change like decriminalization of cannabis use might mitigate associations between genetic vulnerabilities and criminal justice system involvement, because the state ceases to criminalize a behavior to which some individuals have a greater genetic susceptibility. At the same time, increased availability and decreased stigma may create environments more conducive to the development of substance problems among individuals who are genetically at risk^75^. The impact of genetic factors might also depend on other forms of social capital and privilege. For instance, childhood externalizing is associated with greater adult earnings, but only for children not raised in poverty^76,77^. The genetic differences identified here can thus be used in future research as a tool to trace how lifespan development is shaped via complex interactions between genetic predispositions, environmental influences (*e.g.*, parenting, peer, and romantic relationships) and social institutions (*e.g.*, schools, jails, hospitals, creditors, and employers).

## Online methods

The article is accompanied by Supplementary Information with further details. The study was performed according to a preregistered analysis plan (https://doi.org/10.17605/OSF.IO/XKV36), which specified that we would either generate new or collect existing single-phenotype genome-wide association study (GWAS) summary statistics on phenotypes related to the externalizing spectrum (Supplementary Information section 1). In the discovery stage, the summary statistics were to be analyzed with Genomic SEM with the aims of (a) estimating a genetic factor structure underlying externalizing liability, (b) identifying single-nucleotide polymorphisms (SNPs) and genes primarily involved in a shared genetic liability to externalizing, and (c) increasing the accuracy of genetic risk scores for specific externalizing phenotypes that are currently intractable to study in large samples. To ensure satisfying statistical power, we preregistered a minimum sample-size threshold of *N* > 15,000, and that additional exclusions would be based on displaying negligible or inaccurate SNP-based heritability or genetic covariance. The study did not manipulate an experimental condition, and thus, was neither randomized nor blinded.

### Collecting existing single-phenotype GWAS on externalizing phenotypes

A detailed definition of “externalizing phenotypes” was preregistered to delimit the data collection of single-phenotype GWAS summary statistics (Supplementary Information section 2.1). Summary statistics from existing studies were either provided by or downloaded from the public repositories of 23andMe, the Psychiatric Genomics Consortium (PGC), the Million Veterans Program (MVP), the International Cannabis Consortium (ICC), the GWAS & Sequencing Consortium of Alcohol and Nicotine Use (GSCAN), the Social Science Genetics Association Consortium (SSGAC), the Genetics of Personality Consortium (GPC), and the Broad Antisocial Behavior Consortium (Broad ABC), see Supplementary Information section 2.2 for more details. All GWAS that were considered for inclusion are listed in **Supplementary Table 1**, and **Supplementary Table 2** reports the underlying studies that had contributed to the seven GWAS (or GWAS meta-analysis) that were included the final multivariate model specification (see below).

### GWAS in UK Biobank (UKB)

New GWAS were estimated in UKB (Supplementary Information section 2.3), of which summary statistics for “age at first sexual intercourse” and “Alcohol Use Disorder Identification Test problem items” (AUDIT-P) were later included in the final multivariate model (see below). The GWAS were performed with linear mixed models (BOLT-LMM^78^) and were statistically adjusted for sex, birth year, sex-specific birth-year interaction dummies, genotyping array and batch, and 40 genetic principal components (PCs). Two partly overlapping hold-out subsamples of UKB participants were excluded from all single-phenotype GWAS summary statistics that included UKB data, and the participants were instead retained as an independent sample for polygenic score analyses (Supplementary Information section 2.3.2). Genetic relatives (pairwise KING coefficient ≥ 0.0442) of the held-out individuals were excluded from the study altogether to ensure independence between the discovery and follow-up analyses. Whenever an existing GWAS (or meta-analysis) was based on UKB data, we re-estimated the UKB component using the same phenotype definition as in the existing study, while excluding the held-out participants and their genetic relatives. See Supplementary Information section 2.3.2 for further details.

### GWAS inclusion criteria, quality control, and meta-analysis

All GWAS were performed among individuals that (a) were of European ancestry, (b) were observed for all relevant covariates, (c) were successfully genotyped and passed standardized sample-level quality control (according to study-specific protocols^12–15,21,79^), and (d) were unrelated (unless a particular GWAS was estimated with linear mixed models). Genotypes were imputed with reference data from either the 1000 Genomes Consortium^80^, the Haplotype Reference Consortium^81^, the UK10K Consortium^82^, or a combination thereof. We performed quality control of GWAS summary statistics with a whole-genome sequenced reference panel, assembled from 1000 Genomes Consortium^80^ and UK10K Consortium^82^ data (Supplementary Information section 2.4.1). Our quality-control procedure applied recommended^83^ SNP-filtering to remove rare SNPs (minor allele frequency < 0.005), SNPs with an IMPUTE imputation quality (INFO) score less than 0.9, and otherwise low-quality variants (**Supplementary Table 3**). For a complete description of the quality-control procedure, see Supplementary Information section 2.4.

We performed sample-size weighted meta-analysis with METAL^84^ (Supplementary Information section 2.5). Thereafter, we excluded any summary statistics that displayed insufficient SNP-based heritability (*h*^2^ < 0.05) or GWAS association signal 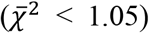, estimated with LD Score regression^18,59^. At this stage, we had collected or generated well-powered summary statistics for eleven phenotype-specific GWAS (or meta-analysis) that satisfied our inclusion criteria and that were kept for exploratory factor analysis (**Supplementary Table 4**): (1) ADHD (*N* = 53,293), (2) age at first sexual intercourse (*N* = 357,187), (3) problematic alcohol use (*N* = 164,684), (4) automobile speeding propensity (*N* = 367,151), (5) alcoholic consumption (drinks per week; *N* = 375,768), (6) educational attainment (*N* = 725,186), (7) lifetime cannabis use (*N* = 186,875), (9) lifetime smoking initiation (*N* = 1,251,809), (9) general risk tolerance (*N* = 426,379), (10) irritability (*N* = 388,248), and (11) number of sexual partners (*N* = 336,121).

### Exploratory factor analysis of genetic correlations

As an initial analysis to inform and guide the multivariate modeling process, we performed hierarchical clustering of a matrix with pair-wise LD Score genetic correlations (*r_g_*) (Supplementary Information section 3). The GWAS effect-sizes of age at first sexual intercourse and educational attainment were reversed to anticipate positive correlations with externalizing liability. The 11 phenotypes displayed moderate-to-substantial genetic overlap with at least one other phenotype (max |*r_g_*| = 0.245–0.773), and the average |*r_g_*| across all pairwise correlations was 0.323 (**Supplementary Table 5**). Three clusters were identified: (1) attention deficit/hyperactivity disorder (ADHD), educational attainment (EDUC), age at first sexual intercourse (FSEX), irritability (IRRT), and smoking initiation (SMOK); (2) problematic alcohol use (ALCP), drinks per week (DRIN); and (3) lifetime cannabis use (CANN), automobile speeding propensity (DRIV), number of sexual partners (NSEX), general risk tolerance (RISK).

Following the preregistration, exploratory factor analysis tested four different factor solutions, specifying 1…*k* + 1 factors (Supplementary Information section 3.2), where *k* corresponds to the number of clusters identified in the genetic correlation matrix, while retaining factors that explained at least 15% of the variance (a preregistered threshold). Exploratory factor analysis found that the fourth factor explained only 12.5% of the variance, and thus, the three-factor solution was considered the most appropriate exploratory model in terms of capturing variation (**Supplementary Table 6**). The pattern of factor loadings was consistent with the hierarchical clustering. However, as we detail in Supplementary Information section 3.2, the second and third factor mainly accounted for complex residual variation and divergent residual cross-trait correlations among the subset of phenotypes that had the weakest loadings on the single common factor. Thus, we learned from the exploratory factor analysis that some of the 11 indicators may not be optimal for identifying a single common genetic liability to externalizing, and that a less complex model specification with fewer indicators would perhaps perform better than a three-factor model in the subsequent confirmatory factor analysis.

### Confirmatory factor analyses with Genomic SEM

We formally modelled genetic covariances (rather than genetic correlations) and performed confirmatory factor analyses using the method genomic structural equation modeling (Genomic SEM)^10^ (Supplementary Information section 3.3). Genomic SEM is unbiased by sample overlap and differences in sample size in the discovery phenotypes, and by applying to GWAS summary statistics it allows for genetic analyses of latent factors in larger samples than is typically possible with individual-level data^10^. We competed four models: (1) a common factor model with the aforementioned 11 phenotypes, (2) a correlated three-factors model with the 11 phenotypes (with and without cross-loadings), (3) a bifactor model with the 11 phenotypes, and (4) a revised common factor model that only included seven of the phenotypes that satisfied moderate-to-large (*i.e.*, ≥ .50) loadings on the single latent factor in model (1) (**Supplementary Table 7**). We found that model (4) was the only model that closely approximated the observed genetic covariance matrix (*χ*^2^(12) = 390.234, AIC = 422.234, CFI = .957, SRMR = .079), fulfilled our preregistered model fit criteria, and coalesced with theoretical expectations of a general shared genetic liability to externalizing. This model was selected as our final factor specification, and we hereafter refer to it as “the latent genetic externalizing factor”, or simply, “the externalizing factor” (*EXT*). To explore the convergent and discriminant validity of the externalizing factor, we estimated genetic correlations between the externalizing factor and 92 traits from various research domains (**Supplementary Table 8**).

### Multivariate GWAS analyses with Genomic SEM

Using Genomic SEM, we performed multivariate GWAS analysis by estimating SNP associations with the externalizing factor (*EXT*), which is our main discovery analysis (Supplementary Information section 3.4). We estimated the effective sample size of the resulting “externalizing GWAS” to be *N*_eff_ = 1,492,085. The GWAS displayed strong association signal, with a mean χ^2^ and genomic inflation factor (λ_GC_) of 3.114 and 2.337, respectively. Analyses with LD Score regression suggest that the strong inflation observed in the association test statistic is attributable to polygenicity rather than bias from population stratification^10,18^, as the LD Score intercept and attenuation ratio were estimated to be 1.115 (*SE* = 0.019) and 0.054 (*SE* = 0.009), respectively.

A conventional “clumping” algorithm was applied to identify near-independent genome-wide significant lead SNPs (two-sided *P* < 5×10^-8^)^85^, which were then subjected to “multi-SNP-based conditional & joint association analysis using GWAS summary data” (COJO) to estimate conditional SNP associations^22,86^ (Supplementary Information section 3.4.2). We identified 579 lead SNPs that were conditionally and jointly associated with *EXT*. We performed lookups of these “579 *EXT* SNPs”, as well as any correlated SNPs (*r^2^* > 0.1), in the NHGRI-EBI GWAS Catalog^7^ (version e96 2019-05-03) to investigate whether the identified loci have previously been found associated with other traits at suggestive significance (two-sided *P* < 1×10^-5^). To evaluate whether each SNP acted through the externalizing factor, we estimated genome-wide *Q*_SNP_ heterogeneity statistics with Genomic SEM (Supplementary Information section 3.5.1). The null hypothesis of the *Q*_SNP_ test is that SNP effects on the constituent phenotypes operate (i.e., are statistically mediated) via the *EXT* factor, so a significant *Q*_SNP_ test indicates that SNP association is better explained by a trait-specific pathway independent of the *EXT* factor. The *Q*_SNP_ analysis was sufficiently powered to identify substantial heterogeneity in the genome (160 near-independent genome-wide significant *Q*_SNP_ loci), but reassuringly, did not identify heterogeneity among 99% (571/579) of the *EXT* SNPs. **Supplementary Table 9–10** reports the results of the externalizing GWAS and the heterogeneity analysis, together with bioannotation with “functional mapping and annotation of genetic associations” (FUMA)^28^.

### Proxy-phenotype and quasi-replication analysis

We performed a preregistered proxy-phenotype^87^ and quasi-replication^24^ analysis by investigating the 579 SNPs (*k*) for association in two independent, second-stage GWAS on (1) alcohol use disorder (*N* = 202,004, *r_g_* = 0.52) and (2) antisocial behavior (*N* = 32,574, *r_g_* = 0.69) (Supplementary Information section 4). For SNPs missing from the two second-stage GWAS, we analyzed highly correlated proxy SNPs (*r^2^* > 0.8). Significant proxy-phenotype associations were evaluated for Bonferroni-corrected significance (two-sided test *P* < 0.05/*k*). For the quasi-replication exercises, we generated empirical null distributions for the two second-stage GWAS by randomly selecting 250 near-independent (*r^2^* < 0.1) SNPs matched on MAF (± 1 percentage point) for each of the *k* SNPs. The quasi-replication approach was performed in three steps: (1) a binomial test of sign concordance, which tested whether the direction of effect of the *k* SNPs were in greater concordance between the externalizing GWAS and each of the second-stage GWAS compared to what would be expected by chance (H0 = 0.5); (2) a binomial test of whether a greater proportion of the *k* SNPs were nominally significant (two-sided *P* < 0.05) in the second-stage GWAS compared to the empirical null distribution; (3) a test of joint enrichment, performed as a non-parametric (one-sided) Mann-Whitney test of the null hypothesis that the *P* values of the *k* SNPs are derived from the empirical null distribution. We strongly rejected the null hypotheses of all quasi-replication tests, suggesting that the externalizing GWAS is not spurious overall and that it was more enriched for association with the second-stage phenotypes than their respective polygenic background GWAS signal (**Supplementary Table 11–12**).

### Polygenic score analyses

We generated polygenic scores by summing genotypes weighed by the effect sizes estimated in the externalizing GWAS, among individuals of European ancestry in five independent study cohorts: (1) Add Health^88,89^, (2) COGA^90–92^, (3) PNC^93,94^, (4) the UKB siblings hold-out cohort^95^, and (5) the BioVU biorepository^96^ (Supplementary Information section 5). In each dataset, we generated three scores, of which two were adjusted for linkage disequilibrium (LD): (1) PRS-CS^45^, (2) LDpred (infinitesimal model)^44^, and (3) unadjusted scores^97^, while using SNPs that overlapped with the high-quality consensus set defined by the HapMap 3 Consortium^98^. Accuracy was evaluated as the incremental *R^2^*/pseudo-*R^2^* (Δ*R*^2^) attained by adding the polygenic score to a regression model with baseline covariates, in accordance with previous efforts^16,99^. The baseline model included covariates for sex, age, and genetic principal components (PCs), and genotyping array and batch. The choice of statistical model (e.g., OLS vs. logit) and adjustment of standard errors depended on (1) the distribution of the phenotype and (2) the structure of the data in the study cohort (independent vs. clustered observations), see Supplementary Information section 5.2.4 for further details. We estimated 95% confidence intervals for Δ*R*^2^ using percentile method bootstrapping with 1000 iterations.

In Add Health and COGA, we performed out-of-sample validation of *EXT* by modeling a latent externalizing factor using phenotypic data corresponding to the seven Genomic SEM phenotypes (Supplementary Information section 5.2.3) (**Supplementary Table 13–14**). In Add Health, COGA, PNC, and the UKB siblings hold-out cohort, we performed exploratory polygenic score analyses with a wide range of preregistered phenotypes from the behavioral, psychiatric, and socioeconomic research domains (**Supplementary Table 16–19**). We performed a phenome-wide association study (PheWAS) of medical outcomes in the BioVU biorepository by fitting a logistic regression to 1,335 case/control disease “phecodes”^100^ (*N* = 66,915) (**Supplementary Table 20**).

We performed within-family analyses in data on full siblings in Add Health, COGA, and the UKB siblings hold-out cohort (Supplementary Information section 5.2.5). We analyzed 492 families in Add Health (*N*_siblings_ = 994), 621 families in COGA (*N*_siblings_ = 1,353), and 19,252 families in the UKB (*N*_siblings_ = 39,640). In Add Health and COGA, we applied OLS to test the externalizing polygenic score for association with a single outcome: the factor scores of the phenotypic externalizing factor (a continuous variable), while adjusting for family fixed-effects (i.e., family-specific dummy variables) (**Supplementary Table 15**). We then compared the magnitude of the within-family coefficient 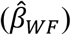 to the coefficient of an OLS model without family-specific intercepts 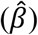. In the UKB siblings hold-out cohort, we performed an analogous within-family analysis of the exploratory phenotypes (**Supplementary Table 19**). We analyzed heteroskedasticity-consistent and cluster-robust standard errors, clustered at the family level.

### Bioannotation

We performed a series of bioannotation and bioinformatic analyses to identify relevant biological pathways (Supplementary Information section 6). The method “functional mapping and annotation of genetic associations” (FUMA v1.3.5e)^28^ was applied to explore the functional consequences of the 579 SNPs (**Supplementary Table 9**), which included ANNOVAR categories (*i.e.*, the functional consequence of SNPs on genes), Combined Annotation Dependent Depletion (CADD) scores (*i.e.*, a measure of how deleterious a SNP is; CADD > 12.37 is classified as deleterious), RegulomeDB scores (*i.e.*, a categorical score from 1a to 7 with 1a corresponding to the most biological evidence that the SNP is a regulatory element), mapping to expression quantitative trait loci (eQTLs), and chromatin states (values range from 1 to 15, with values 1 to 7 referring to an open chromatin state). The sources of the external reference data used by FUMA are described in ref.^28^.

Gene-based association analyses was performed by applying the method “multi-marker analysis of genomic annotation” (MAGMA v1.07)^28,29^ (Supplementary Information sections 6.1.2). The method accounts for LD, which was calculated using reference data from European-ancestry 1000 Genomes participants^80^. Genome-wide SNPs were first mapped to 18,093 protein-coding genes from Ensembl (build 85)^101^, and the SNPs within each gene were then jointly tested for association with *EXT*. We evaluated Bonferroni-corrected significance, adjusted for the number of tested genes (one-sided *P* < 2.76×10^-6^) (**Supplementary Table 21**). Next, MAGMA gene-set analysis was performed using 15,477 curated gene sets and Gene Ontology (GO)^102^ terms obtained from the Molecular Signatures Database (MsigDB v7.0)^103^. We evaluated Bonferroni-corrected significance, adjusted for the number of tested gene sets (one-sided *P* < 3.23×10^-6^) (**Supplementary Table 22**). Lastly, a gene property analysis tested the relationships between 54 tissue-specific gene expression profiles and gene associations, while adjusting for the average expression of genes per tissue type as a covariate (**Supplementary Table 23**), and between brain gene expression profiles and gene associations across 11 brain tissues from BrainSpan^104^ (**Supplementary Table 24**). Gene expression values were log_2_ transformed average Reads Per Kilobase Million (RPKM) per tissue type (after replacing RPKM > 50 with 50) based on GTEx RNA-seq data^105^. We evaluated Bonferroni-corrected significance, adjusted for the number of tested profiles (onesided *P* < 9.26×10^-4^).

We used an extension of MAGMA: “Hi-C coupled MAGMA” or “H-MAGMA” ^30^, to assign non-coding (intergenic and intronic) SNPs to cognate genes based on their chromatin interactions. Exonic and promoter SNPs were assigned to genes based on physical position. We used four Hi-C datasets provided with the software, derived from adult brain^106^, fetal brain^107^, and iPSC derived neurons and astrocytes^108^. We evaluated Bonferroni-corrected significance, adjusted the number of tests within each of the four Hi-C datasets (one-sided *P* < 9.83-9.86×10^-7^) (**Supplementary Tables 25–28**).

The method S-PrediXcan v0.6.2^109^ was used to analyze the association of *EXT* with gene expression levels in different brain tissues. We used pre-computed tissue weights from the Genotype-Tissue Expression (GTEx, v8) project database as the reference transcriptome dataset^105^. As input data, we used the *EXT* summary statistics, LD matrices of the SNPs (available at the PredictDB Data Repository, http://predictdb.org), and transcriptome tissue data related to 13 brain tissues: anterior cingulate cortex, amygdala, caudate basal ganglia, cerebellar hemisphere, cerebellum, cortex, frontal cortex, hippocampus, hypothalamus, nucleus accumbens basal ganglia, putamen basal ganglia, spinal cord and substantia nigra. We evaluated transcriptome-wide significance at the two-sided test *P* < 2.77×10^-7^, which is the Bonferroni-corrected threshold adjusted for 13 tissues times 13,876 tested genes (180,388 gene-tissue pairs) (**Supplementary Table 29**). In **Supplementary Table 30** we summarize the genes findings across the bioannotation analyses.

## Acknowledgements

This research was carried out under the auspices of the Externalizing Consortium. The study was classified as secondary research of de-identified subjects, and the study was awarded ethical approval by the internal review board (IRB) of Virginia Commonwealth University (VCU), with reference number HM20019386. These analyses were made possible by the generous public sharing of summary statistics from published GWAS from the Psychiatric Genomics Consortium (PGC), the Million Veterans Program (MVP), the International Cannabis Consortium (ICC), the GWAS & Sequencing Consortium of Alcohol and Nicotine use (GSCAN), the Social Science Genetics Association Consortium (SSGAC), the Genetics of Personality Consortium (GPC), and the Broad Antisocial Behavior Consortium (BroadABC). We would like to thank the many studies that made these consortia possible, the researchers involved, and the participants in those studies, without whom this effort would not be possible. We would also like to thank the research participants and employees of 23andMe for making this work possible. This research was conducted in part using the UK Biobank Resource under applications 40830 and 11425. We thank all UKB cohort participants for making this study possible. We thank Lea K. Davis for providing access to BioVU. Finally, we thank the Collaborative Study on the Genetics of Alcoholism (COGA), Principal Investigators B. Porjesz, V. Hesselbrock, H. Edenberg, L. Bierut, includes eleven different centers: University of Connecticut (V. Hesselbrock); Indiana University (H.J. Edenberg, J. Nurnberger Jr., T. Foroud; Y. Liu); University of Iowa (S. Kuperman, J. Kramer); SUNY Downstate (B. Porjesz); Washington University in St. Louis (L. Bierut, J. Rice, K. Bucholz, A. Agrawal); University of California at San Diego (M. Schuckit); Rutgers University (J. Tischfield, A. Brooks); Department of Biomedical and Health Informatics, The Children’s Hospital of Philadelphia; Department of Genetics, Perelman School of Medicine, University of Pennsylvania, Philadelphia PA (L. Almasy), Virginia Commonwealth University (D. Dick), Icahn School of Medicine at Mount Sinai (A. Goate), and Howard University (R. Taylor). Other COGA collaborators include: L. Bauer (University of Connecticut); J. McClintick, L. Wetherill, X. Xuei, D. Lai, S. O’Connor, M. Plawecki, S. Lourens (Indiana University); G. Chan (University of Iowa; University of Connecticut); J. Meyers, D. Chorlian, C. Kamarajan, A. Pandey, J. Zhang (SUNY Downstate); J.C. Wang, M. Kapoor, S. Bertelsen (Icahn School of Medicine at Mount Sinai); A. Anokhin, V. McCutcheon, S. Saccone (Washington University); J. Salvatore, F. Aliev, B. Cho (Virginia Commonwealth University); and Mark Kos (University of Texas Rio Grande Valley). A. Parsian and H. Chen are the NIAAA Staff Collaborators. All studies included in the externalizing GWAS are listed in the Supplementary Information.

## Author contributions

D.M.D. and P.D.K. conceived the study. The study protocol was developed by D.M.D., K.P.H., R.K.L., P.D.K, T.T.M., and A.A.P. D.M.D., K.P.H., P.D.K., and A.A.P. jointly oversaw the study. D.M.D. and R.K.L. led the writing of the manuscript, with substantive contributions to the writing from K.P.H., P.D.K. and A.A.P. R.K.L. and T.T.M. were the lead analysts, responsible for conducting genome-wide association studies, quality control, meta-analysis, genetic correlations, and multivariate analyses with Genomic SEM, with assistance from A.D.G. R.K.L. performed the proxy-phenotype analyses. P.B.B. led the polygenic score analyses, and R.K.L. and T.T.M. contributed to those analyses. S.S-R performed the PheWAS in BioVU. S.S-R. led the bioinformatics analyses, and R.K.L contributed to those analyses. P.B.B., R.K.L, T.T.M., and S.S-R. prepared the tables and figures, with assistance from M.N.D, J.W.M., and H.E.P. J.J.T, E.C.J., M.L., H.Z., R.K., and J.A.P. prepared cohort-level GWAS meta-analyses under supervision of K.J.H.V., D.J.L., S. V., H.R.K., and J.G. K.M.H. assisted with analyses performed in the AddHealth study cohort. A.D.G., E.T-D., and I.W. provided helpful advice and feedback on various aspects of the study design. All authors contributed to and critically reviewed the manuscript. R.K.L., T. T.M, P.B.B., and S.S-R. made especially major contributions to the writing and editing; these authors contributed equally.

## Data and code availability

All data sources are described in the Supplementary Information. No new data was collected as part of this study. Only data from existing studies or study cohorts were analyzed, some of which are restricted access to protect the privacy of the study participants. GWAS summary statistics for the externalizing (*EXT*) GWAS (our main discovery analysis) can be obtained by following the procedures detailed at https://externalizing.org/request-data/. The summary statistics are derived from analyses based in part on 23andMe data, for which we can only publicly report results for up to 10,000 SNPs. The full set of externalizing GWAS summary statistics can be made available to qualified investigators who enter into an agreement with 23andMe that protects participant confidentiality. Once the request has been approved by 23andMe, a representative of the Externalizing Consortium can share the full set of summary statistics. All code necessary to replicate this study is available upon request.

## Additional information

Supplementary Information is available for this paper. Online Content Methods, along with any additional Extended Data display items and Source Data, are available in the online version of the paper; references unique to these sections appear only in the online paper. Correspondence and requests for materials should be addressed to Richard Karlsson Linnér at.

