## Supplementary Information for "Multivariate genomic analysis of 1.5 million people identifies genes related to addiction, antisocial behavior, and health"

### Supplementary Methods

#### 1 Study introduction

The externalizing spectrum is a constellation of co-occurring behaviors and disorders that are characterized by under-controlled or impulsive action[1,2]. Central externalizing behaviors include aggression, delinquency, and conduct problems[3]. It has been observed that childhood externalizing precedes various health-risk behaviors later in life, such as smoking, drinking, and illicit substance use[4]. Externalizing psychopathology encompasses multiple clinical diagnoses across development[5], including attention deficit hyperactivity disorder (ADHD), conduct disorder (CD), oppositional defiant disorder (ODD), antisocial personality disorder (ASPD), alcohol dependence (AD), and other substance use disorders (SUDs). Considered together, externalizing behaviors and disorders impose a significant public health burden[6–8].

Multiple twin and family studies have found that much of the genetic influence on any one externalizing disorder is broadly shared with other externalizing spectrum traits and with personality traits that are characterized by behavioral disinhibition or low self-control[9,10]. For example, nearly 70% of the heritability of alcohol dependence is suggested to operate via a general externalizing disposition, rather than via genes specific to alcohol dependence[11]. Here, we broadly refer to a range of clinical and non-clinical traits related to the externalizing spectrum as "externalizing phenotypes" (a detailed working definition is given below).

Previous efforts to identify specific genes involved in a general externalizing liability have been hampered by limited sample size. To surpass that limitation, here we performed multivariate analyses of large-scale genome-wide association studies (GWAS) on externalizing phenotypes with the goals of (a) estimating a genetic factor structure underlying the externalizing spectrum, (b) identifying single-nucleotide polymorphisms (SNPs) and genes primarily involved in a shared genetic liability to externalizing rather than genes that are unique to specific externalizing phenotypes, and (c) increasing the accuracy of polygenic scores for specific externalizing traits that are intractable to study in large samples. The current study was performed according to a preregistered analysis plan, the first version of which was time-stamped on November 8, 2018 (https://doi.org/10.17605/OSF.IO/XKV36).

##### 1.1 Study summary

In this section, we report a brief and illustrative overview of the study procedure, while the remainder of this **Supplementary Information** thoroughly describes all methods and results. The study procedure can broadly be categorized into three major stages:

Stage 1. We amassed a set of phenotype-specific GWAS summary statistics for different externalizing phenotypes, either by collecting existing results or by performing GWAS in UK Biobank (UKB)[12] (**Supplementary Information section 2**). The multivariate method "genomic structural equation modelling" (Genomic SEM)[13] was applied on a subset of the summary statistics ($N$ = 53,293–1,251,809) deemed adequately heritable and statistically powered, in order to estimate a series of model specifications representing different genetic factor structures (**Supplementary Information section 3**). The best-fitting and most parsimonious solution ("the preferred model specification") specified a single common genetic factor with seven indicator phenotypes (which we hereafter refer to as "the latent genetic externalizing factor", or simply, "the externalizing factor"). We estimated genetic correlations between the externalizing factor and 92 other traits from various research domains. Our main discovery analysis is a GWAS on the latent genetic externalizing factor, which we henceforth refer to as "the externalizing GWAS" ($N_{eff}$ = 1,492,085). The externalizing GWAS results were first clumped and then subjected to "conditional and joint multiple-SNP analysis" (GCTA-COJO) to identify a set of "579 jointly associated lead SNPs", which we consider to be our main GWAS findings.

Stage 2. The results of the externalizing GWAS were utilized to perform proxy-phenotype analyses of antisocial behavior and alcohol use disorder[14] (**Supplementary Information section 4**). Similarly, the results were used for polygenic score analyses of a variety of behavioral, health, criminal justice, and substance use measures[15], including a phenome-wide association study (PheWAS) of electronic-health records in the biorepository of the Vanderbilt University Medical Center (BioVU)[16,17] (**Supplementary Information section 5**).

Stage 3. Bioannotation of the externalizing GWAS was performed with the methods "functional mapping and annotation of genetic associations" (FUMA)[18], "multi-marker analysis of genomic annotation" (MAGMA)[19], "Hi-C coupled MAGMA" (H-MAGMA)[20], and "S-PrediXcan"[21,22] (**Supplementary Information section 6**).

### 2 GWAS on externalizing phenotypes

This section, **Supplementary Information section 2**, details the procedure to gather and generate GWAS summary statistics that were later used as input phenotypes in our Genomic SEM analyses (**Supplementary Information section 3**). In summary of this section, the analysis plan delineated a detailed working definition of externalizing phenotypes (including both behaviors and disorders) that we considered to be suitable candidates to represent individual differences in externalizing liability. Based on this definition, on November 8, 2018, we preregistered a set of existing GWAS summary statistics that we had identified in a search of the published GWAS literature. We also specified for inclusion a couple of ongoing studies that we were aware of but that were not yet published. Also, to increase the number of potential input phenotypes, the analysis plan specified that we would perform GWAS on four externalizing phenotypes in UKB, and we excluded a subset of UKB participants from all discovery stage summary statistics to be withheld for follow-up analyses (see below). A quality-control protocol was applied to keep only high-quality single-nucleotide polymorphisms (SNPs). Lastly in this section, we applied LD Score regression to evaluate the power of the GWAS signal, SNP heritability, and the extent of confounding bias from population stratification[23,24], in order to select an adequately powered and heritable subset of summary statistics ($N$ = 53,293–1,251,809) that were retained to be used for multivariate analyses with Genomic SEM.

#### 2.1 Definition of externalizing phenotypes

Psychiatric disorders are commonly comorbid with one another[25]. Patterns of psychiatric comorbidity can be parsimoniously represented in terms of *latent factors* – statistical entities that are not directly observed and that represent broad groupings of disorders that are particularly likely to be comorbid with one another, both contemporaneously and across the lifespan[26]. Factor models of clinically-defined disorders typically differentiate between *internalizing* (characterized by maladaptive fear and withdrawal, such as major depressive disorder or generalized anxiety disorder) and *externalizing* disorders (characterized by under-controlled or impulsive behavior, such as attention deficit/hyperactivity disorder)[5,27].

The psychiatric disorders of childhood in which the cardinal symptoms are under-controlled or impulsive behaviors are (1) attention deficit hyperactivity disorder (ADHD), (2) conduct disorder (CD), and (3) oppositional defiant disorder (ODD). Previous twin research has found evidence for shared genetic influences on these disorders[28–30]. CD, in turn, has been extensively investigated vis-à-vis other psychiatric disorders of adulthood. For example, history of CD in childhood or adolescence is a requirement for a Diagnostic and Statistical Manual (DSM-5) diagnosis of antisocial personality disorder (ASPD), and twin studies have found evidence for genetic overlap between CD and/or ASPD and substance use disorders (SUDs), including alcohol dependence, nicotine dependence, and drug dependence[1,11,31,32]. There is evidence that CD represents an earlier developmental manifestation of the genetic predisposition that impacts SUDs at a later developmental period once there is increased access to alcohol and other drugs[33,34].

Informed by previous multivariate twin research, our analyses therefore aimed to include GWAS of the following psychiatric disorders: ADHD, CD, ODD, ASPD, and SUDs. In addition to clinically-defined disorders, we also consider GWAS of self-reported *symptoms* of these disorders. Previous genetic research on ADHD has found evidence for strong genetic overlap between clinically-defined disorders and quantitative symptom variation within the general population[35]. In the case of SUDs, we also aimed to include GWAS of alcohol and other drug use initiation, as well as quantity/frequency measures of consumption, which show considerable genetic overlap with SUD problems[36].

Further, individuals with externalizing disorders engage in higher rates of health risk behaviors, including *reckless driving* and *risky sexual behavior*[37]. Previous twin research has found that driving while drunk, earlier age at first sex, and measures of riskier sex are all genetically correlated with antisocial behaviors[38–40], and the same literature has also found evidence that genetic liability to externalizing is indexed by the personality traits of *novelty seeking*, *sensation seeking*, lack of *agreeableness*, and lack of *conscientiousness*[1,11,28,41]. Therefore, we also aimed to include GWAS on risky behaviors or personality. Finally, based on the externalizing literature[42–45], the analysis plan listed GWAS on educational attainment and smoking initiation as two traits that could potentially proxy for genetic externalizing liability, with the advantage of being available in huge GWAS samples[46,47].

Putting these lines of psychiatric, psychometric, developmental, and epidemiological research together, our analyses aimed to broadly include GWAS of externalizing disorders and their symptoms, as well as measures of substance use, health risk behaviors, and personality traits. We refer to this category of traits as *externalizing phenotypes*. In the following sections, we report the externalizing phenotypes that we included in the Genomic SEM analyses.

##### 2.1.1 Excluded phenotypes

While we took an inclusive approach to phenotype selection, there are several categories of psychological/psychiatric phenotypes extraneous to the externalizing spectrum that we did not consider. These are briefly outlined with examples below.

- Neurodevelopmental and obsessive-compulsive disorders
    - Examples: autism spectrum disorder, obsessive-compulsive disorder, Tourette syndrome, dyslexia.
    - Note: While ADHD is often conceptualized as a neurodevelopmental disorder, it is also conceptualized as a disruptive behavioral disorder and a core externalizing disorder. Thus, it will be included in analyses.
- Psychotic disorders and symptoms
    - Examples: schizophrenia, bipolar disorder, mania, psychosis.
- Affective disorders and symptoms
    - Examples: major depressive disorder, anxiety disorder, tiredness, loneliness, mood swings.
    - Note: Irritability is a non-specific trait/symptom that will be included in analyses. While it is affective in nature, it is highly relevant to the externalizing spectrum, as it is present in many disorders such as ADHD, oppositional defiant disorder, conduct disorder, substance use disorder, antisocial personality disorder, etc.

- Trauma and stressor-related disorders and symptoms
  - Examples: posttraumatic stress disorder, witness to traumatic experiences, victim of sexual or physical violence, combat exposure.
- Eating disorders and related pathology
  - Examples: anorexia nervosa, binge eating, obesity.

#### 2.2 Collecting GWAS on externalizing phenotypes

**Supplementary Table 1** reports all GWAS on externalizing phenotypes that we considered as potential candidates for inclusion in Genomic SEM. To find these GWAS results, we searched several prominent online GWAS repositories based on the above definition of externalizing phenotypes, restricted to studies in European-ancestry samples with $N > 15{,}000$. The search was conducted in the following resources: the NHGRI-EBI GWAS Catalog[48,49], the LD Hub database of the Broad Institute[50], and the repositories of the Genetics of Personality Consortium (GPC)[51] and the Psychiatric Genomics Consortium (PGC)[52], in the month of June, 2018. Also, we sent out invitations to collaborate addressed to the principal investigators of ongoing studies that we were aware of but that were not yet published. The following research consortia or institutes kindly contributed results from their at-the-time ongoing research efforts (the references refer to the now published studies): the PGC[53,54], the GWAS and Sequencing Consortium of Alcohol and Nicotine use (GSCAN)[47], the Million Veterans Program (MVP)[55], and the International Cannabis Consortium (ICC)[56].

In addition, 23andMe kindly shared GWAS results that they had contributed to ongoing or published studies on impulsivity (the "Barratt Impulsiveness Scale", BIS; and the "Urgency, Premeditation (lack of), Perseverance (lack of), Sensation Seeking, Positive Urgency, Impulsive Behavior Scale", UPPS-P), alcohol use disorder identification test (AUDIT), delay discounting, marijuana initiation (referred to here as "lifetime cannabis use"), and drug experimentation[47,56–60]. However, the sample sizes of most of these GWAS were relatively small ($N \sim$ 20,328–23,127), and only lifetime cannabis use was later included as an indicator phenotype in Genomic SEM, as part of a meta-analysis with other study cohorts that contributed to a recent GWAS, by the ICC[56]. Nonetheless, we instead utilized the other summary statistics with smaller sample size to estimate genetic correlations with the latent externalizing factor (**Supplementary Information section 3**). Also, 23andMe shared their contribution to the GSCAN Consortium's recent GWAS meta-analysis on lifetime smoking initiation (among other drinking and smoking phenotypes), and lifetime smoking initiation was included as an indicator in Genomic SEM.

Beyond collecting existing GWAS, we also performed GWAS in UKB on four externalizing phenotypes that were considered for inclusion in Genomic SEM: (1) addictive behaviors, (2) age at first sex, (3) Alcohol Use Disorder Identification Test Problem scores (AUDIT-P), and (4) irritability. Of note, we defined two UKB Hold-out cohorts of individuals that were excluded from all GWAS included in Genomic SEM, which were instead analyzed in the proxy-phenotype and polygenic score analyses (**Supplementary Information sections 4–5**). We give a detailed definition of the UKB Hold-out cohorts below. In other words, from the four aforementioned GWAS, as well as from any of the existing GWAS (or GWAS meta-analysis) that had analyzed UKB data, we excluded the held-out participants (and their genetic relatives) by re-estimating the existing GWAS (or GWAS meta-analysis) using the same phenotype definition as in the original study. See below for details on the GWAS protocol we applied in UKB. This procedure applies to the following existing GWAS that were considered for inclusion: automobile speeding propensity[9], drinks per week[9], educational attainment[46], lifetime cannabis use[56], lifetime smoking initiation[47], general risk tolerance[9], and number of sexual partners[9].

After applying our quality-control protocol and meta-analysis (described below), but before performing analyses with Genomic SEM, we decided to exclude a few GWAS because of negligible heritability or GWAS association signal. We did this to avoid zeros on the diagonal of the genetic covariance matrix ($S_{\mathrm{LDSC}}$), as well as noise in the sampling covariance matrix ($V_S$) which could have negatively influenced the precision of the Genomic SEM analyses[13]. Specifically, we excluded addictive behaviors because the genetic variance component (pseudo-$h^2$) estimated with BOLT-LMM (see below) was not statistically distinguishable from zero[61]; and we excluded GWAS for which we estimated (a) LD Score regression $h^2$ less than 0.05 and/or (b) GWAS mean $\chi^2$ less than 1.05. In summary, the following GWAS summary statistics were excluded because of not satisfying either or both of these conditions: the Barratt Impulsiveness Scale (BIS-11), the UPPS-P Impulsive Behavior Scale, drug experimentation[59], delay discounting[60], and Alcohol Use Disorders Identification Test Total score (AUDIT-T)[57,58], by 23andMe; as well as agreeableness and conscientiousness by the GPC[62]. After deciding about these exclusions, we amended and registered a second version of the analysis plan (OSF March 29, 2019) before proceeding with any further analyses.

Further, the second version of the analysis plan specified that we would meta-analyze GWAS summary statistics on Alcohol Use Disorder Identification Test Consumption scores (AUDIT-C) and alcohol use disorder (AUD), which were contributed by MVP, with other alcohol-related phenotypes with which they were highly genetically correlated, in order to avoid redundant elements and rank deficiency in the empirical genetic covariance matrix of Genomic SEM ($S_{\mathrm{LDSC}}$). However, our final data sharing agreement with MVP did not allow us to include their summary statistics in a meta-analysis with other cohorts. In addition, we identified that their results included a smaller than expected number of SNPs after applying our quality-control protocol. Specifically, only about 3.9 million SNPs remained after quality control (the number of SNPs in the other indicator GWAS ranged from 6.4–9.5 million). Thus, including the MVP GWAS as indicators in Genomic SEM would have drastically restricted the number of SNPs in the externalizing GWAS. Also, as we explain in **Supplementary Information section 3**, non-problematic drinking phenotypes (such as AUDIT-C) were initially considered for inclusion in the exploratory Genomic SEM analysis, but non-problematic drinking phenotypes were not retained in our preferred model specification. These issues led to the decision to exclude the two MVP GWAS from the discovery stage, and we instead preregistered in the third and final version of the analysis plan (OSF October 28, 2019) that we would retain the summary statistics on AUD for proxy-phenotype analyses (**Supplementary Information section 4**).

Notably, we only succeeded to identify a single childhood externalizing disorder that satisfied our primary sample-size threshold ($N > 15{,}000$): a GWAS on ADHD by the PGC ($N = 53{,}293$)[53], which emphasizes how limited the samples sizes are of studies on this constellation of childhood behavioral disorders. Thus, we did not include neither CD nor ODD as indicators in Genomic SEM, as we had originally intended. Also, we identified a published GWAS on broad antisocial behavior by the Broad Antisocial Behavior Consortium ($N = 16{,}400$)[63], which is a central externalizing phenotype in adulthood. However, to be able to evaluate whether the externalizing GWAS could actually tag genetic signal for a central externalizing trait that was not included in the discovery stage, we preregistered that we would exclude antisocial behavior from the discovery stage, and that we would instead use these summary statistics for proxy-phenotype analyses (**Supplementary Information section 4**). With respect to SUDs, our search could only identify adequately-sized GWAS on alcohol dependence or alcohol use disorder, as well as lifetime cannabis use, but no other adequately-sized GWAS on SUDs or drug initiation measures.

At this stage, we had collected or generated eleven phenotype-specific GWAS (or GWAS meta-analysis) summary statistics that satisfied our inclusion criteria and were forwarded for an exploratory analysis with Genomic SEM: (1) ADHD ($N$ = 53,293), (2) age at first sexual intercourse ($N$ = 357,187), (3) problematic alcohol use ($N$ = 164,684), (4) automobile speeding propensity ($N$ = 367,151), (5) alcoholic consumption (drinks per week; $N$ = 375,768), (6) educational attainment ($N$ = 725,186), (7) lifetime cannabis use ($N$ = 186,875), (9) lifetime smoking initiation ($N$ = 1,251,809), (9) general risk tolerance ($N$ = 426,379), (10) irritability ($N$ = 388,248), and (11) number of sexual partners ($N$ = 336,121). In **Supplementary Information section 3**, we describe a series of exploratory and confirmatory Genomic SEM analyses that led to the preferred model specification, in which we narrowed down the selection to seven out of the eleven indicators: (1) ADHD, (2) age at first sexual intercourse, (3) problematic alcohol use, (4) lifetime cannabis use, (5) lifetime smoking initiation, (6) general risk tolerance, and (7) number of sexual partners, which were eventually used to estimate the latent genetic externalizing factor. In **Supplementary Table 2**, we report a summary of all individual study cohorts there were part of the final seven GWAS meta-analyses. We approximated a lower bound of the number of independent observations to be 1,373,240, by summing the maximum number of samples contributed by a particular study cohort to either of the seven final GWAS meta-analyses. This should be considered a conservative estimate, as it is likely that non-overlapping samples from the same study cohort were contributed to the different GWAS based on phenotype availability.

##### *2.2.1 Conceptual advances to previous GWAS on externalizing phenotypes*

In a recent GWAS effort by some of the authors[9], genetically correlated measures of self-reported willingness to take risks (general risk tolerance) and four real-world risky behaviors (automobile speeding propensity, drinks per week, number of sexual partners, and smoking initiation) were analyzed. Here, we build upon that earlier work. These previously studied traits were all considered here to be externalizing phenotypes, and thus, eligible for inclusion in Genomic SEM (see **Supplementary Information section 2.1**). We make the following advances in the current study:

First, it is important to note the conceptual differences between risk tolerance and externalizing. Risk tolerance can be thought of as a sub facet of externalizing, but externalizing also includes various psychiatric disorders, normative and abnormal behaviors, and other personality traits (e.g., callous and unemotional traits). Hence, the analyses described here reflect an effort designed to study a much broader construct of human behavior, psychology, health, and well-being.

Second, while large-scale single-trait GWAS have been performed on several externalizing phenotypes (e.g., smoking and drinking), our literature review revealed that virtually no adequately powered GWAS are available for several central externalizing outcomes, such as antisocial personality disorder. This study attempts to bridge that gap by collecting and leveraging the extensive degree of genetic overlap among all externalizing phenotypes we could identify to have been studied in large GWAS, and then applying a multivariate GWAS framework to estimate SNP effects on a shared genetic liability to externalizing, rather than the individual traits. We demonstrate here that this approach is of great benefit to studying the genetic architecture of externalizing disorders that are unavailable in large samples and would otherwise remain elusive.

Third, in our previous study[9], we applied two types of multivariate analyses: (1) a GWAS on the first principal component of four risk taking behaviors in the UKB ($N$ = 315,894), and (2) an MTAG[64] analyses of GWAS summary statistics for general risk tolerance, adventurousness, automobile speeding propensity, drinks per week, ever being a smoker, and the self-reported number of sexual partners across the lifespan. The aim of MTAG is not to identify general associations that are broadly related to all input phenotypes (that is the aim of our current study), but rather to augment phenotype-specific summary statistics. Thus, these prior analyses used different methods, a different set of phenotypes, and a noticeably smaller GWAS sample size compared to the present study. The effective sample size of our current study is about 58% larger than our previous study on general risk tolerance ($N$ = 939,908)[9], 20% larger than the largest input GWAS (smoking initiation; $N$ = 1,251,809), and 28 times larger than the smallest (ADHD; $N$ = 53,293). This leads to a substantial increase in the number of genome-wide significant loci that we can report here (Supplementary Information section 3.5), as well as a substantial increase in the accuracy of polygenic scores that can be derived from the GWAS results (Supplementary Information section 5).

Thus, our current study investigates a partly different set of phenotypes using a different method and a much larger sample size. As a result, we are able to report here novel genetic associations for many phenotypes and substantially improved polygenic scores, including scores for several traits that remained elusive to previous GWAS efforts.

#### 2.3 GWAS protocol in UKB

##### *2.3.1 Phenotype definitions*

To complement the existing GWAS we collected, we performed GWAS on four externalizing phenotypes in UKB: addictive behaviors, age at first sexual intercourse, AUDIT-P, and irritability. The phenotypes were defined in the following way:

**Addictive behaviors**

The addictive behaviors phenotype was defined with the following survey item:

*"Have you been addicted to or dependent on one or more things, including substances (not cigarettes/coffee) or behaviours (such as gambling)?"*

The response options were (1) "Prefer not to answer", (2); "Do not know", (3); "No"; and (4) "Yes". We excluded participants who answered "Prefer not to answer" or "Do not know", and those who answered "Yes" or "No" were coded as cases ($N_{\text{cases}}$ = 7,689) or controls ($N_{\text{cases}}$ = 122,893), respectively. However, this GWAS was excluded from any further analysis because GWAS with linear mixed models estimated a genetic variance component (pseudo-$h^2$) that was not statistically distinguishable from zero[61].

**Age at first sexual intercourse**

The age at first sexual intercourse phenotype has previously been studied in the first release of the UKB genetic data, see refs.[9,65], but to our knowledge, not in the full release. The phenotype definition has previously been described in depth in ref.[9]. In summary, the measure was constructed with the following survey item:

*"What was your age when you first had sexual intercourse? (Sexual intercourse includes vaginal, oral or anal intercourse)"*

The respondents were requested to specify an age in full years, and the answers were subsequently subjected to three validity checks: (a) reject answers less than 3; (b) reject answers greater than participant age; and (c) ask for confirmation for answers less than 12. This GWAS included 357,187 participants and it remained an indicator in the preferred model specification.

**AUDIT-P**

The measure AUDIT-P was defined with 7 items in the Alcohol Use Disorder Identification Test, for more details see ref. [58]. This measure is only available for a subset of UKB participants, as part of the online mental health follow-up[66]. This GWAS ($N$ = 130,999) was later meta-analyzed with a PGC GWAS on alcohol dependence ($N$ = 33,685, excluding our follow-up study cohorts: Add Health and COGA)[54]. This GWAS meta-analysis, which we call "problematic alcohol use" ($N$ = 164,684), remained an indicator in the preferred model specification.

**Irritability**

The inclusion of irritability is based on the rationale that angry/irritable mood is a core symptom of ODD in childhood and is typical of aggressive behavior in adulthood[67] The irritability phenotype was defined with the following survey item in UKB, previously studied in ref.[68]:

*"Are you an irritable person?"*

The response options consist of (1) "Prefer not to answer", (2) "Do not know", (3) "No", and (4) "Yes". We excluded participants who answered "Prefer not to answer" or "Do not know", and those who answered "Yes" or "No" were coded as 1 or 0, respectively. Then, because there are repeated measures available for this item, we averaged each person's response across the measures, similar to previous efforts[9]. This GWAS included 388,248 participants, and it was an indicator in the exploratory Genomic SEM analyses, but not in the final model specification.

##### *2.3.2 Definition of the UKB Hold-out cohorts*

We preregistered that we would define two partly overlapping hold-out samples with UKB participants: (1) the UKB Siblings Hold-out cohort and (2) the UKB Problematic Alcohol Use Hold-out cohort, which were to be excluded from all GWAS used as indicators in Genomic SEM. Instead, the held-out participants were retained to be used for proxy-phenotype (**Supplementary Information section 4)** and polygenic score analyses (**Supplementary Information section 5**). In addition, to avoid overfitting because of relatedness across the discovery and follow-up stages, we also excluded from further analysis anyone genetically related to the held-out individuals (pairwise KING coefficient ≥ 0.0442). The two UKB Hold-out cohorts were defined in the following way:

***UKB Siblings Hold-out.*** We defined this hold-out cohort as all participants with at least one full sibling in the UKB. We thereafter kept respondents that (i) passed the UKB genotype sample quality control, described in ref.[12], (ii) were of European ancestry. After applying these filters, we excluded any family unit for which only one sibling had passed the two filters (this step removed 295 family units with only one remaining sibling). In total, we retained 39,640 full siblings of European ancestry, divided across 19,252 family units. Thus, most family units in UKB only observe data on two siblings, no matter the true underlying family size. The UKB Siblings Hold-out cohort allowed us to perform within-family polygenic score analyses with family-specific intercepts (**Supplementary Information section 5**).

***UKB Problematic Alcohol Use Hold-out.*** We defined this hold-out cohort based on a working definition of problematic alcohol use, which was defined as having either an ICD diagnosis (ICD10: F10.X – Mental and behavioral disorders due to use of alcohol; ICD9: 291.X – Alcoholic psychoses, 303.X – Alcohol dependence syndrome, 305.0X – Nondependent abuse of alcohol), or self-reported alcohol addiction or dependence (UKB data-fields 20404, 20406, 20415). We identified 4,400 non-sibling cases of problematic alcohol use that (i) passed the UKB genotype sample quality control[12], (ii) were of European ancestry. These cases were then pooled with a randomly drawn sibling from each family unit (which could be either a case or control). In this hold-out cohort, we also included the 295 single-sibling family units which we excluded from the UKB Siblings Hold-out cohort. We performed GWAS in the final sample of 4,630 cases and 19,334 controls using our UKB GWAS protocol, described below. The summary statistics from this GWAS were never considered for inclusion in Genomic SEM, but were instead generated to be used for proxy-phenotype analyses (**Supplementary Information section 4**).

##### *2.3.3 Estimating genetic PCs to adjust for population stratification*

To account for population stratification, as is standard in genetic epidemiology[69,70], we included 40 genetic principal components (PCs) as covariates in all GWAS estimated in UKB. Many recent studies have relied on the pre-supplied PCs, described in ref.[12]. But because there are recent reports on potential residual population structure when adjusting for the pre-supplied PCs[71], we instead re-estimated the PCs in a genetically more homogenous sample to better capture subtle population stratification in UKB, by using the software flashPCA2[72].

Our procedure is similar to the method described in ref.[12], but with a narrower inclusion of ancestries. In summary, we re-estimated the PCs using the intersection of individuals that (a) had been used in the original estimation, which had been determined to be unrelated and had passed all sample-level quality control, and (b) that had been determined to be of "White British" ancestry based on their self-reports, as well as the investigation of ancestry performed internally by the UKB organization. Thus, at this stage we considered 337,545 individuals for inclusion, while the pre-supplied PCs had instead been estimated with 407,219 individuals of which 69,674 individuals were not of "White British" ancestry. The latter approach can lead to that the first few PCs tend to capture the largest differences across major ancestral groups rather than the subtler stratification within them[69,70].

Next, we applied SNP and sample quality-control, using thresholds similar to those recommended in refs.[12,73] (i.e., we used directly genotyped SNPs outside of long-range LD regions that were filtered on minor allele frequency $\geq 0.01$, genotyping call rate $\geq 0.02$, and a

Hardy-Weinberg equilibrium threshold ≥ $5\times10^{-6}$, and samples were filtered on missingness rate ≥ 0.05). After this step, there were 322,886 individuals remaining. Next, we applied LD pruning (window size = 1000 kb; variant step-size = 50; $r^2 \geq 0.05$; with the PLINK "indep-pairwise" flag, ref.[74]) to produce a set of 77,355 independent markers before estimating the first 40 principal components. We exported the SNP-loadings for each PC, and the projected the remaining of the 459,635 individuals that self-reported to be "White", "White British", "White Irish", and "Any other white background", which also included any related individuals that were excluded from the estimation.

###### *2.3.4 GWAS*

We performed GWAS with linear mixed models (LMM) as implemented in the BOLT-LMM software, version 2.3.2[61]. The method corrects for a genetic variance component (pseudo-$h^2$), which was estimated using a set of 483,680 directly genotyped, autosomal SNPs that passed the genotype quality-control described in ref.[12]. These SNPs had also been filtered on minor allele frequency (MAF) greater than 0.005, Hardy-Weinberg-Equilibrium (HWE) $P$ value greater than $10^{-16}$, and light LD-pruning (window size = 50 kb; variant step-size = 5; $r^2 \geq 0.9$; ref.[74]).

The GWAS excluded participants (1) that were part of the UKB Hold-out samples (and any relatives, pairwise KING coefficient ≥ 0.0442); (2) that did not self-report to categorize their ethnic background as "White", "White British", "White Irish", or "Any other white background"; (3) whose self-reported sex did not correspond to their genetic sex; (4) that had putative sex chromosome aneuploidy or (5) that did not pass the UKB sample quality-control thresholds, described in detail in ref.[12]; and (6) with missing observations with respect to the outcome or model control variables.

The GWAS included control variables for sex, birth year, sex-specific birthyear dummies, genotyping batch (and effectively array), and the first 40 genetic PCs (that we had estimated ourselves). In practice, we first regressed the phenotype on the covariates with OLS, and then applied GWAS on the residuals from that regression. This approach has been shown to lead to virtually identical results and is nonetheless performed as an initial step in the BOLT-LMM estimating procedure to reduce runtime and computational requirement[61]. We analyzed the third release of the UKB imputed genotype data (UKB), which was imputed by first prioritizing variants available in the Haplotype Reference Consortium (HRC) reference panel[75], and secondly, with variants available in a merged reference panel across the 1000 Genomes and UK10K[12,75–77], which were not available in the HRC reference panel.

In summary, we performed GWAS of the following externalizing phenotypes in UKB (some of which were used as a replacement in meta-analysis to exclude our two UKB Hold-out cohorts from the discovery stage): (1) age at first sexual intercourse ($N$ = 357,187), (2) automobile speeding propensity ($N$ = 367,151), (3) drinks per week ($N$ = 375,768), (4) educational attainment ($N$ = 401,024), (5) general risk tolerance ($N$ = 390,934), (6) lifetime cannabis use ($N$ = 131,862), (7) lifetime smoking initiation ($N$ = 403,349), (8) irritability ($N$ = 388,248), (9) number of sexual partners ($N$ = 336,121), (10) addictive behaviors ($N$ = 130,582; excluded because of zero pseudo-$h^2$), and (11) AUDIT-P ($N$ = 130,999). We performed all GWAS with males and females together. As our main discovery analysis is a GWAS on the latent genetic externalizing factor, we do not report GWAS findings for any of the single-phenotype analyses, except with respect to quality control and LD Score regression analyses, described in the next section.

#### 2.4 Quality control analyses

##### *2.4.1 Main reference panel*

For the purpose of performing quality control and calculating LD between SNPs, we assembled a whole-genome sequenced reference panel (hereafter called "the main reference panel"). The main reference panel is a combination of a subset of European-ancestry samples in the 1000 Genomes phase 3 version 5 reference panel[77], together with the British UK10K reference panel[76]. The main reference panel is thus very similar to the reference panel that was used to impute SNPs in UKB that were not available in HRC, as described in ref.[12]. As we discuss below, the main reference panel can substitute for the HRC, as about 90% of common and low-frequency SNPs overlap between the two, and since the vast majority of samples in the HRC are of European ancestry.

We assembled the main reference panel in the following way. The publicly archived 1000 Genomes phase 3 version 5 whole-genome sequencing data (October 2014 haplotype release) was downloaded from a public FTP server hosted by the 1000 Genomes Project Consortium[a]. The restricted-access UK10K whole-genome sequencing data was downloaded after an application procedure from the European Bioinformatics Institute (EMBL-EBI) European Genome-phenome Archive (EGA). Both datasets had already undergone strict quality control, which has previously been described in depth, see refs. [76,77]. The WGS data consisted of variant call format (VCF) files. We used the software "BCFtools", v.1.3.1[b], for all VCF processing, which is distributed by the Sanger Institute. Genomic positions, as well as reference and alternative alleles, were aligned to the Genome Reference Consortium (GRC) human build 37 reference sequence [78].

Before merging the datasets, similar to other efforts[79], we first restricted the 1000 Genomes samples to samples belonging to either of the three European subpopulations (1) Utah Residents with Northern and Western European Ancestry (CEU), (2) Toscana in Italy (TSI), or (3) British in England and Scotland (GBR), after which 297 samples remained. The UK10K samples are virtually all from the GBR ancestral group[76]. We then restricted both reference panels to bi-allelic SNPs with MAF greater than 0. Based on recommendations in ref.[79], we removed SNPs that were inconsistent between the 1000 Genomes phase 1 version 3 and phase 3 version 5 releases of the reference data. Specifically, we removed 5,112 SNPs with inconsistent reference and alternative alleles (e.g., G/C in phase 1 and G/A in phase 3), as well as 10,513 SNPs with large differences in reference allele frequency (i.e., those exceeding 0.25), which is indicative of flipped reference allele or other potential strand issues.

Next, we merged the datasets per chromosome, and then concatenated the chromosomes across the two datasets. Using PLINK v.1.9b3.29[80], we converted the VCF data to PLINK binary format. We restricted the sample to exclude one member of each pair of individuals with genomic relatedness greater than 0.025 (based on 18,270,102 autosomal bi-allelic SNPs with MAF > 0 that were available for all individuals), after which 3,780 out of 4,078 individuals remained, and we applied filters to remove monomorphic SNPs (i.e. SNPs with MAF = 0). We also removed multi-allelic SNPs without retaining any of them as bi-allelic markers, and we removed the aforementioned SNPs with inconsistent alleles or large differences in allele

[a] ftp://ftp.1000genomes.ebi.ac.uk/vol1/ftp/release/20130502/).

[b] http://samtools.github.io/bcftools/

frequency between 1000 Genomes phase 1 and phase 3. Finally, we removed 3,429 SNPs for which the absolute value of the difference in reference allele frequency was greater than 0.25 between 1000 Genomes phase 3 and the merged reference panel, as an extra precaution against misattributed reference alleles.

In summary, after performing these steps, the main reference panel consisted of 3,780 unrelated, European-ancestry samples, merged across the 1000 Genomes phase 3 version 5 and UK10K Consortium reference panels. The main reference panel covers 46,518,418 bi-allelic SNPs, out of which 9,739,256 are autosomal with MAF greater than 0.005. Of the latter, 8,337,793 are among the 9,283,216 autosomal SNPs with MAF greater than 0.005 in a quality-controlled version of the HRC, described in ref.[9]. Thus, among SNPs that would pass our preregistered MAF-filter (MAF > 0.005, see next section), the main reference panel includes about 90% of the common and low-frequency SNPs available in the HRC, suggesting that the main reference panel can be considered a reasonable substitute.

##### *2.4.2 Quality-control protocol of GWAS summary statistics*

We applied a stringent quality-control protocol with the EasyQC software (version 9.2), which is developed by the GIANT consortium[81]. The protocol is similar to that developed in recent GWAS efforts by the Social Science Genetic Association Consortium (SSGAC)[9,46], while a few aspects of the protocol were modified to align with the Genomic SEM pre-processing step. The main aim was to ensure that only high-quality SNPs were used in the multivariate analyses with Genomic SEM. Also, an important step of the protocol is to align the effect-coded alleles of the input summary statistics to match the non-reference (or alternative) allele reported in the main reference panel, in order to ensure that the direction of effect is consistent across the summary statistics prior to any meta- or multivariate analysis. To exclude variants from further analysis (in case these filters had not already been applied prior to or during GWAS analysis, or in the quality control already performed by the contributing study cohorts), we applied the following filters in chronological order to drop:

1. (i) insertions and deletions (INDELs); (ii) SNPs with missing values for the SNP identifier, the effect-coded and other allele, the association *P* value, the effect-size or its standard error, the effect-coded allele frequency, the sample size, as well as the imputation status or quality; and (iii) SNPs with non-sensical values that were outside of the defined variable range (such as *P* values below zero or above one).
2. SNPs that did not satisfy MAF equal to or greater than 0.005.
3. SNPs with an IMPUTE imputation quality (INFO) score[82] less than 0.9.
4. multi-allelic SNPs, as well as SNPs with duplicated chromosome and base pair positions.
5. SNPs that could not be successfully mapped to the main reference panel.
6. SNPs for which the reported alleles did not match those in the main reference panel.

The result of this filtering of the GWAS summary statistics is reported in **Supplementary Table 3**. For brevity, we only report the results for summary statistics that were eventually considered for inclusion in Genomic SEM. After applying these filters, we investigated several standard diagnostic plots, such as QQ-plots and allele frequency plots, a procedure that has been described in detail elsewhere[9,79]. We found that the allele frequencies reported in the GWAS summary statistics correlated strongly with the main reference panel ($r$ ~ 0.999–1), which alleviates concerns about strand issues and suggests that the summary statistics and the main reference panel match in terms of genetic ancestry. The number of SNPs that deviated more than 0.2 from the reference allele frequency in the main reference panel was trivial (0–3,620), and these were retained for further analyses.

#### 2.5 Meta-analysis and LD Score regression

We used the METAL software[83] to perform sample-size weighted meta-analysis to either (a) mimic an existing GWAS meta-analysis that had included UKB data and from which we excluded individuals to create the UKB Hold-out cohorts, or (b) to meta-analyze similar phenotypes to avoid redundant elements and rank deficiency in the empirical genetic covariance matrix of Genomic SEM ($S_{\mathrm{LDSC}}$). The following externalizing phenotypes were meta-analyzed to remove the UKB Hold-out cohorts from an existing GWAS meta-analysis: educational attainment, general risk tolerance, lifetime cannabis use, lifetime smoking initiation. In the final version of the analysis plan (OSF October 28, 2019), we only specified a single GWAS meta-analysis to avoid redundant elements in the genetic covariance matrix. That is, we meta-analyzed a GWAS on alcohol dependence by the PGC[54] with our own GWAS on AUDIT-P in UKB ($r_g$ = 0.794), which we named "problematic alcohol use".

Next, as the final step before performing analyses with Genomic SEM (**Supplementary Information section 3**), we applied LD Score regression on the GWAS summary statistics to (1) estimate SNP-heritability ($h^2$), (2) evaluate the GWAS signal (mean $\chi^2$), and (3) assess the extent of confounding bias from population stratification by evaluating the LD Score regression intercept and attenuation ratio (described in detail elsewhere, see refs.[23,24]). In **Supplementary Table 4**, we report the LD Score regression estimates for the eleven phenotypes for which we estimated $h^2$ and mean $\chi^2$ greater than 0.05 and 1.05, respectively, and thus, considered for inclusion in Genomic SEM. The eleven indicators were weakly to moderately heritable ($h^2$ ~ 0.053–0.235), showed strong to substantial GWAS signal (mean $\chi^2$ = 1.267–3.152). The intercepts ranged from 1.013 to 1.126, and excluding smoking initiation they ranged from 1.013 to 1.047. The attenuation ratio, which is defined as (Intercept – 1) / (mean $\chi^2$ – 1), ranged from 0.0299 to 0.1129. Taken together, these latter statistics suggest that only a very small proportion of the GWAS signal can be attributed to confounding bias from population stratification, and that a vast majority of the signal is due to polygenic effects.

### 3 Genomic structural equation modeling

Genomic structural equation modeling (Genomic SEM)[13] is a recent statistical method that can model the shared and unique genetic architecture of complex traits by applying conventional structural equation modeling principles to GWAS summary statistics. The method's multivariate framework is robust to sample overlap and allows for greater flexibility and accuracy in specifying and estimating genetic covariance matrices relative to other methods, with the advantage of not requiring individual-level genetic data. Thus, Genomic SEM allows for the discovery of connections between phenotypes not naturally studied together because they span different domains, fields of study, or life stages. In the present study, we applied Genomic SEM to investigate the multivariate genetic architecture of the externalizing spectrum by jointly analyzing up to 11 indicator phenotypes (ordered by abbreviation): attention deficit/hyperactivity disorder (ADHD), problematic alcohol use (ALCP), lifetime cannabis use (CANN), drinks per week (DRIN), automobile speeding propensity (DRIV), educational attainment (reverse-coded[c]; EDUC), age at first sexual intercourse (reverse-coded; FSEX), irritability (IRRT), number of sexual partners (NSEX), general risk tolerance (RISK), and lifetime smoking initiation (SMOK). For details on how we selected specifically these 11 indicators, see **Supplementary Information section 2**. The aim of the analyses reported in this section is three-fold: (i) to identify the genetic factor structure that best represents the genetic architecture of externalizing liability, (ii) to estimate the effects of individual SNPs on the latent factor(s), and (iii) to evaluate whether the estimated SNP effects are homogenous across the discovery phenotypes with respect to the latent factor(s).

#### 3.1 Hierarchical clustering

Hierarchical clustering is a type of cluster analysis that aims to partition features of a dataset into groups, where group membership is determined by within-group features that are similar to one another and dissociable from features in other groups[84]. Cluster analysis can serve as a precursor to structural equation modeling by empirically guiding model specification decisions in factor analysis[85]. The initial preregistered analysis plan (November 8, 2018) specified that we would apply hierarchical clustering to guide the decision of how many factors we would explore in subsequent analyses with Genomic SEM. To this end, prior to any structural equation modeling, we applied a hierarchical clustering algorithm to a matrix of pair-wise genetic correlations ($r_g$) for the 11 phenotypes, estimated with LD Score regression[23,24]. Specifically, we applied the Ward hierarchical clustering algorithm[84], as implemented in the *hclust* function included in the R software environment. The results are reported in **Supplementary Table 5.** The 11 phenotypes displayed moderate-to-substantial genetic overlap with at least one other phenotype (max $|r_g|$ = 0.245–0.773), and the average $|r_g|$ across all pairwise correlations was 0.323. The algorithm identified three clusters to be present in the matrix (ordered by abbreviation within each cluster):

1. Attention deficit/hyperactivity disorder (ADHD), educational attainment (EDUC), age at first sexual intercourse (FSEX), irritability (IRRT), and smoking initiation (SMOK).
2. Problematic alcohol use (ALCP), drinks per week (DRIN).

[c] We reversed the effect sizes in the GWAS summary statistics for educational attainment and age at first sexual intercourse so that we could anticipate positive genetic correlations between these traits with externalizing liability.

3. Lifetime cannabis use (CANN), automobile speeding propensity (DRIV), number of sexual partners (NSEX), general risk tolerance (RISK).

After identifying three clusters[d], we updated and timestamped the preregistered study protocol before proceeding (March 29, 2019). Following the initial preregistered analysis plan (November 8, 2018), these new empirical results led us to test four different factor solutions in the exploratory factor analysis, specifying $1...k + 1$ factors, where $k$ corresponds to the number of clusters identified in the genetic correlation matrix.

#### 3.2 Factor analysis

Factor analysis is a multivariate statistical technique used to explain variance and covariance among sets of observed, correlated variables in terms of unobserved latent factors [86]. By modeling the shared variance amongst observed variables as higher-order latent factors, factor analysis is a useful technique for reducing dimensionality of data and accounting for measurement error in observed variables. Factor analysis of genetic correlation matrices is identical to factor analysis of any other type of observed covariance matrix, in which $k$ observed variables are described as linear functions of $m$ latent variables, such that the model can be expressed as

$$y = \Lambda\eta + \varepsilon$$

where $y$ is a $k \times 1$ vector of observed variables, $\varepsilon$ is a $k \times 1$ vector of observed variable residuals, $\eta$ is a $m \times 1$ vector of latent variables, and $\Lambda$ is a $k \times m$ matrix of factor loadings that relate the observed variables to the latent variables.

Here, we used the *factanal* function of R to conduct an exploratory factor analysis of the genetic correlation matrix (estimated with the *ldsc* function of Genomic SEM) with promax rotation. Results for the exploratory factor analysis are presented in **Supplementary Table 6**. Guided by the hierarchical clustering results described above, we estimated four exploratory factor solutions, specifying between one to four latent factors to capture the observed genetic covariance amongst our 11 phenotypes, while retaining factors that explained at least 15% of the variance (a preregistered threshold). As the fourth factor explained only 12.5% of the variance, the three-factor solution was identified as the most appropriate exploratory factor model.

The pattern of factor loadings estimated with the three-factor model was largely in concordance with the results of the hierarchical clustering. Based on the observed loadings, we broadly characterized the three factors as (i) adult risk-taking phenotypes, on which lifetime cannabis use (CANN), automobile speeding propensity (DRIV), number of sexual partners (NSEX), and general risk tolerance (RISK) loaded strongly ($\lambda$ = 0.534–0.885); (ii) developmentally-relevant phenotypes, on which ADHD, educational attainment (EDUC), and age at first sex (FSEX) loaded strongly ($\lambda$ = 0.811–0.966); and (iii) drinking phenotypes, on which problematic alcohol

[d] The second version of the analysis plan (March 29, 2019) reported a preliminary cluster analysis that included the MVP study cohort. Here we report the most recent analysis that excludes MVP (see **Supplementary Information section 2**), which also identified three clusters with similar membership as those identified in the preliminary analysis.

use (ALCP) and drinks per week loaded strongly (λ ~ 0.784–1.003). Lifetime smoking initiation loaded most strongly with (ii) (λ = 0.472), but also moderately with (i) (λ = 0.347). Among the phenotypes, irritability (IRRT) displayed the weakest factor loadings (the strongest loading was with (iii), λ = 0.187), and substantial unique variation (0.927), which is in accordance with it being the weakest genetically correlated across the 11 phenotypes (max $|r_g|$ = 0.245). These findings suggest that the irritability phenotype may not be optimal for modeling the externalizing spectrum, even though it displayed satisfying heritability and GWAS signal. Also, the results of the exploratory analysis suggest that it is unlikely that a single common factor model will be able to closely approximate the observed genetic covariance matrix of the 11 phenotypes.

#### 3.3 Structural equation modeling

Structural equation modeling is a statistical framework that encompasses an array of modeling and methodological approaches for explaining the variance and covariance structure among sets of variables. While the mathematical background and many applications of structural equation modeling are extensive (see refs.[86,87] for a review), we briefly review below several fundamental principles and how they relate to the Genomic SEM framework.

Structural equation models can be represented as a pair of equations: the *measurement model*, which describes how observed variables relate to latent variables, and the *structural model*, which describes how latent variables relate to one another[13]. As in exploratory factor analysis, $k$ observed variables are described as linear functions of $m$ continuous latent variables. In confirmatory factor analysis, this is referred to as the measurement model, which is expressed analogous to the above equation

$$y = \Lambda\eta + \varepsilon$$

By contrast, a structural model is specified when theory is used to model the associations between latent variables via directed regression coefficients. The structural model can be expressed as

$$\eta = \mathrm{B}\eta + \zeta$$

where B is a $m \times m$ matrix of regression coefficients that relate latent variables to one another and $\zeta$ is a $m \times 1$ vector of latent variable residual variances. In this full structural equation model, the observed sample covariance matrix is represented by a set of parameters that relates observed variables to latent variables, and latent variables to each other in a series of linear equations.

Genomic SEM leverages the above framework to model the genetic covariances between a set of observed phenotypes. Using a two-stage approach, the genetic covariance matrix ($S$) and the sampling covariance matrix ($V_S$) are estimated (Stage 1), and a structural equation model is then estimated by minimizing misfit between the model-implied and empirical genetic covariances (Stage 2). To estimate the genetic covariance matrix and its associated sampling covariance matrix, Genomic SEM uses a multivariable form of LD Score regression. $S$ is a symmetric matrix of order $k$, where $k$ equals the number of observed phenotypes, with diagonal elements representing SNP heritabilities and off-diagonal elements representing genetic covariances between phenotypes. Comprised of $k^* = \frac{k(k+1)}{2}$ nonredundant elements, $S$ can be written as

$$S_{LDSC} = \begin{bmatrix} h_1^2 & & & \\ \sigma_{g1,g2} & h_2^2 & & \\ \vdots & & \ddots & \\ \sigma_{g1,gk} & \sigma_{g2,gk} & \cdots & h_k^2 \end{bmatrix}$$

To obtain unbiased estimates of test statistics and standard errors, the nonredundant elements in the $S$ matrix are then used to construct the asymptotic sampling covariance matrix of the LD Score regression estimates, $V_S$. The matrix $V_S$ is symmetric of order $k^*$, in which diagonal elements are sampling variances and off-diagonal elements are sampling covariances. Thus, it can be written as

$$V_S = \begin{bmatrix} SE(h_1^2)^2 & & & & & \\ cov(h_1^2, \sigma_{g_1,g_2}) & SE(\sigma_{g_1,g_2})^2 & \ddots & & & \\ \vdots & \vdots & & & & \\ cov(h_1^2, \sigma_{g_1,g_k}) & cov(\sigma_{g_1,g_2}, \sigma_{g_1,g_k}) & SE(\sigma_{g_1,g_k})^2 & \ddots & & \\ \vdots & \vdots & \vdots & & & \\ cov(h_1^2, h_j^2) & cov(\sigma_{g_1,g_2}, h_j^2) & cov(\sigma_{g_1,g_k}, h_j^2) & SE(h_j^2)^2 & & \\ \vdots & \vdots & \vdots & & \ddots & \\ cov(h_1^2, \sigma_{g_j,g_k}) & cov(\sigma_{g_1,g_2}, \sigma_{g_j,g_k}) & cov(\sigma_{g_1,g_k}, \sigma_{g_j,g_k}) & cov(h_j^2, \sigma_{g_j,g_k}) & SE(\sigma_{g_j,g_k})^2 & \\ cov(h_1^2, h_k^2) & cov(\sigma_{g_1,g_2}, h_k^2) & cov(\sigma_{g_1,g_k}, h_k^2) & cov(h_j^2, h_k^2) & cov(\sigma_{g_j,g_k}, h_k^2) & SE(h_k^2)^2 \end{bmatrix}$$

The diagonal elements of $V_S$ are then estimated with a jackknife resampling procedure analogous to the procedure used in the original bivariate version of LD Score regression.

The $S$ matrix from Stage 1 is then used in Stage 2 to estimate the parameters of the specified structural equation model with either weighted least squares (WLS) or maximum likelihood (ML) estimators. The estimators minimize misfit between the model-implied and empirical genetic covariances, but differ in how information is weighted (see [13] for further detail). A sandwich correction that incorporates the sampling covariance matrix is used to obtain unbiased standard errors and corresponding test statistics. In this study, all models were estimated using WLS estimation, as described in ref.[13], in which a fit function is optimized using the diagonal elements of $V_S$, standard errors are subsequently adjusted using the off-diagonal elements of $V_S$.

Importantly, the off-diagonal elements of $V_S$ index to what extent the sampling errors across the summary statistics correlate. Thus, just like its predecessor LD Score regression[24,88], Genomic SEM has been shown to be unbiased and robust to varying degrees of, or even complete, sample overlap[13]. Also, the method is capable of handling differences in GWAS sample size. These are important properties, as large-scale GWAS summary statistics are often generated as meta-analyses that span several biobanks and cohort studies.

##### *3.3.1 Confirmatory factor analysis*

Confirmatory factor analysis is a common application of structural equation modeling, where the observed covariances among a set of observed variables are modeled according to both theory and exploratory data inspection. Competing models are tested to identify the model that best fits the data, where good fit reflects that the specified latent variable structure adequately explains the observed covariances among the set of observed variables.

Guided by the results of the exploratory factor analysis, as well as psychiatric and psychometric theory, we examined a series of confirmatory factor models to identify the factor solution that best explained the observed genetic covariances among the set of discovery phenotypes. As a baseline comparison throughout the analyses reported below, we contrasted each specified model with a single common factor model with the 11 indicators (i.e., a single latent dimension of genetic risk for externalizing).

Model fit was assessed using preregistered thresholds for conventional indices in structural equation modeling: the model $\chi^2$ statistic, the Akaike information criterion (AIC), the comparative fit index (CFI), and the standardized root mean square residual (SRMR). All of these indices retain their standard interpretations within a Genomic SEM framework with the exception of the model $\chi^2$ statistic[13]. In large samples like those used in GWASs, $\chi^2$ tests are overpowered and likely to be significant. As such, the model $\chi^2$ statistic was used as a comparative measure of fit to evaluate competing models (akin to AIC), rather than as a measure of statistical significance. For CFI and SRMR, values greater than .90 and less than .08, respectively, were considered reflective of good model fit[89]. Results for the confirmatory factor analysis are summarized below and presented in **Supplementary Table 7**.

***Common factor model (11 indicators)***
As a baseline, we evaluated a common factor model with all 11 phenotypes operating as indicators for a single latent factor. While easily interpretable, this particular model exhibited poor fit, as indicated by model fit indices ($\chi^2(44)$ = 8007.35, AIC = 8051.35, CFI = .662, SRMR = .161). This result is in accordance with the exploratory factor analysis that suggested that a single factor may not be optimal for approximating the observed covariance structure of the 11 phenotypes.

***Correlated factors model (11 indicators)***
We next tested a three-factor model, a decision that was guided by the exploratory analysis, where each phenotype loaded onto three correlated latent factors based on their strongest loading observed in the exploratory factor analysis. No cross-loadings were estimated. Correlations between the latent factors were freely estimated. This simple correlated-factors model did not fit the data well, as indicated by model fit indices ($\chi^2(41)$ = 6152.194, AIC = 6202.194, CFI = .741, SRMR = .126). We then evaluated a correlated factors model that allowed for cross-loadings, retaining all loadings with an absolute value ≥ .30 in the exploratory factor analysis. However, this model also showed suboptimal fit and did not meet our preregistered model fit criteria ($\chi^2(39)$ = 2610.544, AIC = 2664.544, CFI = .891, SRMR = .089).

***Bifactor model (11 indicators)***
We then tested a more complex bifactor model, in which the observed covariance of all phenotypes is modeled as general common factor, but residual variance among sets of indicators is modeled as specific factors. As these residual variance factors are conceptually orthogonal to one another, between-factor covariances were fixed to zero. Here, we modeled three latent factors: a general latent factor of externalizing with all phenotypes as indicators, a specific latent factor with group (ii) developmentally-relevant phenotypes as indicators, and a second specific latent factor with all other phenotypes in groups (i) and (iii) as indicators. This model also exhibited suboptimal fit per our pre-registered criteria ($\chi^2(33) = 3016.033$, AIC = 3082.033, CFI = .874, SRMR = .097), and the resulting factor structure would have been difficult to interpret.

***Revised common factor model (7 indicators)***
Finally, we evaluated a revised and more parsimonious common factor model that only included phenotypes with moderate-to-large (*i.e.*, ≥ .50) loadings on the single latent factor estimated in the common factor model with 11 indicators. That is, we decided to exclude automobile speeding propensity ($\lambda = 0.211$), irritability ($\lambda = 0.270$), educational attainment ($\lambda = 0.273$), and alcohol consumption ($\lambda = 0.373$). We freely estimated correlations between the residual variance in age at first sexual intercourse and lifetime cannabis use, as well as problematic alcohol use and lifetime smoking initiation. These pairs of phenotypes were selected as they had notable loadings in the exploratory factor analysis (*e.g.*, opposite direction of effect for F2 and cross loadings on F3), suggesting there was appreciable covariance not accounted for by a common factor with respect to these pairs. We found that this parsimonious model specification fit the data the best across all tested specifications, and it closely approximated the observed genetic covariance matrix ($\chi^2(12) = 390.234$, AIC = 422.234, CFI = .957, SRMR = .079). This model was selected as our final factor model, as it identified a latent genetic factor of externalizing psychopathology, offered an easily interpretable factor solution, and satisfied our pre-registered selection criteria on the basis of model fit indices, and we hereafter refer to it as "the latent genetic externalizing factor", or simply, "the externalizing factor"

##### *3.3.2 Genetic correlation*

We used Genomic SEM to estimate genetic correlations between the latent genetic externalizing factor and 92 other phenotypes, which were broadly related to four domains: demography, health and medicine, psychopathology, and socioeconomic outcomes. We note that this approach is equivalent to LD Score regression when modeling relationships between observed phenotypes, and more appropriate than LD Score regression when modeling relationships that involve a latent genetic factor. This is due to the fact that Genomic SEM can directly model the covariance between a latent genetic factor and an exogenous phenotype rather than rely on the estimated SNP effects, which may or may not operate via the latent genetic factor (*e.g.*, $Q_{\text{SNP}}$ loci). The selection of *a priori* phenotypes were pre-registered on the Open Science Framework (OSF, October 28, 2019). Genetic correlations results are presented in **Supplementary Table 8** and **Extended Data Fig. 1**.

#### 3.4 Multivariate genome-wide association analyses

Our main discovery analysis is a GWAS on the latent genetic externalizing factor, which we henceforth refer to as "the externalizing GWAS". After identifying the confirmatory factor model that best explained the observed genetic covariances among the externalizing phenotypes, we estimated individual SNP effects on the latent externalizing factor. A brief overview of this multivariate GWAS method is provided below. The method is described in detail in ref.[13].

First, individual SNP effects and their squared standard errors and sampling covariances are included in the genetic covariance matrix ($S$) and the sampling covariance matrix ($V_S$). The genetic covariance matrix is expanded to include covariances between SNP $j$ and the latent genetic components of each phenotype, $g_1$ through $g_k$.

$$S_{Full} = \begin{bmatrix} \sigma^2_{SNP} & & & & & \\ \sigma_{SNP,g_1} & h_1^2 & & & & \\ \sigma_{SNP,g_2} & \sigma_{g_1,g_2} & h_2^2 & & & \\ \sigma_{SNP,g_3} & \sigma_{g_1,g_3} & \sigma_{g_2,g_3} & h_3^2 & & \\ \vdots & \vdots & \vdots & \vdots & \ddots & \\ \sigma_{SNP,g_k} & \sigma_{g_1,g_k} & \sigma_{g_2,g_k} & \sigma_{g_3,g_k} & & h_k^2 \end{bmatrix}$$

The associated sampling covariance matrix, $V_S$, then includes the following: (i) the sampling variances and sampling covariances of the SNP heritabilities and genetic covariances, (ii) the variance of SNP $j$ as derived from reference panel data, and (iii) the sampling covariances of the SNP-genotype covariances. Finally, $m$ models are estimated in order to obtain GWAS summary statistics for the latent factors, where $m$ is the number of SNPs present across all included summary statistics.

We note that unit loading identification is used to set the scale of latent factors for models including SNP effects. This is a difference from the structural equation models without SNP effects, where unit variance identification is used to facilitate easy interpretation of factor loadings (for explanation, see Mallard and colleagues[90]). Here, we set the scale of the externalizing latent factor by fixing the factor loadings of number of sexual partners to one, as it had a strong loading on the common factor and moderate SNP heritability relative to the other phenotypes. Again, we note that this does not have any appreciable effect on the estimated SNP effects and their test statistics, it simply sets the scale of the latent genetic factor.

##### *3.4.1 Effective sample size*

We estimated the effective sample size for a given SNP in a latent factor model following the procedure described by Mallard and colleagues[90]. Just like the overall Genomic SEM framework (see above), this estimator is robust to sample overlap[13]. First, we assumed that the effect of SNP $j$ follows:

$$\beta_j = \frac{Z_j}{\sqrt{n_j \times 2 \times MAF_j\left(1 - MAF_j\right)}}$$

Here, $Z_j$ is the association test statistic, $n_j$ is the unknown effective sample size that we seek to estimate, and $MAF_j$ is the minor allele frequency of SNP $j$. Note that the variance of SNP $j$ ($\sigma_j^2$) is assumed to be $2 \times MAF_j\left(1 - MAF_j\right)$. Therefore, if we know the association test statistic and minor allele frequency of SNP $j$, then we can estimate its effective sample size by solving for $n_j$, which yields

$$n_j = \frac{(Z_j/\beta_j)^2}{\sigma_j^2}$$

As this formula can produce inflated estimates for SNPs with low MAF, we set a lower and upper MAF limit of 10% and 40%, respectively, when estimating effective *N* for the overall multivariate GWAS results ($N_{eff}$). As $N_{eff}$ is approximately equal to the mean $n_j$ for $m$ SNPs with a MAF between $a$ and $b$, this is approximated as

$$N_{eff} \approx \frac{1}{m} \sum_{MAF=a}^{b} n_j$$

We apply this formula to estimate the effective sample size for the latent externalizing factor, yielding an $N_{eff}$ of 1,492,085.

###### *3.4.2 Identifying near-independent and jointly associated lead SNPs*

To define near-independent "lead SNPs", we applied a conventional "clumping" algorithm[79], implemented in the PLINK software[74,91]. The algorithm uses four parameters: a primary (two-sided test) *P*-value threshold ($5\times10^{-8}$), a secondary *P*-value threshold to drop weakly associated SNPs from the procedure ($1\times10^{-4}$), and an $r^2$ threshold (0.1) together with a SNP window defined in kilobases (1,000,000 kb) to assign SNPs to near-independent "clumps", each lead by the most strongly associated SNP, according to *P* value. By setting an extremely wide SNP window, we effectively consider only LD for determining independence between SNPs. LD was calculated with the main reference panel.

Next, to investigate whether the lead SNPs were conditionally and jointly associated with the externalizing factor when considered simultaneously in the same model, we applied the standard method "multi-SNP-based conditional & joint association analysis using GWAS summary data" (COJO)[92], as implemented in the GCTA software[93]. This method is specifically developed for the scenario where it is infeasible to consider the SNPs jointly using individual-level data, and instead uses LD from a reference panel to derive conditional SNP effects. Specifically, we applied the default step-wise model selection procedure on the lead SNPs identified with the clumping algorithm. The selection procedure is described in detail in ref.[92], and it assumes that SNPs located further than 10 million base pairs from each other are in linkage equilibrium ($r^2$ = 0). We consider any SNPs identified by the COJO analysis as conditionally and jointly associated with the externalizing factor to be our main GWAS findings.

#### 3.5 Results of the multivariate externalizing GWAS

With Genomic SEM, we estimated individual SNP effects for 6,132,068 SNPs, which is the intersection of SNPs available after quality control, on the latent genetic externalizing factor ($N_{eff}$ = 1,492,085). We display the GWAS results in a Manhattan plot in **Fig. 1**, and in a quantile-

quantile (Q-Q) plot in **Extended Data Fig. 2**. The externalizing GWAS showed strong association signal, with a mean $\chi^2$ and genomic inflation factor ($\lambda_{GC}$) of 2.98 and 2.22, respectively, when calculated with the ~6 million SNPs, and 3.114 and 2.337, respectively, when restricted to the 1,019,632 SNPs used in LD Score regression. We estimated the LD Score regression intercept and attenuation ratio to be 1.115 ($SE$ = 0.019) and 0.054 ($SE$ = 0.009), respectively, which suggests that almost all of the inflation we observed in the association test statistic is attributable to polygenicity rather than bias from population stratification[13,23].

The clumping algorithm identified 855 near-independent lead SNPs from the 58,896 SNPs that passed genome-wide significance (two-sided test $P < 5\times10^{-8}$). With COJO, we identified that 579 of the 855 lead SNPs were conditionally and jointly associated, meaning they were significantly associated with EXT even after statistically adjusting for each other, as well as the other of the 855 lead SNPs. We consider these 579 conditionally and jointly associated SNPs to be our main GWAS findings. In **Supplementary Table 9**, we report the GWAS and COJO results for the 579 SNPs, together with basic bioannotation with "functional mapping and annotation of genetic associations" (FUMA, ref.[18]), which is further described in **Supplementary Information section 6**.

We investigated whether the 579 SNPs were reported to be genome-wide significant in any of the input GWAS. We did that by identifying the smallest $P$ value reported for each of the 579 SNPs, as well as all correlated SNPs within their linkage disequilibrium (LD) regions ($r^2 > 0.1$). The result of this lookup is reported in **Supplementary Table 9**. We found that 121 (21%) of the 579 SNPs and SNPs in their LD regions were not genome-wide significant any of the seven input GWAS, and 8 (1%) did not reach $P < 1\times10^{-5}$. Thus, the externalizing GWAS could identify SNP-associations that were not previously genome-wide significant in the input GWAS. Finally, we looked up the 579 SNPs and their LD regions ($r^2 > 0.1$) in the GWAS Catalog and found that 41 (7%) could be considered novel GWAS findings, as they had never before been reported to be associated with any trait in the GWAS literature at suggestive significance ($P < 1\times10^{-5}$; **Supplementary Table 10**). We further discuss the GWAS Catalog lookup in **Supplementary Information section 6**.

##### *3.5.1 $Q_{SNP}$ heterogeneity tests*

Variation in the size of factor loadings across discovery phenotypes suggests the possibility that individual SNP effects might not operate (i.e., is statistically mediated) through the common externalizing factor[13]. The null hypothesis of the $Q_{SNP}$ test is that SNP effects on the constituent phenotypes are mediated via a common pathway through the *EXT* factor, so a significant $Q_{SNP}$ test indicates that SNP association is better explained by a trait-specific pathway independent of the *EXT* factor. To evaluate this potential heterogeneity in SNP effects, we estimated genome-wide $Q_{SNP}$ statistics for each SNP in the multivariate GWAS, which are $\chi^2$-distributed test statistics. As described by Grotzinger and colleagues[13], larger values for $Q_{SNP}$ reflect a violation of the null hypothesis that a SNP is mediated through the latent factor(s). Genome-wide results for 6,107,583 $Q_{SNP}$ tests for which the method converged are presented in a Manhattan plot in **Fig. 1D** and a Q-Q plot in **Extended Data Fig. 2**.

We applied the clumping algorithm described above to the $Q_{SNP}$ results and found 160 near-independent genome-wide significant (two-sided test $P < 5\times10^{-8}$) $Q_{SNP}$. The strongest and most salient example of a trait-specific association is SNP rs1229984 (two-sided $Q_{\mathrm{SNP}}$ $P = 1.67\times10^{-51}$; two-sided GWAS $P$ with EXT = 0.022). This particular SNP, located in the gene *ADH1B*, is a known missense variant with a well-established role in alcohol metabolism[94], and it is only associated with a single input phenotype—problematic alcohol use (two-sided GWAS = $6.43\times10^{-57}$). Notably, however, we found that 99% (571/579) of the conditionally and jointly associated SNPs were not among the 10,665 genome-wide significant $Q_{\mathrm{SNP}}$ (**Supplementary Table 9**). That is, there was strong evidence that the 579 genetic variants we identified in the externalizing GWAS really capture a unitary dimension of genetic liability rather than simply representing an amalgamation of variants with divergent associations with the constituent phenotypes. We estimated mean $\chi^2$ and genomic inflation factor ($\lambda_{\mathrm{GC}}$) of the $Q_{\mathrm{SNP}}$ results to be 1.956 and 1.864, respectively, when calculated with the ~6 million SNPs, and 2.013 and 1.9424, respectively, when restricted to the 1,016,650 SNPs used in LD Score regression. Thus, the $Q_{\mathrm{SNP}}$ analysis was sufficiently powered to identify substantial heterogeneity across the genome, but reassuringly, not with respect to the vast majority of our main findings. This aligns with the expectation of modelling a latent common factor with SEM, which is that the common factor should primarily identify shared variance and not the unique features of the model indicators. Finally, we estimated an LD Score regression intercept of 0.956 ($SE$ = 0.013), which suggests that the inflation we observed in the $Q_{\mathrm{SNP}}$ test statistic is not attributable to bias from population stratification[13,23].

### 4 Proxy-phenotype and quasi-replication analyses

In this section, we report a series of proxy-phenotype and quasi-replication analyses. The proxy-phenotype method is a two-stage stage approach that leverages SNP associations identified in a well-powered, first-stage GWAS on a proxy phenotype (here, the externalizing GWAS), as empirically plausible candidates that can then be tested for association in independent, second-stage GWAS samples on genetically correlated phenotypes[79,95]. The smaller number of hypotheses that are tested in the second stage yields an advantage in terms of statistical power compared to evaluating significance at the genome-wide significance threshold in the second-stage GWAS samples. This approach has proven advantageous in situations where there is no independent, adequately-sized GWAS sample available to study a trait of interest directly, as well as to perform SNP-level "quasi-replication" when no independent replication sample of the same phenotype exists[79,95], the latter of which is the case for our current study.

Here, the externalizing GWAS was used as the first-stage GWAS. To avoid overfitting, the second-stage GWAS samples were not part of the externalizing GWAS. As second-stage phenotypes, we studied two central externalizing traits for which we estimated moderate-to-substantial genetic overlap with the externalizing GWAS: (1) antisocial behavior (ASB; $r_g$ = 0.69, *SE* = 0.08) and (2) alcohol use disorder (AUD; $r_g$ = 0.52[e], *SE* = 0.03). Notably, ASB was not an indicator phenotype in the Genomic SEM analyses while a GWAS on problematic alcohol use was an indicator. By performing this analysis, we aimed to (a) holistically "quasi-replicate" the 579 jointly associated lead SNPs (**Supplementary Information section 3**) and (b) perform an informed search to potentially identify novel SNPs enriched for association with the second-stage phenotypes. To our knowledge, as part of this analysis, we generated the largest meta-analysis of GWAS on antisocial behavior to date (*N* = 32,574), a trait for which there are still no genome-wide significant findings reported in the NHGRI-EBI GWAS Catalog (Buniello et al. 2018; accessed on March 31, 2020).

#### 4.1 Methods

##### *4.1.1 Auxiliary GWAS meta-analyses of the second-stage phenotypes*

We generated a meta-analysis of GWAS on ASB. First, we performed GWAS adjusted for age, sex, genetic PCs, and technical covariates in three hold-out cohorts: (1) Add Health (*N* = 4,884), (2) COGA (*N* = 6,323), and (3) PNC (*N* = 4,142). In Add Health, the ASB phenotype was defined as a continuous measure of the average of the rule-breaking/delinquency scale across four waves, and GWAS was performed with OLS in unrelated individuals. In COGA, the phenotype was the maximum DSM-IV criteria count of either ASPD (adulthood, 18 or older) or CD (childhood, under age 18) interviews, as ASPD is only assessed in those 18 years and older, and GWAS was performed with linear mixed models. In PNC, ASB was defined as a composite score of conduct disorder symptoms, assessed with the Kiddie Schedule for Affective Disorders

[e] Because of the lower-than-expected number of SNPs available in the MVP summary statistics (see below and **Supplementary Information section 2**), the LD Score regression correlation estimation was performed with only 583,627 SNPs, which we believe may have attenuated the estimate. The genetic correlation between the externalizing GWAS with the PGC alcohol dependence GWAS was estimated to 0.76 (SE = 0.06), which we believe more accurately reflects the true genetic overlap between the two phenotypes.

and Schizophrenia-Present and Lifetime Version (KSADS-PL), and GWAS was performed with OLS in unrelated individuals. After applying the QC protocol described in **Supplementary Information section 2**, we meta-analyzed the newly estimated GWAS together with a published GWAS ($N$ = 16,400) on ASB by Tielbeek et al.[63]. As COGA had contributed data to that previous GWAS, we made sure to exclude that particular subsample ($N$ = 1,379) from our internal GWAS in that cohort. In addition, we included association results for directly genotyped SNPs (imputed genotypes were not available) from one of the replication cohorts in Tielbeek et al., namely, the Michigan State University Twin Research study cohort (MSUTR; $N$ = 825). The total sample size of the meta-analysis was 32,574.

With respect to AUD, we performed proxy-phenotype analyses in a recently published GWAS in the Million Veterans Program (MVP; $N$ = 202,004)[55]. Initially, we planned to use the MVP GWAS results for inclusion as an input in Genomic SEM. However, our final data sharing agreement with MVP did not allow us to meta-analyze their results with other cohorts. We also found a limited overlap of SNPs between MVP and SNPs in our Genomic SEM analyses. After QC in the MVP data (**Supplementary Information section 2**), only about 3.9 million SNPs remained (the number of SNPs in the other indicator GWAS ranged from 6.4–9.5 million). Therefore, in the third version of the analysis plan (OSF October 28, 2019), we amended a change to the study protocol and decided that the MVP GWAS would instead be used for proxy-phenotype analysis on AUD. As a complementary analysis, we also performed an ancillary meta-analysis of internal GWAS on alcohol use disorder/alcohol problems that included three hold-out cohorts: (1) Add Health ($N$ = 4,166), (2) COGA ($N$ = 7,335), and (3) the UKB Problematic Alcohol Use Hold-out cohort (described in **Supplementary Information section 2**; $N$ = 23,937). The total sample size of this ancillary meta-analysis on AUD was 34,426.

###### *4.1.2 Defining the first- and second-stage SNP associations*

We looked up the 579 jointly associated lead SNPs identified in the externalizing GWAS in the independent, second-stage GWAS results on ASB and AUD. The second-stage GWAS were all restricted to SNPs that satisfied at least 80% of the total sample size (the ancillary meta-analysis of GWAS on AUD was restricted based on its own sample size and not the much larger sample size of the GWAS in MVP).

With respect to ASB, we first checked whether the 579 SNPs themselves were immediately available in the second-stage GWAS results. For any SNPs that were missing, we attempted to identify suitable proxy SNPs in high LD ($r^2 > 0.8$). In total, 500 of the 579 SNPs were immediately available, and we could identify 53 suitable proxies for the other 79 missing SNPs. Thus, the proxy-phenotype analysis of ASB used a total of 553 ($k$) SNPs (or their proxies) as "first-stage associations".

With respect to AUD, we prioritized lookups of the 579 SNPs, or suitable proxy SNPs ($r^2 > 0.8$), in the larger MVP AUD GWAS. Only when a SNP was both missing and had no suitable proxy, did we perform lookups in the smaller, ancillary meta-analysis of GWAS on AUD. In total, 409 of the 579 SNPs were directly available in the MVP GWAS on AUD, 39 could be proxied, and the other 131 missing SNPs were immediately available in the ancillary meta-analysis. Thus, the proxy-phenotype analysis of AUD used a total of 579 ($k$) SNPs (or their proxies) as "first-stage associations".

Because we did not perform COJO analysis with the proxy SNPs, for all first-stage associations, we analyzed the direction of effect as estimated in the externalizing GWAS rather than the adjusted effect-size estimates from the COJO analysis. This decision will not influence the results as the correlation between the adjusted and non-adjusted effect sizes was >0.99 and the direction of effect was consistently the same. Before proceeding, we aligned the direction of effect for the second-stage lookups to match the effect-coded allele of the first-stage associations.

###### *4.1.3 Investigation of joint enrichment for association*

As in previous studies[9,79], we investigated whether the first-stage SNP associations with externalizing were more enriched for association with the second-stage phenotypes than an empirical null distribution based on a random sample of near-independent ($r^2 < 0.1$) SNPs from the second-stage GWAS. We perform this test in comparison to an empirical null distribution because we expect that the second-stage phenotypes are polygenic, with many true associations that have yet to reach significance. Therefore, it would be inappropriate to test for enrichment against a uniform (null) $P$ value distribution. For each of the $k$ first-stage associations we looked up ($k$ is 553 and 579 for antisocial behavior and alcohol use disorder, respectively), a sample of 250 near-independent SNPs matched on MAF (± 1 percentage point) was drawn from the second-stage GWAS (with respect to AUD, all SNPs were drawn from the GWAS in MVP). Each set of SNPs were ranked according to $P$ value. Then, as a joint test of whether the first-stage SNPs are more enriched for association with the second-stage phenotypes than the background polygenic signal, we performed a non-parametric (one-sided) Mann-Whitney test of the null hypothesis that the $P$ values of the $k$ SNPs are from the same distribution as the 138,250 and 144,750 SNPs that were drawn from the GWAS on antisocial behavior and alcohol use disorder, respectively.

###### *4.1.4 Holistic quasi-replication*

Because the second-stage GWAS samples are too small to quasi-replicate our findings at genome-wide significance, we had prespecified three tests with the objective to holistically quasi-replicate the first-stage SNPs (or their proxies). The tests are reported in order of descending statistical power. First, we performed a test of sign concordance by investigating whether the direction of effect across the first- and second-stage GWAS were in greater concordance than what could be expected by chance. In the circumstance that the GWAS would be entirely spurious, the expectation is that 50% of the signs would be in concordance simply by chance. Secondly, we tested whether a greater proportion of the first-stage associations were nominally significant (two-sided $P < 0.05$) in the second-stage GWAS than expected under the empirical null distribution. Thirdly, we report associations after Bonferroni correction for the $k$ look-ups we conducted (two-sided $P < 0.05/k$). Any first-stage SNPs that satisfied that last criterion was considered to be "second-stage associations". Any second-stage associations, including any SNPs in weak LD ($r^2 > 0.1$), were looked up in the NHGRI-EBI GWAS Catalog for previously reported associations with the second-stage phenotype itself[48], as well as related phenotypes.

#### 4.2 Results

The results of the proxy-phenotype analyses are reported in **Supplementary Tables 11–12** and displayed in **Extended Data Fig. 3**. For ASB, the Mann-Whitney test rejected the null hypothesis of no enrichment (one-sided $P = 1.10\times10^{-5}$), which suggests that the first-stage associations identified in the externalizing GWAS are more enriched for association with this trait than the background polygenic signal of the second-stage GWAS. Out of 553 first-stage associations, 370 (66.9%) have concordant direction of effect ($H_0 = 276.5$; binomial two-sided test $P = 1.39\times10^{-15}$), and 58 (10.5%) are nominally significant ($P < 0.05$), which is more than would be expected compared to the empirical null distribution (empirical $H_0 = 25.92$ (~4.7%); binomial two-sided test $P = 1.64\times10^{-8}$). These findings suggest that the GWAS are not entirely spurious and that it was possible to identify genetic signal that overlaps with a central externalizing trait, which itself was not an indicator in Genomic SEM. We identified one second-stage association on chromosome 5 at ~87.8 Mb that survived experiment-wide Bonferroni correction (rs10044618; a proxy for rs6452785 at $r^2 = 0.851$) with an ASB GWAS association $P$ value of $8.15\times10^{-5}$, which is more than what is expected under the null ($H_0 = 0.05$, binomial two-sided test $P = 1.81\times10^{-3}$). As there are no previously reported SNPs that are robustly associated with ASB in the GWAS Catalog, if this association would replicate in future studies, then this would be the first SNP association for ASB. The proxied second-stage association, rs6452785, has previously been reported to be associated with smoking initiation in the GWAS Catalog, and various SNPs in weak LD have been reported to be associated with a range of behavioral phenotypes, including depression, neuroticism, and educational attainment.

For AUD, the Mann-Whitney test of joint enrichment strongly rejected the null hypothesis of no enrichment (one-sided $P < 5.89\times10^{-26}$). Out of 579 first-stage associations, 437 (75.4%) have concordant direction of effect ($H_0 = 289.5$; binomial two-sided test $P < 6.84\times10^{-36}$), and 124 SNPs (21.4%) are nominally significant ($P < 0.05$), which is more than would be expected compared to the empirical null distribution (empirical $H_0 = 38.0$ (~6.6%); binomial two-sided test $P = 1.87\times10^{-31}$). Again, these results suggest that the GWAS are not entirely spurious and that our results could be advantageous for identifying genes associated with AUD. We identified four second-stage associations: (1) on chromosome 13 at ~27.9 Mb (rs1333351; second-stage $P = 1.33\times10^{-5}$), (2) on chromosome 11 at ~121.6 Mb (rs7945853; second-stage $P = 2.47\times10^{-5}$), (3) on chromosome 3 at 157.9 Mb (rs1724679; second-stage $P = 2.98\times10^{-5}$), and (4) on chromosome 18 at ~53.0 Mb (rs72926932; second-stage $P = 5.85\times10^{-5}$), which is more than what is expected under the null ($H_0 = 0.05$, binomial test $P = 2.48\times10^{-7}$).

Neither of the four second-stage associations with AUD (nor any SNPs in weak LD, $r^2 > 0.1$) are genome-wide significant in the GWAS in MVP. A SNP in LD (rs9512637, $r^2 = 0.83$) with the first of the second-stage associations, rs1333351, has previously been reported to be associated with a trait cataloged under the label "alcoholism (heaviness of drinking)" at suggestive, but not genome-wide significance (reported $P = 1\times10^{-7}$). Similarly, a SNP in LD (rs9512637, $r^2 = 0.37$) with the third of the second-stage associations, rs1724679, has previously been reported to be associated with drinks per week (reported $P = 6\times10^{-10}$). Neither of the two remaining second-stage associations, rs7945853 and rs72926932, nor any SNPs in weak LD have previously been reported to be associated with any alcohol-related phenotypes, though they previously been reported to be associated with other behavioral traits, such as smoking initiation and number of sexual partners. In summary, the proxy-phenotype analyses with AUD identified two second-stage associations that have previously been identified with respect to alcohol-related phenotypes, and two second-stage associations that are novel to the GWAS literature.

### 5 Polygenic score analyses

#### 5.1 Introduction and summary

In this section, we report a series of analyses that aim to evaluate the out-of-sample accuracy of an externalizing polygenic score, which was computed with weights from the externalizing GWAS. We define accuracy as the incremental $R^2$ attained by adding the polygenic score to a regression model with baseline covariates, in accordance with previous efforts[9,46]. The analyses reported here can be used to assess the potential benefit of applying an externalizing polygenic score for risk stratification or for various empirical research applications[96,97]. For example, an accurate polygenic score can be used to explore the association of the externalizing factor with other traits in cross-trait analyses[98], which we study below, or as a control variable in epidemiological research[97]. To evaluate whether the externalizing polygenic score is robust to bias from population stratification or other sources of bias that can lead to indirect associations between genes and complex traits, we also performed within-family analyses[99].

We generated polygenic scores using three methods, of which two were adjusted for linkage disequilibrium (LD): (1) PRS-CS[100], (2) LDpred[101], as well as (3) unadjusted polygenic scores (henceforth referred to as "classical polygenic scores")[102]. Across the three methods, we only generated scores using SNPs that overlap with the high-quality consensus genotype set defined by the HapMap 3 Consortium[103]. The main polygenic score analyses were performed in the following study cohorts, which were all excluded from the externalizing GWAS to prevent overfitting[104]: (a) Add Health[105,106], (b) COGA[107–109], (c) PNC[110,111], and (d) the UKB Siblings Hold-out cohort (see **Supplementary Information section 2**). We also performed a phenome-wide association study (PheWAS) of electronic health record data in the Vanderbilt University Medical Center biobank (BioVU), using the externalizing polygenic score.

As part of our holistic replication strategy, we first report an investigation of how well the externalizing polygenic score could explain variation in a latent externalizing factor that was created in samples independent from the externalizing GWAS. In Add Health and COGA, we tested a phenotypic externalizing factor that corresponded one-to-one with the seven indicator phenotypes of the preferred Genomic SEM model, by constructing a latent factor from observations of these traits (**Supplementary Table 13**). The externalizing polygenic score was strongly associated with these latent externalizing factor scores and it captured a substantial proportion of their variation (**Supplementary Table 14**, $R^2 \sim$ 9–10%). When we evaluated its robustness in within-family analyses that exploit random genetic differences between sibling and which are therefore immune to population structure and environmental biases that vary between families. The standardized regression coefficient ($\hat{\beta}$) of the score attenuated by 38% and 11.3% in within-family analyses in Add Health and COGA, respectively, while remaining statistically distinguishable from zero (**Supplementary Table 15**, two-sided $P < 0.05$). Considered together, these findings suggest (a) that the externalizing GWAS results capture a substantial part of the variation in externalizing in independent data, (b) population structure, genetic nurture, and other between-family environmental factors that are correlated with genetic variation play a role, but the majority of the signal in our externalizing GWAS results is robust to these factors[99].

Next, we performed a series of exploratory cross-trait analyses with a variety of phenotypes broadly related to externalizing (**Supplementary Tables 16–18**). Whenever possible, we sought to harmonize phenotypes across the study cohorts by using the same or similar observational measures or survey items. A few phenotypes of interest were not measured in all study cohorts, and thus, could only be analyzed in a subset of them. Overall, the externalizing polygenic score was significantly associated with almost all of the tested phenotypes across the behavioral, health, socioeconomic, and criminal justice domains. This finding suggests that genetic liability for externalizing is pervasive and is related to a wide range human behavior. When we focused on within-family analyses in the UKB Siblings Hold-out cohort (**Supplementary Table 19**), we found that the proportion of variance explained by the polygenic score decreased. However, for many of the phenotypes the estimated regression coefficient of the polygenic score remained statistically distinguishable from zero ($P < 0.05$). This result aligns with the aforementioned within-family findings in Add Health and COGA, which suggests that the signal in the externalizing polygenic scores is not just the result of overlooked population stratification while also suggesting that genetic nurture and other between-family environmental factors play an important role in shaping externalizing phenotypes. Overall, the cross-trait associations that we identified indicate that the random allotment of genetic externalizing liability between siblings has predominantly negative consequences for a range of important health and life outcomes.

Finally, we evaluated the association between the externalizing polygenic score and broad-based medical outcomes by conducting a PheWAS in the Vanderbilt University Medical Center Biobank, BioVU (**Supplementary Table 20**). The PheWAS identified a variety of medical conditions that were associated with the externalizing polygenic score. These conditions cover a range of clinical diagnoses, including those related substance use disorders, mental disorders, respiratory disease, type 2 diabetes, and cardiovascular health. The PheWAS findings further emphasize the role played by the genetic liability towards externalizing in shaping negative health outcomes.

The results section below contains further details.

#### 5.2 Methods

##### *5.2.1 Adjustment of GWAS effect sizes for linkage disequilibrium*

We applied two methods to perform LD-adjustment of the effect-size estimates that were used as polygenic score weights, as modeling LD between SNPs is known to increase the signal-to-noise ratio in polygenic scores[96]. The first is a recently developed method called "PRS-CS"[100] (the October 20, 2019 software release), and the second is the often-applied method "LDpred"[101] (because of reported issues with a recent version, we used the older version 0.9.09). As the reference panel for estimating LD, PRS-CS used the 1000 Genomes European reference files distributed with the software and LDpred used the main reference panel (described in **Supplementary Information section 2**). Also, as the PRS-CS method is currently restricted to the ~1.3 million SNPs in the high-quality consensus genotype set defined by the HapMap 3 Consortium[103,112], for comparability, we only generated polygenic scores using HapMap 3 SNPs.

In both cases, we applied the default parameters of the respective software. LDpred has an important tuning parameter that defines the Gaussian mixture weight that represents the fraction of SNPs in the genome that are causal ($p$). As this parameter is not known for most phenotypes, the method developers recommend testing a range of values[101]. However, because we expect externalizing to be highly polygenic and because assuming an infinitesimal model has a closedform solution, we simply chose to adjust the weights using the so-called "LDpred-inf" model. As is recommended[101], we set the LDpred parameter "ld radius" to 340 (this parameter value was determined by dividing the 1,019,937 SNPs that overlap between the externalizing GWAS, the main reference panel, and the HapMap 3 genotype set, by 3,000).

##### *5.2.2 Polygenic scores*

We only computed polygenic scores in individuals of European ancestries[f]. Polygenic scores were computed as the weighted sum of the effect-coded alleles for a given individual *i*:

$$S_i = \sum_{j=1}^{M} \hat{\beta}_j g_{ij}$$

where $S_i$ is the polygenic score, $\hat{\beta}_j$ is the estimated additive effect of the effect-coded allele at SNP *j*, and $g_{ij}$ is the genotype at SNP *j*. For comparability, we computed all scores using the ~1.3 million SNPs in the HapMap 3 consensus genotype set[103]. We performed all analyses below using each of the three scoring methods separately (i.e., never altogether in the same regression model). In our presentation, we highlight the results of the analyses with the PRS-CS polygenic scores, as this method performed consistently best out of the three methods, while the results across the methods were in overall concordance. The complete results are available upon request. The polygenic scores were standardized within each study cohort.

##### *5.2.3 Modeling a latent phenotypic externalizing factor in Add Health and COGA*

We modeled a latent phenotypic externalizing factor in Add Health and COGA that aimed to match the indicator phenotypes of the latent genetic externalizing factor as closely as possible (i.e., ADHD, age at first sexual intercourse, problematic alcohol use, lifetime smoking initiation, general risk tolerance, lifetime cannabis use, and number of sexual partners). In order to generate this latent factor, we fit confirmatory factor models (CFA) in Add Health ($N$ = 15,107) and COGA ($N$ = 16,857) by using all individuals, regardless of ancestry, with non-missing phenotypic data. We estimated all models using Mplus, which allows CFA models to contain indicators of different levels of measurement. To assess model fit and ensure that a single factor specification fit the data adequately, we used a variety of standard fit indices[87] including the comparative fit index (CFI, values closer to 1 indicating better fit), the Tucker-Lewis index (TLI, values closer to 1 indicating better fit), the root mean square error of approximation (RMSEA, values less than .05 indicating good fit), and the standardized root mean squared residual (SRMR, values less than .08 indicating good fit). **Supplementary Table 13** and **Extended Data Fig. 4** present the fit statistics and factor loadings. Further, as it is not straight-forward in a latent variable framework to fit a large number of fixed-effect terms with few observations per term[113], for the within-family analysis described below, we generated observed factor scores from the above CFA model (using the FSCORES function in Mplus). These observed factor scores are simple sums of the seven input phenotypes in the CFA model, weighted by their factor loading.

---

[f]Ancestry assignment was estimated from genetic data. See Braudt and Harris (2018) for full description of Add Health ancestry assignment. In COGA, ancestry was empirically assigned using the 1000 Genomes (phase 3) reference panel (YRI, CEU, JPT and CHB populations) as reference points [164].

Beyond testing the externalizing polygenic score for association with a latent externalizing factor, we also preregistered a variety of exploratory phenotypes for cross-trait analysis. The phenotype definitions are listed in **Supplementary Table 16** and described in detail below. Phenotypes covered a variety of domains thought to be correlates and consequences of externalizing. For illustrative purposes, we categorized these exploratory phenotypes in the following way: (1) substance use initiation; (2) substance use disorders; (3) behavioral problems/disorders; (4) involvement with the criminal justice system; (5) sexual and reproductive health; and (6) socioeconomic outcomes.

##### *5.2.4 Main regression analysis*

In Add Health ($N$ = 5,107), PNC ($N$ = 4,172), and the UKB Siblings Hold-out cohort ($N$ = 39,640), for each tested phenotype ($Y$), we performed two regressions to estimate the accuracy of the polygenic score in explaining phenotypic variation. Specifically, we analyzed regression equations of the following form:

$$\text{Baseline model:} \qquad Y = X\beta + \varepsilon$$

$$\text{Polygenic score model:} \qquad Y = S\gamma + X\beta + \varepsilon$$

where $S$ and $X$ are matrices for the polygenic score and covariates with corresponding vectors of regression coefficients to be estimated, $\gamma$ and $\beta$, respectively. The baseline model included covariates for sex, age, and genetic principal components (PCs), as well as genotyping batch when applicable. The accuracy of the externalizing polygenic score was defined as the difference in $R^2$ (or pseudo-$R^2$) between the two models, a measure that is sometimes called "incremental $R^2$" (or $\Delta R^2$) [46]. In order to demonstrate uncertainty in the incremental $R^2$/pseudo-$R^2$, we estimated 95% confidence intervals using percentile method bootstrapping over 1000 bootstrap samples.

Our choice of statistical model and adjustment of standard errors depended on (1) the distribution of the phenotype and (2) the structure of the data in the study cohort (independent vs. clustered or genetically related observations). In Add Health and PNC, we used ordinary least squares (OLS) for continuous or ordinal outcomes, and logistic regression for binary outcomes. In the UKB Siblings Hold-out cohort we used OLS for continuous and ordinal outcomes, and the linear probability model (LPM) for binary outcomes. A motivation for applying LPM instead of logistic regression in the UKB Siblings Hold-out cohort is given below. With regards to OLS and LPM, we evaluated the traditionally defined coefficient of determination ($R^2$). In the case of logistic regression, we evaluated Nagelkerke's pseudo-$R^2$ [(114)].

In Add Health and PNC, the vast majority of the study participants are unrelated. Therefore, we analyzed one randomly drawn individual from any related pair (pairwise KING coefficient ≥ 0.0442), and thus, did not perform any statistical adjustment for clustering or family structure. In COGA ($N$ = 7,483), which is a family-based cohort study with a variety of different pedigree structures[107–109], to adjust for familial clustering we utilized linear mixed models for continuous and ordinal outcomes (LMM), or generalized linear mixed models (GLMM) with a logistic link function for binary outcomes (except in the within-family analysis, see below). That is, in COGA we estimated the following regression equations:

$$\text{Baseline LMM/GLMM model:} \qquad Y = X\beta + Z\mu + \varepsilon$$

$$\text{Polygenic score LMM/GLMM model:} \qquad Y = S\gamma + X\beta + Z\mu + \varepsilon$$

where we also included a design matrix $Z$ with a binary indicator for each family unit and vector of unobserved random effects $\mu$ (specified as a variance component of the error term)[115]. For the LMM/GLMM models we estimated in COGA, we evaluated a different pseudo-$R^2$ designed specifically for mixed models, described in refs.

We did not adjust the standard errors in Add Health nor PNC, as we analyzed independent observations. Similarly, for COGA, since we used LMM/GLMM, we did not adjust the standard errors either (except in the within-family analysis, see below). To adjust the standard errors for the non-independence of the observations in the UKB Siblings Hold-out cohort, we estimated heteroskedasticity-consistent and cluster-robust standard errors, clustered at the family level.

###### *5.2.5 Within-family analysis in Add Health, COGA, and the UKB Siblings Hold-out*

To evaluate whether the externalizing polygenic score is robust to bias from population stratification or other unaccounted-for between-family differences, we performed within-family analyses in data on full siblings in Add Health, COGA, and the UKB Siblings Hold-out cohort, by comparing the baseline and polygenic score models described above. In this analysis, we studied subsamples of Add Health and COGA, restricted to participants for which we could observe at least one sibling pair in the data (the UKB Siblings Hold-out cohort was already restricted to full siblings). We identified 492 families in Add Health (2–4 siblings in each; $N_{\text{siblings}} = 994$), and 621 families in COGA (2–8 siblings in each family; $N_{\text{siblings}} = 1{,}353$).

In Add Health and COGA, we applied OLS to test the externalizing polygenic score for association with a single outcome: the factor scores of the latent externalizing factor (a continuous variable), while adjusting for family fixed-effects (i.e., family-specific dummy variables)[116,117]. The reason for analyzing factor scores instead of the latent phenotype is that the CFA framework we used above to test the externalizing polygenic score for association is not suitable for modelling a large number of fixed-effect terms with few observations per term [113]. Also, in contrast to the above, the within-family analysis in COGA did not model random family effects ($\mu$) as we instead included fixed effects. Because of the family structure, we analyzed heteroskedasticity-consistent and cluster-robust standard errors, clustered at the family level. Once estimated, we compared the within-family coefficient ($\hat{\beta}$) of the polygenic score with the coefficient estimated in an analogous model without the family fixed-effects.

In the UKB Siblings Hold-out cohort, we performed an analogous within-family analysis with family fixed-effects. In this cohort, we tested the externalizing polygenic score for association with 33 phenotypes in up to 39,640 full siblings, divided across 19,252 family units. Some of the phenotypes are binary, and thus, should arguably be analyzed with e.g., logistic regression. However, estimating logistic regression with a very large number of dummy variables, each with very few observations, can lead to severe bias (i.e., "incidental parameter problem")[115,118,119]. Therefore, we instead applied linear probability models (LPM) for binary outcomes, which has its own drawbacks but is arguably more flexible for modelling a large number of fixed-effect terms[120]. Thus, for comparability, we estimated LPM in both the between- and within-family analysis for binary outcomes this cohort, again with heteroskedasticity-consistent and cluster-robust standard errors.

##### *5.2.6 Phenome-wide association study (PheWAS) in Bio VU*

BioVU is one of the largest biobanks in the United States, consisting of electronic health records from the Vanderbilt University Medical Center on ~250,000 patients spanning 1990 to 2017[121]. A subset of BioVU patients ($N$ = 91,602) have been genotyped as part of various institutional and investigator-initiated projects on the Illumina MEGA$^{EX}$ platform, which contains more than 2 million markers. Quality control (QC) and genotype imputation proceeded as previously described[122]. We computed the externalizing polygenic score in BioVU with the PRS-CS method only, in 66,915 genotyped individuals of European ancestry that are unrelated. Logistic regression was estimated for each of 1,335 case/control medical conditions to estimate the odds of each condition given the externalizing polygenic score, while adjusting for sex, median age of the longitudinal EHR measurements, and 10 genetic PCs. In BioVU, we did not estimate the baseline regression to evaluate $\Delta R^2$. The medical conditions included 42 infectious diseases, 117 neoplasms, 118 endocrine/metabolic diseases, 42 hematopoietic diseases, 63 mental disorders, 68 neurological disorders, 85 sense organ disorders, 145 circulatory system disorders, 76 respiratory diseases, 125 digestive diseases, 120 genitourinary diseases, 31 pregnancy complications, 65 dermatologic disorders, 91 musculoskeletal disorders, 34 congenital anomalies, 37 disease symptoms, and 76 injuries/poisonings. To assign case status, we applied the previously-used requisite of the presence of at least two International Classification of Disease (ICD) codes that mapped to a single so-called "phecode" (Phecode Map 1.2; https://phewascatalog.org/phecodes)[123–125]. We analyzed 1,335 phecodes for which we observed at least 100 cases, and evaluated statistical significance at the Bonferroni-corrected experiment-wide significance threshold ($P < 3.74\times10^{-5}$). This threshold, however, is likely conservative because it assumes independence between phecodes, which is unlikely to hold true due to comorbidity. We ran PheWAS analyses using the PheWAS package v0.12 that is available for the *R* software environment[126].

#### 5.3 Phenotype definitions

##### *5.3.1 Externalizing factor in Add Health*

The phenotypes included for the latent externalizing factor in Add Health match the indicators from the genomic SEM model almost perfectly. *Lifetime smoking initiation* was constructed as a binary measure from the question "Have you ever smoked cigarettes regularly, that is, at least 1 cigarette every day for 30 days?" If individuals indicated yes at any point in the four waves of data, they were coded as being a smoker. Individuals who answered no across all waves were coded as never being a smoker. *Lifetime cannabis use* was constructed in a similar manner to *lifetime smoking initiation*, from the question: "During your life, how many times have you used marijuana?" If participants indicated more than zero at any point in the four waves of data, they were coded as having used cannabis. *Problematic alcohol use* was constructed as an ordinal measure from the lifetime number of symptoms individuals endorsed for DSM-IV alcohol dependence or alcohol abuse criteria (range 0 to 11) at Wave IV when participants received the Composite International Diagnostic Interview-Substance Abuse Module (CIDI-SAM). *ADHD* in Add Health is measured at Wave III using a retrospective scale for ADHD symptoms. The retrospective ADHD scale contains 18 items with responses ranging from "never or rarely" (0) to "very often" (3) and an overall scale ranging from 0 to 54. *Age at first sexual intercourse* was constructed in a stepwise manner. First, we took the earliest reported age at first sexual intercourse across each wave (e.g. if a respondent reported age 14 at Wave I and age 15 at Wave IV, we used the Wave I response as it was closer to the event in time). Next, for those who never reported intercourse, we used Wave IV responses for age at first oral or anal intercourse with the understanding that not all individuals may engage in opposite-sex sexual intercourse. Finally, we coded all individuals with responses below the age of 12 as missing, as this could reflect childhood sexual abuse rather than earlier onset of sexual behavior. *Number of sexual partners* was constructed from the sum of same and/or opposite sex partners an individual reported at the Wave IV interview. Because of the extreme positive skew, we set the maximum number of reported sexual partners at 250 (those reporting > 250 were recoded as 250) as this was the smallest maximum we could set without creating a large number of individuals at the maximum end of the response distribution. Finally, *general risk tolerance* was measured using a single item at Wave IV asking respondents how much do they agreed with the following statement: "I like to take risks." Response ranged from "strongly disagree" (1) to "strongly agree" (5).

##### *5.3.2 Externalizing factor in COGA*

The phenotypes included for the latent externalizing factor in COGA also closely match the indicators from the genomic SEM model. COGA participants received the Semi-Structured Assessment for the Genetics of Alcoholism (SSAGA) [127]. The initial COGA sample (alcohol dependent probands, their family members, and community comparison families) received the SSAGA interview once. A portion of this initial sample received a second SSAGA interview. In addition to the main sample, the COGA Prospective sample (children of the original sample) was followed longitudinally receiving a SSAGA every 2 years (currently 8 waves in total). The SSAGA is a diagnostic interview that covers a variety of psychiatric disorders, including DSM-3R, IV and 5 diagnoses of alcohol, marijuana, cocaine, stimulants, sedatives, opioids, and tobacco use disorders. Additional diagnoses covered by SSAGA include attention-hyperactivity deficit disorder (ADHD), oppositional defiant disorder (ODD), conduct disorder (CD), and antisocial personality disorder (ASPD) (these were not included in the factor analysis but are studied below). Non-diagnostic sections also include demographics and use patterns of alcohol and drugs. Individuals were classified as yes on *lifetime smoking initiation* if they ever answered yes to "Over your lifetime, have you smoked a total of 100 cigarettes (smoked 5 or more packs)?" Individuals who answered no on the initial interview or across each point of data collection (for those who were interviewed more than once) were coded as never being a smoker. For *lifetime cannabis use,* participants were coded as having used cannabis if participants indicated yes to using cannabis at any point. *Problematic alcohol use* was constructed from the number of criteria individuals endorsed for DSM-5 alcohol use disorder (range 0 to 11). Because some COGA participants received more than one interview, we used the maximum value across all waves of participation.

*ADHD* in COGA is measured using DSM-III-R/IV ADHD symptom counts. *Age at first sexual intercourse* was constructed in a manner similar to that in Add Health. We took the earliest reported age at first sexual intercourse across each wave (for those who interviewed more than once) or the reported age from a single item among those who received only one interview. *Number of sexual partners* was constructed from the number of partners an individual reported at the last interview in which they participated. We set the maximum number of reported sexual partners at 300, to overcome similar issues as mentioned in Add Health. Finally, *general risk tolerance* was measured using the Thrill and Adventure Seeking (TAS) subscale of the Sensation

Seeking Scale, or SSS [128]. The SSS provides respondents with 40 questions in which they are given two options to choose one of which best describes how they feel about themselves. For example, items in the TAS asked respondents to choose between options such as: "1) I often wish I could be a mountain climber; or 2) I can't understand people who risk their necks climbing mountains." Individuals who choose the riskier option for each question were coded as 1 and the others were coded 0. The TAS contains ten questions, with total scores ranging from 0 – 10 and higher scores indicating greater tolerance for risky behavior.

##### *5.3.3 Substance use*

Externalizing reflects a broad category of behaviors and psychiatric disorders that reflect a common genetic etiology[11,27,129,130]. Because traits in this category show such strong genetic overlap, we tested whether polygenic scores derived from the genomic SEM model were associated with a variety of substance use phenotypes. For substance use, we created measures of ever use for a variety of substances using each wave/interview in each sample. Respondents were classified as yes on *lifetime smoking initiation* if they responded yes to "Have you ever smoked cigarettes regularly, that is, at least 1 cigarette every day for 30 days?" at any point in Add Health, or yes to "Over your lifetime, have you smoked a total of 100 cigarettes (smoked 5 or more packs)?" at any point in COGA. The definition of *lifetime smoking initiation* and *cigarettes per day* in UKB has been described elsewhere[9]. *Lifetime alcohol use* was coded as yes if participants responded yes to "Have you had a drink of beer, wine, or liquor--not just a sip or taste of someone else's drink--more than 2 or 3 times in your life?" at any wave in Add Health or yes to either "Have you ever had a drink of alcohol?" or "So, you have never had even one full drink of alcohol?" in COGA. In UKB, we used a previously generated measure of *drinks per week* from questions about how often respondents drink alcohol (ranging from 1 "daily or almost daily" to 6 "never") and how much they consumed of various types of alcoholic beverages (wine, beer, spirits, other)[9]. For cannabis use, participants were coded as a yes on *lifetime cannabis use* in Add Health, COGA, and UKB if they responded yes to questions regarding ever using marijuana or hashish across any of the waves/interviews. Participants were classified as a lifetime opioid user (*lifetime opioid use*, COGA only) if they indicated opioids (among a list of many possible substances) for the question "Have you ever used any of these drugs to feel good or high, or to feel more active or alert? Or did you use any prescription drugs when they were not prescribed, or more than prescribed?" Finally, *lifetime other substance use* indicates whether participants have indicated they had ever used a variety of other illicit drugs or prescription medications outside their intended use. In Add Health this included ever using sedatives, tranquilizers, stimulants, painkillers, steroids, cocaine, crystal meth, and/or some other illicit substance. In COGA, the list of other substances included cocaine, stimulants, and/or sedatives.

##### *5.3.4 Substance use disorders*

In addition to substance use, we created measures of substance use disorders and/or problematic use, as the genetic overlap between use and problems is only partial[131]. Both Add Health and COGA contained some form of clinical interview[127,132], while UKB included diagnoses from hospital records and self-reports from interviews. In UKB, we created a binary measure of *problematic alcohol use* from a combination of electronic health records and medical conditions disclosed during an interview, comprised of the following diagnoses: ICD-10 diagnoses (F10.X – Mental and behavioral disorders due to use of alcohol); ICD-9 diagnoses (291.X – Alcoholic psychoses, 303.X – Alcohol dependence syndrome, 305.0X – Nondependent abuse of alcohol), and verbal reports of alcohol dependence. In COGA and Add Health, we constructed measures substance use disorders that correspond to the substances as described above. In Add Health, *alcohol use disorder (AUD) symptoms*, *cannabis use disorder (CUD) symptoms*, and *other substance use disorder (other SUD) symptoms* were measured from the combined criteria counts of DSM-IV dependence and abuse of each of their corresponding substances. The total range of possible criteria ranged from 0 to 11. In COGA, *alcohol use disorder symptoms*, *cannabis use disorder symptoms*, *opioid use disorder (OUD) symptoms*, and *other substance use disorder symptoms* were measured from criteria counts of DSM-5 substance use disorder symptoms. These responses again ranged from 0 to 11. The only differences between combining abuse and dependence from DSM-IV criteria and using the DSM-5 criteria (which mostly reflects the combination of abuse and dependence into a single disorder) are in a single item. DSM-IV abuse contains the criteria of "[i]n the past year, have you more than once gotten arrested, been held at a police station, or had other legal problems because of your drinking?" which was not included in DSM-5. Instead, DSM-5 added, "[i]n the past year, have you wanted to drink so badly you couldn't think of anything else." Finally, in both Add Health and COGA, we used the Fagerstrom test for nicotine dependence (FTND) to assess *nicotine dependence symptoms*. The FTND assesses six criteria and has values ranging from 0 to 10. Overall, these measures of substance use disorders provide good coverage of problematic substance use.

###### *5.3.5 Behavioral problems/disorders*

Each holdout study cohort contains a variety of measures of antisocial and other risky behaviors. Measures for *rule-breaking* were constructed from an index of rule-breaking type behaviors. In Add Health, we took the average of standardized values across the four waves for items comprised of how often individuals reported engaging in the following behaviors over the previous 12 months: painting graffiti or signs on someone else's property or in a public place (adolescence only), deliberately damaging others property, lying to their parents or guardians about where they had been or whom they were with (adolescence only), taking something from a store without paying for it (adolescence only), running away from home (adolescence only), driving a car without its owner's permission (adolescence only), stealing something worth more than $50, stealing something worth less than $50, going into a house or building to steal something, selling marijuana or other drugs, acting loud, rowdy, or unruly in a public place, deliberately writing a bad check (adulthood only), using someone else's credit card, bank card, or automatic teller card without their permission or knowledge (adulthood only), and buying/selling/holding stolen property (adulthood only). Responses ranged from "never" (0) to "5 or more times" (3). In COGA, we used respondent's maximum value from the rule-breaking subscale of the Achenbach Self Report[133] available in the COGA prospective sample only ($N$ = 1,699). The ASR asks respondents to describe their behavior over the past 6 and whether responses are "Not True" (0), "Somewhat or Sometimes True" (1), or "Very True or Often True" (2) and includes items such as "I damage or destroy things belonging to others"; "I break rules at work or elsewhere"; and "I steal." We included previously described measures of *automobile speeding propensity* and *general risk tolerance* from UKB[9].

*Aggression* was measured compiled from a list of aggressive behaviors in both COGA and Add Health. In Add Health respondents whether or not in the past year they had: gotten into a physical fight, pulled a knife or gun on someone, shot or stabbed someone, used/threatened to use a weapon to get something from someone, and/or taken part in a group fight. Because respondents were not asked the specific question of whether or not they had been in a fight, we coded respondents as having been in a fight if they responded yes to either "In the past 12 months, how many times did you take part in a physical fight in which you were so badly injured that you were treated by a doctor or nurse?" or "In the past 12 months, how often did you hurt someone badly enough in a physical fight that he or she needed care from a doctor or nurse?" Responses were coded as yes/no and summed to create a count of 0-5 instances of these events. We used the maximum count across the four waves. In COGA we used the aggression subscale of the ASR (coded the same way as described above). This subscale included statements such as "I get in many fights"; "I physically attack people"; and "I threaten to hurt people."

Finally, we considered several other psychiatric disorders/traits relevant to the externalizing spectrum. *Attention-deficit/hyperactivity disorder (ADHD) diagnosis* is measured from a single item at Wave IV asking respondents if a health care provider ever told them that they had ADHD (Add Health only). *ADHD symptoms* were measured in both Add Health and COGA. In Add Health, participants were asked a series of retrospective questions related to ADHD during the Wave III interview. In COGA, respondents who completed the child version of the SSAGA (CSSAGA) during the initial data collection or were part of the Prospective sample received a section on ADHD. ADHD symptoms were measured as a total criterion count of either DSM-IIIR or DSM-IV criteria for ADHD. In PNC, DSM-IVADHD was measured using a computerized version of the Kiddie-SADS Family Study Interview [134]. In addition to ADHD we also have symptom counts of DSM-IV *conduct disorder* and *oppositional defiant disorder* in PNC. Finally *conduct disorder/antisocial personality disorder (CD/ ASPD) symptoms* (COGA only) were measured from the maximum DSM-IV criteria count of either ASPD (adulthood, 18 or older) or CD (childhood, under age 18) interviews, as ASPD is only assessed in those 18 years and older. Because ASPD and CD have different numbers of criteria, CD criterion counts were proportionally scored in order to create a comparable range (0-7) with ASPD scores (e.g., a participant endorsing 7/15 CD criteria would receive a proportional score of 3.23/7).

##### *5.3.6 Involvement with the criminal justice system*

Involvement with the criminal justice system was captured by a variety of questions included in both Add Health and COGA. *Ever arrested* was measured from the questions "Have you ever been arrested?" at Wave IV in Add Health and "Have you ever been arrested for anything other than moving violations?" from the most recent interview in COGA. In addition, as to ever being arrested, *times arrested* was a count of the number of times one was arrested. *Ever convicted* was measured using "Have you ever been convicted of or pled guilty to any charges other than a minor traffic violation?" in Add Health and "Have you ever been convicted of a felony?" in COGA. Finally, *ever incarcerated* was measured from questions asking "Have you ever spent time in a jail, prison, juvenile detention center or other correctional facility?" in Add Health and "Have you ever spent time in jail for something other than using drugs or alcohol?" in COGA. In each of these measures, individuals were coded as having been arrested, convicted, or incarcerated if they answered yes to any of the corresponding questions in their respective sample.

##### 5.3.7 Sexual and reproductive health behaviors

In addition to problem behaviors and psychiatric conditions, other behaviors related to reproductive and sexual health show genetic overlap with other externalizing traits[38,39]. We therefore investigated the association between polygenic scores for externalizing and a variety of phenotype related to sexual and reproductive health. *Number of sexual partners* was measured using two items in Add Health asking respondents the total number of male and/or female sexual partners with whom they had engaged in any type of sexual activity throughout their lives (retrospectively at Wave IV). In COGA and UKB, we used a single item that asked respondents to list the total number of sexual partners they had been with in their life (from the most recent interview on record in COGA). In Add Health, COGA, and UKB we used the first reported age (across all waves/interviews) at which respondents indicated they first engaged in sexual intercourse to measure *age at first sexual intercourse.* We coded all individuals with a reported age below 12 years old as missing to omit those who were potentially the victims of sexual abuse. Add Health also specifies the type of sexual behavior (vaginal, oral, and/or anal). We started with the first reported age at vaginal intercourse. If individual reported never having vaginal intercourse, we used the first reported age for oral or anal intercourse). *Number of pregnancies* comes from a single item in Wave IV of Add Health that asks respondents to report the number of times an individual has been pregnant (females) or gotten someone else pregnant (males). In COGA, a single question asked to report the number of pregnancies was asked only of female respondents. *Number of live births* comes from a single item asking respondents to report the number of pregnancies that have resulted in a live birth (females only in COGA). In UKB, we broke this measure item into three items: *number of live births* (females only)*, number of children fathered* (males only)*, and number of children ever born* (females and males combined). We also created measures of *age at first birth* (females only) as a continuous measure of age at the first reported birth and *teenage conception* (females only) as a binary measure of whether or not the respondent was a teenager when their first child was born in UKB. Next, in Add Health we measured *lifetime sexually transmitted infection(s) (STI)* from a checklist of STI's in the Wave IV interview which included chlamydia, gonorrhea, trichomoniasis, syphilis, genital herpes, genital warts, hepatitis B, human papilloma virus, pelvic inflammatory disease, cervicitis or mucopurulent cervicitis, urethritis, vaginitis, HIV/AIDS, and/or any other STI. Individuals were coded as having a lifetime STI if they reported yes to any of these diagnoses. In COGA respondents were asked if they had ever been diagnosed with HIV/AIDS or any other STI (Phase IV only; $N$ = 1,774). Finally, *condom use* (Add Health only) was measured from two questions that asked respondents whether they had used condoms and/or female condoms in the previous 12 months.

##### 5.3.8 Socioeconomic outcomes

Because early manifestation of externalizing problems can influence educational trajectories and future socioeconomic attainment[135–137], we examined the association between polygenic scores and outcomes related to these domains. *Educational attainment* was coded as the years of education. In Add Health, respondents were asked the highest grade they had achieved. When indicated, we used the exact number of years to achieve the corresponding grade (e.g. high school graduate = 12). Individuals who reported an educational level of 8$^{th}$ grade or less were coded as 8. In the case where the respondents reported completing some education at a given level and were still enrolled, we took the midpoint for the time between the previous educational milestone and the next (e.g. some college, still enrolled = 14). In COGA and UKB, responses were coded in a similar manner to Add Health. *Personal income* was only available in Add Health and was derived from reported past year earnings in whole dollars. For those who did not know the exact amount they earned, we used the midpoint for categories from a follow-up question that provided ranges of possible dollar amounts. *Household income* was coded as the midpoint of twelve categories ranging from “less than $5000 annually” to “$150,000 or more annually” in Add Health (top category coded as $250,000/year, bottom category coded at $2,500/year), the midpoint of ten categories ranging from “$1-$9,999/year” to “$150,000 or more annually” in COGA (top category coded as $250,000/year, bottom category coded at $5,000/year), and the yearly household income reported in pounds per year in UKB. After recoding income measures into the midpoint of each category (in dollars/pounds per year), we performed a log10 transformation. *Occupational prestige* (Add Health only) was measured using the averaged Hauser and Warren Occupational Income and Occupational Education scales[138]. These scales were created from current or most recent occupation reported at Wave IV using the SOC2000 coding scheme for using pre 2000 occupational codes on post 2000 data[139] which have been used in previously in Add Health[140]. *Full time employment* was measured using a single item (COGA only) that asked participants if they were currently employed in a full-time position. *Fired from work* was measured only of Add Health participants using a single item asking respondents “Thinking back over the period from 2001 to the previous year, how many times have you been fired, let go or laid off from a job?” *Neighborhood disadvantage (ND)* was constructed from the corresponding Wave I (childhood) and Wave IV (adulthood) Census-tract-level data linked to Add Health participants’ home addresses used in prior research[141]. We coded the proportions of: 1) female-headed households, 2) individuals living below the poverty line, 3) individuals receiving public assistance, 4) adults with less than a high school education, and 5) adults who were unemployed into deciles and scored each tract on a scale of 1–10 (corresponding to the decile in which the value fell). Finally, we summed each of these for possible scores ranging from 5-50 at each time point. We also created a measure of *change in neighborhood disadvantage* using the difference of Wave I and Wave IV measures of ND. In UKB, neighborhood conditions were measured using the *Townsend Deprivation Index*[142]; a local social deprivation score based census data (unemployment, non-car ownership, non-home ownership, and household overcrowding), where a higher score implies more social deprivation.

##### *5.3.9 General health and psychological outcomes*

Our UKB holdout sample included several measures related to general health and psychological well-being. For *overall health*, we used a measure of self-rated health ranging from 1) ‘Poor’ to 4) ‘Excellent’, which is widely used as a valid measure of health status [143]. For psychological well-being, we used a total *neuroticism score*, which is the sum of 12 yes/no items including: *irritable person* (“Are you an irritable person”)*, miserableness* (“Do you ever feel 'just miserable' for no reason?”)*, nervous personality* (“Would you call yourself a nervous person?”)*, often feel ‘fed-up’* (“Do you often feel 'fed-up'?”)*, often feel lonely* (“Do you often feel lonely?”)*, often mood swings* (“Does your mood often go up and down?”)*, often troubled by feelings of guilt* (“Are you often troubled by feelings of guilt?”)*, suffers from ‘nerves’* (“Do you suffer from 'nerves'?”)*, tense or ‘high-strung’* (“Would you call yourself tense or 'highly strung'?”)*, worrier* (“Are you a worrier?”)*, worries long after embarrassment* (“Do you worry too long after an embarrassing experience?”)*, feelings easily hurt* (“Are your feelings easily hurt?”). In addition to the total score, we also focused on each individual item that went into the index. Finally, we included a measure of *happiness (subjective well-being)* from a single item asking "In general how happy are you?" ranging from 1) 'Very unhappy' to 6) 'Extremely happy'.

#### 5.4 Results

##### *5.4.1 The latent externalizing phenotype in Add Health and COGA*

We first estimated the fit of the CFA model for the latent externalizing phenotype in Add Health and COGA to ensure that the phenotypes we used as indicators in Genomic SEM also measured a cohesive latent phenotype in these study cohorts. **Supplementary Table 13** and **Extended Data Fig. 4** present the fit statistics and factor loadings. A single factor model, analogous to the latent externalizing factor, demonstrated satisfactory fit in both study cohorts. A similar pattern can be seen when comparing the standardized parameter estimates across the models in each sample. The estimates for *lifetime cannabis use* were strongest in both models ($\beta_{\text{Add Health}}$ = 0.84; $\beta_{\text{COGA}}$ = 0.79), followed closely by *lifetime smoking initiation* ($\beta_{\text{Add Health}}$ = 0.74; $\beta_{\text{COGA}}$ = 0.63). The loadings for *problematic alcohol use* ($\beta_{\text{Add Health}}$ = 0.42; $\beta_{\text{COGA}}$ = 0.65), *number of sexual partners* ($\beta_{\text{Add Health}}$ = 0.40; $\beta_{\text{COGA}}$ = 0.46), and *age at first sexual intercourse* ($\beta_{\text{Add Health}}$ = –0.43; $\beta_{\text{COGA}}$ = –0.48) were similar across the study cohorts. Finally, the loadings for *ADHD symptoms* ($\beta_{\text{Add Health}}$ = 0.27; $\beta_{\text{COGA}}$ = 0.27) and *general risk tolerance* ($\beta_{\text{Add Health}}$ = 0.21; $\beta_{\text{COGA}}$ = 0.21) had the weakest loadings. The overall similarity in both the fit and factor loadings in these diverse samples with the preferred Genomic SEM model specification suggest that the indicators are indeed measuring a consistent latent concept of externalizing both at the phenotypic and genetic level.

We report the results from testing the externalizing polygenic score for association with the latent externalizing factor in **Fig. 2** and **Supplementary Table 14**. Among the three polygenic score methods, the polygenic score derived using PRS-CS had the strongest association ($\hat{\beta}_{Add\ Health}$= 0.328, $\Delta R^2$ = 10.5%; $\hat{\beta}_{COGA}$ = 0.304, $\Delta R^2$ = 8.9%), followed closely by the LDpred derived score ($\hat{\beta}_{Add\ Health}$ = 0.318, $\Delta R^2$ = 9.9%; $\hat{\beta}_{COGA}$ = 0.291, $\Delta R^2$ = 8.3%). In both samples, the classical score (uncorrected for LD) was the least accurate, though its association with the latent externalizing phenotype was still similar to the LD-adjusted methods ($\hat{\beta}_{Add\ Health}$ = 0.253, $\Delta R^2$ = 6.3%, $P = 2.6\times10^{-56}$; $\hat{\beta}_{COGA}$ = 0.273, $\Delta R^2$ = 6.5%, $P = 2.8\times10^{-65}$). These results strongly suggest that our polygenic scores generated with the externalizing GWAS capture a substantial proportion of variation in independent samples. Next, we compared the between- and within-family estimates (**Supplementary Table 15**). The parameter estimates from the within-family model attenuated somewhat (between 11.3–39.3% depending on the study cohort and polygenic score method). At the same time, the within-family estimates remained statistically distinguishable from zero ($P < 0.05$). These results suggest that the externalizing GWAS is relatively robust to bias from population stratification and environmental confounds, while a small-to-moderate proportion of the association observed in the between-family analysis is likely to act via indirect genetic effects, such as genetic nurture.

##### *5.4.2 Results of the exploratory analyses in Add Health and COGA*

We list the availability of the exploratory phenotypes across the study cohorts in **Supplementary Table 16**. The results of the cross-trait exploratory polygenic score analyses in Add Health and COGA are presented in **Supplementary Table 17**. In total, we tested the externalizing polygenic score for association with 34 different exploratory phenotypes, of which 22 were available in both Add Health and COGA. The externalizing polygenic scores was found significantly associated at $P$ less than 0.05 with 31 of the phenotypes, but not with (a) number of live births and (b) number of pregnancies in COGA, nor (c) change in neighborhood disadvantage (childhood to adulthood) in Add Health. The direction of effect was consistent for all phenotypes that were available in both study cohorts. The following sections summarize the results in order of the illustrative categories (1) substance use initiation; (2) substance use disorders; (3) behavioral problems/disorders; (4) involvement with the criminal justice system; (5) sexual and reproductive health; and (6) socioeconomic outcomes.

The externalizing polygenic score was found most strongly associated with phenotypes related to substance use initiation ($\Delta R^2$ ~ 1.09–7.04%). The association with *lifetime smoking initiation* was the strongest among all exploratory phenotypes (Add Health $\Delta R^2$ = 7.04%; COGA $\Delta R^2$ = 5.86%). Notably, the externalizing polygenic score captured more variation in *lifetime smoking initiation* than a polygenic score based on the previously largest genetic study on this phenotype[47], which was used as an indicator GWAS in Genomic SEM. That study reported an incremental pseudo-$R^2$ of 4.2% in Add Health. Thus, this comparison suggests that Genomic SEM was able to increase the accuracy of polygenic scores with respect to an indicator phenotype, which was also the largest by far in terms of sample size. Next, strong associations were identified with *lifetime cannabis use* (Add Health $\Delta R^2$ = 4.91%; COGA $\Delta R^2$ = 2.83%), *lifetime other substance use* (Add Health $\Delta R^2$ = 3.28%; COGA $\Delta R^2$ = 3.95%), and *lifetime opioid use* (only measured in COGA, $\Delta R^2$ = 3.7%). The weakest association among the measures of substance use initiation, was identified with *lifetime alcohol use* (Add Health $\Delta R^2$ = 1.6%; COGA $\Delta R^2$ = 1.09%), which is likely reflection of the ubiquitous nature of this phenotype (95–96% of the Add Health and COGA participants report lifetime alcohol initiation). To put these effect sizes in to context, in Add Health, those with low polygenic scores (–1.5 SD) have a 0.17 projected probability of having ever used other illicit substances. Those with high polygenic scores (+1.5 SD) have projected probability of 0.37, a 2-fold increase in risk.

When we consider the substance use disorder (SUDs) symptoms, we found smaller incremental $R^2$. Nonetheless, we identified positive associations with *alcohol use disorder symptoms* (Add Health $\Delta R^2$ = 0.66%; COGA $\Delta R^2$ = 2.28%), *cannabis use disorder symptoms* (Add Health $\Delta R^2$ = 0.35%; COGA $\Delta R^2$ = 1.17%), *nicotine dependence symptoms* (Add Health $\Delta R^2$ = 0.98%; COGA $\Delta R^2$ = 1.17%), *opioid use disorder symptoms* (only available in COGA, $\Delta R^2$ = 1.71%), and *other substance use disorder symptoms* (Add Health $\Delta R^2$ = 1.49%; COGA $\Delta R^2$ = 1.17%). Overall, our results suggest that the genetic liability for externalizing is strongly associated with increased levels of all the different substance use phenotypes that we tested, while many of these were not indicators in Genomic SEM, which emphasizes that the externalizing GWAS could be leveraged in future studies on various substance use or addiction phenotypes. Again, we see stark differences in those at the extremes of the polygenic score continuum for SUDs in COGA. Those at the top have ~1.5 more projected opioid use disorder symptoms than those at the bottom (+1.5 SD projects 2.69 OUD symptoms; –1.5 SD projects 1.21 OUD symptoms).

We found that polygenic score was associated with a variety of behavioral problems/disorders. First, we identified substantial associations with externalizing psychopathology characterized by disinhibition: DSM-IV *ASPD/CD symptoms* in COGA ($\Delta R^2$ = 2.52%), *ADHD symptoms* (Add Health $\Delta R^2$ = 1.97%; COGA $\Delta R^2$ = 1.77%), and *lifetime ADHD diagnosis* (only available in Add Health $\Delta R^2$ = 1.48%). Next, the polygenic score was associated with both *rule-breaking behavior*

(Add Health $\Delta R^2$ = 1.15%; COGA $\Delta R^2$ = 3.13%) and self-reported *aggression* (Add Health $\Delta R^2$ = 2.31%; COGA $\Delta R^2$ = 1.99%). The stronger effect identified in COGA for *rule-breaking behavior* could reflect differences in the items used to measure these phenotypes across samples. These results suggest that the externalizing GWAS could tag genetic signal with core externalizing traits and psychopathology, and e.g., those at the top of the polygenic distribution (+1.5 SD) in COGA have a projected 4.45 ADHD symptoms while those at the bottom (–1.5 SD) have a projected 2.46 ADHD symptoms.

The externalizing polygenic score was found to be associated with all tested measures of involvement with the criminal justice system, from arrest to incarceration, where higher levels of externalizing liability was associated with greater likelihood of experiencing these conditions. Specifically, the polygenic score was associated with *ever arrested* (Add Health $\Delta R^2$ = 2.45%; COGA $\Delta R^2$ = 3.11%), the *number of times arrested* (Add Health $\Delta R^2$ = 1.54%; COGA $\Delta R^2$ = 0.45%), and with *ever convicted* (Add Health $\Delta R^2$ = 1.39%; COGA $\Delta R^2$ = 4.58%). The difference between Add Health and COGA in terms of $\Delta R^2$ for *ever convicted* likely reflects a difference in severity (see above). Briefly, in Add Health, the question covered multiple levels of conviction, including misdemeanor, felony, and/or being adjudicated as a juvenile, while in COGA the question asked specifically about being convicted of a felony (apparent in the difference in prevalence of being ever convicted, which is 13.78% and 3.31% in Add Health and COGA, respectively). The polygenic score was associated with *ever incarcerated* (Add Health $\Delta R^2$ = 2.45%; COGA $\Delta R^2$ = 3.10%). In Add Health, these effect sizes translate into those with low polygenic scores (–1.5 SD) having a 0.18 projected probability of ever being arrested and those with high polygenic scores (+1.5 SD) having a projected probability of 0.36 of ever being arrested. (We remind the reader that these comparisons were only performed among individuals of European ancestry.)

The fifth set of phenotypes we examined in our exploratory polygenic score analyses fall into the domain of sexual and reproductive health. Because previous work shows genetic overlap between many of the traits on the externalizing spectrum and sexual behaviors[38,39], we expected the polygenic scores to be associated with multiple phenotypes in this category. The two phenotypes for which we estimated the strongest associations, *age at first sexual intercourse* (Add Health $\Delta R^2$ = 4.57%; COGA $\Delta R^2$ = 2.87%) and *number of sexual partners* (Add Health $\Delta R^2$ = 1.49%; COGA $\Delta R^2$ = 1.06%) were indicators in the preferred Genomic SEM model. The polygenic score was also associated with greater *number of pregnancies* (Add Health $\Delta R^2$ = 1.53%; COGA $\Delta R^2$ = 0.37%, $P$ = 0.73), and greater *number of live births* (Add Health $\Delta R^2$ = 0.29%; COGA $\Delta R^2$ = 0.0%, $P$ = 0.86), but only in Add Health. The difference in effect sizes across the study cohorts could reflect the fact that in COGA, questions related to pregnancy and births were only asked to female participants, whereas in Add Health it was asked to all participants (for males, reflecting pregnancies of a partner). The polygenic score had a significant, but small association with having a *lifetime sexually transmitted infection* (Add Health $\Delta R^2$ = 0.6%; COGA $\Delta R^2$ = 0.91%). Finally, the polygenic score had a weak but significant association with reporting less *condom use* in the previous 12 months (Add Health $\Delta R^2$ = 0.17%; not measured in COGA). The projected age at first sexual intercourse in Add Health is 1.86 years younger for those at the top of the polygenic distribution (+1.5 SD = 15.67) relative to those at the bottom (–1.5 SD = 17.53). Thus, the genetic liability for externalizing was associated with what could arguably be considered riskier sexual and reproductive behavior.

The final set of exploratory phenotypes we tested for association in Add Health and COGA were categorized as socioeconomic measures. As childhood and adolescent externalizing is known to be associated with a lower educational trajectory and reduced future social mobility[135–137], we expected the externalizing polygenic score to be negatively associated with measures in this category. The strongest association was found between the polygenic score and lower *educational attainment* (Add Health $\Delta R^2$ = 3.03%; COGA $\Delta R^2$ = 1.64%), followed by less *occupational prestige* (Add Health $\Delta R^2$ = 1.91%; not measured in COGA), lower *personal income* (Add Health $\Delta R^2$ = 1.00%), and lower *household income* (Add Health $\Delta R^2$ = 0.97%; COGA $\Delta R^2$ = 0.90%). In addition, the polygenic score was associated with other labor market measures, including reporting an increased *number of times fired* (Add Health $\Delta R^2$ = 1.24%) and *fulltime employed* to a less extent (COGA $\Delta R^2$ = 0.12%; not measured in Add Health), an index of neighborhood disadvantage[141] in both childhood/adolescence (*childhood neighborhood disadvantage*; Add Health $\Delta R^2$ = 0.7%; not measured in COGA) and adulthood (*adult neighborhood disadvantage*; Add Health $\Delta R^2$ = 0.51%; not measured in COGA). The differences in projected household income between the top and bottom of the polygenic score distribution is approximately \$10,000 (+1.5 SD = \$44,640; –1.5 SD = \$33,737). Overall, these findings align with the literature on the relationship between externalizing and socioeconomic status, suggesting that externalizing is generally associated with lower socioeconomic status.

##### *5.4.3 Results of the exploratory analyses in PNC*

We considered symptom counts of three psychiatric phenotypes related to behavioral problems/disorders in PNC: DSM-IV *attention-deficit/hyperactivity disorder* (ADHD), *oppositional defiant disorder* (ODD), and *conduct disorder* (CD), which are all considered central diagnoses with respect to externalizing psychopathology. The results are reported in **Supplementary Table 18**. The externalizing polygenic score was significantly associated with an increased number of symptoms for each of the three disorders, and the score explained a modest proportion of the variance in each measure (ADHD $\Delta R^2$ = 1.19%; CD $\Delta R^2$ = 3.51%; ODD $\Delta R^2$ = 1.92%). The results in PNC further increase our confidence in that the externalizing GWAS captures genetic signal of important not only for externalizing behaviors but also externalizing psychopathology.

##### *5.4.4 Results of the analyses in the UK Biobank Siblings Hold-out cohort*

Results for the analyses in the UKB Siblings Hold-out cohort are reported in **Supplementary Table 19**. In the between-family models, the externalizing polygenic score was found associated with 30 out of 33 tested phenotypes, with the exceptions of (1) *cigarettes per day*, (2) *happiness (subjective well-being)*, and (3) *suffers from "nerves"*. Across the outcomes, greater genetic liability for externalizing was associated with more risky health behaviors (drinking, smoking, substance use initiation, etc.), lower socioeconomic status, and poorer mental, physical, and sexual health. The incremental $R^2$ for these between-family associations ranged from tiny (*feelings easily hurt*, $\Delta R^2$ = 0.02%) to modest (*lifetime smoking initiation*, $\Delta R^2$ = 3.89%). The generally greater incremental $R^2$ that we identified in Add Health, COGA, and PNC compared to the UKB Siblings Hold-out cohort, is likely the results of the much richer and detailed phenotypic data available in the former three study cohorts.

Of the 30 significant associations, 21 remain statistically distinguishable from zero (two-sided test $P < 0.05$) in the within-family analyses. The phenotypes that did not remain associated in the within-family model included *children fathered (males with children)*, *feelings easily hurt*, *fluid intelligence*, *household income*, *live births (females)*, *neuroticism score*, *often feels lonely*, *often troubled by feelings of guilt*, *problematic alcohol use*, and *tense or highly strung*. While the polygenic score remained associated with the majority of the phenotypes in the within-family analysis, when comparing the ratio of parameter estimates from the between and within models, we see that the between-family estimates were on average 1.49 times larger than the within family estimates ($Beta/Beta_{WF}$ = 0.254–4.856). The ratio in these estimates suggests that the association between our polygenic score and many of the phenotypes is due, in part, to between-family confounding by phenomena like genetic nurture or population stratification.

##### 5.4.5 Results of the PheWAS in BioVU

In BioVU, we tested 1,335 medical outcomes for association with the externalizing polygenic score, of which 84 were found significantly associated at Bonferroni-corrected experiment-wide significance ($P < 3.27\times10^{-5}$). We note that Bonferroni correction is overly conservative here because it ignores comorbidities between medical outcomes. The results are displayed in **Fig. 3** and **Supplementary Table 20**. As expected, many associations were identified in the mental disorder category ($k = 14$). Noteworthy associations in that group are tobacco use disorder ($N_{cases}$ = 6,155, OR = 1.31, $P = 1.65\times10^{-82}$), substance addiction and disorders ($N_{cases}$ = 2,062, OR = 1.30, $P = 2.46\times10^{-32}$), alcoholism ($N_{cases}$ = 1,020, OR = 1.29, $P = 4.5\times10^{-15}$), mood disorders ($N_{cases}$ = 9,588, OR = 1.10, $P = 1.03\times10^{-14}$) and suicidal ideation or attempt ($N_{cases}$ = 689, OR = 1.20, $P = 3.30\times10^{-6}$), as well as bipolar disorder ($N_{cases}$ = 1,565, OR = 1.18, $P = 2.13\times10^{-10}$) and major depressive disorder ($N_{cases}$ = 3,990, OR = 1.101, $P = 8.79\times10^{-9}$). The score was not experiment-wide significantly associated with either ADHD (OR = 1.13, $P = 5.91\times10^{-5}$) or conduct disorders (OR = 1.15, $P = 3.7\times10^{-3}$), likely because of the relatively limited number of cases $N_{cases}$ = 1,027 and $N_{cases}$ = 426, respectively. However, both ADHD and conduct disorders were significant at the more liberal false-discovery rate of 0.05. Overall, we again find strong links with substance use disorders. Further, these findings are in concordance with the genetic correlations we estimated, which suggest that the genetic liability for externalizing may be partly shared with other major mental disorders.

Next, the score was also associated with a range of different medical outcomes in various disease categories, including the circulatory system ($k = 17$), such as ischemic heart disease score ($N_{cases}$ = 9,991, OR = 1.10, $P = 3.66\times10^{-12}$); respiratory diseases ($k = 17$), such as chronic airway obstruction ($N_{cases}$ = 4,436, OR = 1.17, $P = 2.74\times10^{-22}$); infectious diseases ($k = 7$), such as viral hepatitis C and HIV disease ($N_{cases}$ = 1,195, OR = 1.39, $P = 1.57\times10^{-28}$; and $N_{cases}$ = 677, OR = 1.21, $P = 2.11\times10^{-6}$, respectively); endocrine/metabolic conditions ($k = 7$), such as type 2 diabetes ($N_{cases}$ = 8,959, OR = 1.05, $P = 1.73\times10^{-5}$, respectively); digestive diseases ($k = 6$), including cirrhosis of liver (e.g., $N_{cases}$ = 1,928, OR = 1.21, $P = 1.87\times10^{-15}$); neurological ($k = 2$), such as chronic pain ($N_{cases}$ = 3,172, OR = 1.15, $P = 2.09\times10^{-13}$); neoplasms ($k = 5$), including lung cancer ($N_{cases}$ = 2,260, OR = 1.14, $P = 9.05\times10^{-10}$); and other categories including injuries and poisonings ($k = 2$), genitourinary ($k = 4$), hematopoietic ($k = 4$), musculoskeletal ($k = 3$), dermatologic ($k = 1$), sense organs ($k = 4$), and symptoms ($k = 1$).

It is likely that the externalizing polygenic score is associated with many medical outcomes via a range of risky health behaviors. In particular, increased drinking and alcohol use disorders may explain the association with e.g., liver cirrhosis and injuries, while lifetime smoking initiation likely explains the associations with various neoplasms and respiratory diseases known to be caused by tobacco smoking, such as lung cancer, emphysema, and chronic airway obstruction. Notably, we identified associations with Viral hepatitis C ($N_{cases}$ = 1,195, OR = 1.39, $P$ = $1.57\times10^{-28}$) and HIV diagnosis ($N_{cases}$ = 677, OR = 1.21, $P$ = $2.11\times10^{-6}$), which could be due to riskier sexual behaviors or unsafe substance use practices, such as needle sharing. These findings align with both those in Add Health and COGA on number of sexual partners and age at first sexual intercourse, and in COGA on lifetime opioid use and OUD symptoms. In conclusion, these results display the importance of considering the influence of externalizing liability in shaping a range of negative health outcomes and substance use.

### 6 Bioannotation

In this section, we describe analyses investigating the biological function across all SNPs in the externalizing GWAS, as well as of the 579 jointly associated lead SNPs that we identified (**Supplementary Information section 3**), by using a variety of bioinformatics tools. Specifically, we applied functional genomic tools to annotate and prioritize putative regulatory variants, including functional annotations (i.e., CADD scores to identify highly deleterious SNPs), mapping annotations (i.e., eQTL SNP-gene expression association), gene-, and transcriptome-based analyses (MAGMA, S-PrediXcan). Details of each method are presented below, and the results are reported in **Supplementary Tables 9–10** and **21–30**, and displayed in **Extended Data Figs. 5–8**.

#### 6.1 Methods

##### *6.1.1 Functional mapping and annotation with FUMA*

We used the method "functional mapping and annotation of genetic associations" (FUMA v1.3.5e)[18] to study the functional consequences of the 579 jointly associated lead SNPs (the results are reported in **Supplementary Table 9**), which included ANNOVAR categories (*i.e.*, the functional consequence of SNPs on genes), Combined Annotation Dependent Depletion (CADD) scores (*i.e.*, a measure of how deleterious a SNP is; greater than 12.37 is the suggested threshold to classify a SNP as deleterious), RegulomeDB scores (*i.e.*, a categorical score from 1a to 7 with 1a corresponding to the most biological evidence that the SNP is a regulatory element), mapping to expression quantitative trait loci (eQTLs are SNPs that influence gene expression; herein we focused on brain tissue eQTLs), and chromatin states (characterization of chromatin state; values range from 1 to 15 with values 1 to 7 referring to an open chromatin state). The sources of the external reference data used in these analyses are fully described in ref.[18].

With FUMA, we also performed lookups in the GWAS Catalog (version e96 2019-05-03, data analysis performed on 2020-03-25) to investigate whether the loci identified in the externalizing GWAS have previously been reported as associated with other traits at suggestive significance (two-sided $P < 1\times10^{-5}$). The GWAS Catalog compiles results from all published GWAS[48]. We extracted information from the GWAS Catalog for any of the 579 jointly associated lead SNPs (as well as for any SNPs in LD, $r^2 > 0.1$) that were reported in the catalog (the results are reported in **Supplementary Table 10**).

##### *6.1.2 Gene-based, gene-set, and gene-property analyses with MAGMA*

We performed competitive gene-based association analyses using the genome-wide summary statistics from the externalizing GWAS by applying the method "multi-marker analysis of genomic annotation" (MAGMA v1.8)[18,19]. First, we assigned SNPs to genes based on physical position (gene-based analysis). SNPs were mapped to 18,235 protein-coding genes from Ensembl build 85. This approach uses multiple regression methods to account for LD between SNPs. All variants within all protein-coding genes were tested, using default settings, with LD structure estimated using the 1000 Genomes European sample as a reference. We evaluated

Bonferroni-corrected significance, adjusted for testing 18,235 genes (one-sided $P < 2.74\times10^{-6}$). The results are reported in **Supplementary Table 21**.

Next, to study the relationship between the externalizing GWAS and sets of genes that share specific functional or biological characteristics, we performed a MAGMA gene-set analysis (the results are reported in **Supplementary Table 22**). We used 15,481 curated gene sets and Gene Ontology (GO) terms obtained from the Molecular Signatures Database (MsigDB v7.0, https://www.gsea-msigdb.org/gsea/msigdb/index.jsp)[144], which characterize the biological processes, molecular function and cellular component of individual gene products. We evaluated Bonferroni-corrected significance, adjusted for testing 15,481 gene sets (one-sided $P < 3.23\times10^{-6}$).

Lastly, we performed a gene property analysis to test the relationships between 54 tissue-specific gene expression profiles and gene associations (the results are reported in **Supplementary Table 23**). We performed this analysis using the average expression of genes per tissue type as a gene covariate. Gene expression values were $\log_2$ transformed average RPKM (Reads Per Kilobase Million) per tissue type (after replacing RPKM > 50 with 50) based on GTEx RNA-seq data. We applied Bonferroni correction (one-sided $P < 9.26\times10^{-4}$) to correct for testing 54 gene expression profiles. In addition, to examine the relationship between the externalizing GWAS and general developmental stages, we performed a MAGMA gene-set analysis using 11 developmental stages from brain samples obtained from BrainSpain (http://www.brainspan.org/static/download). The results of this analysis are reported in **Supplementary Table 24**. Further details on these methods are described in refs.[18,19].

###### ***6.1.3 Gene-based analysis using chromatin interaction profiles from human brain tissue with H-MAGMA***

We used an extension of MAGMA v1.8, "Hi-C coupled MAGMA" or "H-MAGMA" [20], to assign non-coding (intergenic and intronic) SNPs to cognate genes based on their chromatin interactions. Exonic and promoter SNPs were assigned to genes based on physical position. We used four Hi-C datasets derived from adult brain[145], fetal brain[146], and iPSC derived neurons and astrocytes[147] (all available for download: https://github.com/thewonlab/H-MAGMA). The results are reported in **Supplementary Tables 25–28**. We evaluated Bonferroni corrected *P*-value thresholds, adjusted for multiple testing within each analysis (one-sided $P < 9.84\times10^{-7}$, $P < 9.86\times10^{-7}$, $P < 9.84\times10^{-7}$, and $P < 9.83\times10^{-7}$, respectively).

###### ***6.1.4 Gene-based association using transcriptomic data with S-PrediXcan***

We used S-PrediXcan v0.6.2[22] to analyze gene expression levels in multiple brain tissues, and to test whether the gene expression correlated with the genetic liability of externalizing. The results are reported in **Supplementary Table 29**. We used pre-computed tissue weights from the Genotype-Tissue Expression (GTEx, v8) project database (https://www.gtexportal.org/) as the reference transcriptome dataset[148]. As input data, we used the summary statistics for the externalizing GWAS, transcriptome tissue data, and covariance matrices of the SNPs within each gene model (based on HapMap SNP set; available to download at the PredictDB Data Repository, http://predictdb.org) from 13 brain tissues: anterior cingulate cortex, amygdala, caudate basal ganglia, cerebellar hemisphere, cerebellum, cortex, frontal cortex, hippocampus, hypothalamus, nucleus accumbens basal ganglia, putamen basal ganglia, spinal cord and substantia nigra. We used a transcriptome-wide significance threshold of $P < 2.73\times10^{-7}$, which is the Bonferroni-corrected threshold when adjusting for 13 tissues times 14,095 tested genes (183,235 gene-tissue pairs).

#### 6.2 Results

##### *6.2.1 Results from the functional mapping and annotation with FUMA*

Out of the 579 jointly associated lead SNPs, 233 were intronic, 13 are exonic, 5 were in the 3' UTR, and 3 were in the 5' UTR; 106 variants were found significantly associated with an eQTL previously linked to expression in brain tissue; 60 were annotated with CADD scores greater than 12.37, indicating high probability of being deleterious (**Supplementary Table 9**). Several of the variants with CADD scores greater than 12.37 were located within genes previously related to drug use and risk tolerance, such as Cell Adhesion Molecule 2 (*CADM2*)[9,59] (strongest signal rs993137, beta = 0.02, $P = 4.61\times10^{-53}$), Microtubule Associated Protein Tau (*MAPT*)/Corticotropin Releasing Hormone Receptor 1 (*CRHR1*)[58,149] (rs2258689, beta = –0.01, $P = 1.68\times10^{-8}$); brain volume, such as Zic Family Member 4 (*ZIC4*)[150] (rs2279829, beta = 0.01, $P = 2.88\times10^{-18}$). Other genes are less prominent in the previous literature, such as the Calcium Voltage-Gated Channel Subunit Alpha1 D (*CACNA1D*) gene (rs312480, beta = –0.01, $P = 2.14\times10^{-10}$), or the gene Protein Kinase C And Kinase Substrate in Neurons 3 (*PACSIN3*; rs901750, beta = –0.01, $P = 1.21\times10^{-10}$). Interestingly, overexpression of *PACSIN3* impairs internalization of Solute Carrier Family 2, Facilitated Glucose Transporter Member 1 (*SLC2A1*)/Glucose Transporter 1 (*GLUT1*). In the brain, *GLUT1* protein is involved in moving glucose, the brain's major energy source, across the blood-brain barrier. Of note, the S-PrediXcan analysis, reported next, also identified that more expression of *PACSIN3* in the nucleus accumbens was significantly associated with externalizing ($P = 1.46\times10^{-6}$). In summary, and in alignment with other GWAS, most of the loci we identified in the externalizing GWAS are located outside of genes or are eQTLs, and thus, are likely to affect the phenotype by altering the amount or timing of protein production[151]. Notably, a substantial subset (~10%) of the identified loci had high CADD scores and these are therefore likely to directly change the type or structure of the gene products.

##### *6.2.2 Results from the analyses with MAGMA, H-MAGMA, and S-PrediXcan*

In order to identify associations at the level of genes rather than SNPs, we performed two types of gene-based analyses based on GWAS summary statistics: (1) MAGMA, which aggregates SNP effects at the gene level using positional annotations, and (2) S-PrediXcan, which uses reference data on expression quantitative-trait loci (eQTL) annotations to assign SNPs to genes. The summary statistics for the externalizing GWAS was the input used to compute gene-based $P$ values. In the MAGMA analysis, a total of 928 genes were found associated at a Bonferroni-corrected significance (one-sided $P < 2.74\times10^{-6}$) (**Extended Data Fig. 5** and **Supplementary Table 21**), of which 244 have one or more genome-wide significant SNPs from the externalizing GWAS within their gene breakpoints (**Supplementary Table 9**). Next, the MAGMA gene-property analysis identified that the externalizing GWAS was significantly ($P < 9.26\times10^{-4}$)

enriched for association in multiple brain tissues, including the cerebellar hemisphere ($P = 1.10\times10^{-22}$), cerebellum ($P = 1.54\times10^{-22}$) and frontal cortex BA9 ($P = 2.66\times10^{-19}$), as well as and pituitary gland tissues ($P = 2.80\times10^{-6}$). (**Extended Data Fig. 6** and **Supplementary Table 23**). Interestingly, this "tissue-wide" analysis did not suggest any other tissues than those located in the brain. Intriguingly, out of 11 developmental stages, we found that genes were primarily expressed in the brain prenatally (**Extended Data Fig. 7** and **Supplementary Table 24**). Additionally, the MAGMA gene-set analysis identified that sets relating to synaptic plasticity were significantly associated with externalizing. Out of the 15 significant gene-sets ($P < 3.23\times10^{-6}$), 5 gene-sets involved neuron development/differentiation (e.g. neuron differentiation, $P = 1.16\times10^{-7}$), and 4 gene-sets involved synapses (e.g. synapse, $P = 3.46\times10^{-8}$) (**Supplementary Table 22**).

By analyzing gene regulatory relationships using H-MAGMA, we identified significant associations in adult brain tissue (2,033 genes), fetal brain tissue (1,953 genes), iPSC-derived astrocytes (1,974 genes), and iPSC-derived neurons (1,973 genes; **Supplementary Tables 25–28**). Using S-PrediXcan, we identified changes in predicted gene expression from 348 genes (of which 156 were also significant in the MAGMA analysis) in multiple brain regions as significantly associated with externalizing, at a Bonferroni-corrected significance threshold of $P < 2.73\times10^{-7}$ (**Supplementary Table 29**).

We identified 34 genes that were consistently implicated by all methods we applied, as these have jointly associated SNPs within their breakpoints, and were consistently associated across the MAGMA, H-MAGMA (adult tissue) and S-PrediXcan analyses; these include *CADM2*, *PACSIN3*, *ZIC4*, *MAPT*, *GABRA2*. The full list of overlapping and unique genes is shown in the **Supplementary Table 30**. The number of implicated genes that overlap across the methods is displayed in a Venn diagram in **Extended Data Fig. 8**. In summary, the results of the analyses we performed with MAGMA, S-PrediXcan, and H-MAGMA all suggest that the externalizing GWAS is enriched for association with genetic variants that are involved in brain development, function, and structure.

##### *6.2.3 Results from the GWAS Catalog lookup*

We report the results of the lookups of the 579 jointly associated SNPs (and any SNPs in LD, $r^2 > 0.1$) in **Supplementary Table 10**. In summary, we found that 538 of the SNPs or their correlates have previously been reported in the GWAS Catalog at suggestive significance ($P < 1\times10^{-5}$). Thus, we were able to identify 41 novel genetic loci that have previously not been reported for association with any trait in the GWAS literature. Virtually all of the reports we found overlap with multiple other phenotypes. Most of the previously reported associations are with traits related to the externalizing spectrum, including risk tolerance[9], smoking[9,47], alcohol consumption[9,47,152], and cannabis use[56,153], or other behavioral or mental traits. At the same time, we also found overlap with many seemingly unrelated traits, such as heel bone mineral density[154], acne[155], or blood protein levels[156]. Overall, these findings align with the genetic correlations we estimated, alongside with the known widespread pleiotropy in the human genome[157], which together suggest great genetic overlap between the externalizing factor with a range of different complex traits.

#### 6.3 Discussion

Broadly, the results of the performed bioinformatic analyses converge to reveal an abundance of pleiotropic genes that are known to play a major role in neurodevelopment. Of the 60 jointly associated SNPs with CADD scores greater than 12.37, gene- or transcriptome-based analyses identified genes that have previously been implicated in multiple studies, such as the brain-derived neurotrophic factor (*BDNF*, $P = 7.85\times10^{-16}$), regulator of brain plasticity; RNA Binding Fox-1 Homolog 1 (*RBFOX1*, $P = 6.40\times10^{-17}$) and Netrin 1 Receptor genes (*DCC*, $P = 3.58\times10^{-28}$), which were also significant in a recent cross-disorder GWAS meta-analysis by the Psychiatrics Genomics Consortium[85], which appear to play a role in neuronal development. Gene- and transcriptome-based analyses also identified previously suggested genes that have been shown to be extremely pleiotropic, including Paired Basic Amino Acid Cleaving Enzyme (*FURIN;* involved in at least 40 other GWAS studies, including multiple psychiatric disorders[158] and the recent cross disorder[85], cardiovascular disease[159,160]), Potassium Inwardly Rectifying Channel Subfamily J Member 3 (*KCNJ3*; involved in smoking, alcohol consumption, cognitive performance, among others[47,161]), Gamma-Aminobutyric Acid Type A Receptor Subunit Alpha 2 (*GABRA2*; the major inhibitory neurotransmitter in the mammalian brain, suggested for virtually all major psychiatric disorders[9,162]), and Forkhead Box P2 (*FOXP2*)[9,163].

### Supplementary Notes

#### 8 Author contributions

Danielle Dick and Philipp Koellinger conceived the study. The study protocol was developed by Danielle Dick, Paige Harden, Richard Karlsson Linnér, Philipp Koellinger, Travis Mallard, and Abraham Palmer. Danielle Dick, Paige Harden, Philipp Koellinger, and Abraham Palmer jointly oversaw the study. Danielle Dick and Richard Karlsson Linnér led the writing of the manuscript, with substantive contributions to the writing from Paige Harden, Philipp Koellinger, and Abraham Palmer.

Richard Karlsson Linnér and Travis Mallard were the lead analysts, responsible for conducting genome-wide association studies, quality control, meta-analysis, genetic correlations, and multivariate analyses with Genomic SEM, among other analyses reported in **Supplementary Information sections 2–3**, with assistance from Andrew Grotzinger. Richard Karlsson Linnér performed the proxy-phenotype analyses in **Supplementary Information section 4**. Peter Barr led the polygenic score analyses in **Supplementary Information section 5**, and Richard Karlsson Linnér and Travis Mallard contributed to those analyses. Sandra Sanchez-Roige performed the PheWAS in BioVU. Sandra Sanchez-Roige led the bioinformatics analyses in **Supplementary Information section 6**, and Richard Karlsson Linnér contributed to those analyses.

Peter Barr, Richard Karlsson Linnér, Travis Mallard, and Sandra Sanchez-Roige prepared the tables and figures, with assistance from Morgan Driver, James Madole, and Holly Poore.

Jorim Tielbeek, Emma Johnson, Mengzhen Liu, Hang Zhou, Rachel Kember, and Joëlle Pasman prepared cohort-level GWAS meta-analyses. These analyses were supervised by Karin Verweij, Dajiang Liu, Scott Vrieze, Henry Kranzler, and Joel Gelernter. Kathleen Mullan Harris assisted analyses performed in the AddHealth study cohort.

Andrew Grotzinger, Elliot Tucker-Drob, and Irwin Waldman provided helpful advice and feedback on various aspects of the study design.

All authors contributed to and critically reviewed the manuscript. Richard Karlsson Linnér, Travis Mallard, Peter Barr, and Sandra Sanchez-Roige made especially major contributions to the writing and editing.

##### 8.1 Additional acknowledgments

###### *8.1.1 Investigator acknowledgments and competing interests*

Danielle M. Dick acknowledges funding provided by the National Institute of Alcohol Abuse and Alcoholism of the National Institutes of Health under award numbers R01AA015146, K02AA018755, U10AA008401, and P50AA0022537.

K. Paige Harden acknowledges funding provided the Eunice Kennedy Shriver National Institute of Child Health and Human Development of the National Institutes of Health under award numbers HD092548 and HD083613; and funding provided by the Jacobs Foundation.

Abraham A. Palmer funding provided by the National Institute of Alcohol Abuse and Alcoholism and the National Institute of Drug abuse of the National Institutes of Health under award numbers AA026281 and P50DA037844; and funding from the California Tobacco-Related Disease Research Program (TRDRP) under grant numbers 28IR-0070 and T29KT0526.

Philipp D. Koellinger acknowledges financial support from the European Research Council (consolidator grant 647648) and from the Netherlands Organization for Scientific Research (NWO project 17184 for access to the Cartesius supercomputer hosted by SURFsara on which the majority of statistical analyses were carried out).

Henry R. Kranzler acknowledges support by Million Veteran Program grant I01 BX003341 from the U.S. Department of Veterans Affairs Biomedical Laboratory Research and Development Service. Dr. Kranzler is a member of the American Society of Clinical Psychopharmacology's Alcohol Clinical Trials Initiative, which was supported in the last three years by AbbVie, Alkermes, Ethypharm, Indivior, Lilly, Lundbeck, Otsuka, Pfizer, Arbor, and Amygdala Neurosciences.

Henry R. Kranzler and Joel Gelernter are named as inventors on PCT patent application #15/878,640 entitled: "Genotype-guided dosing of opioid agonists," filed January 24, 2018.

Joel Gelernter reported receiving grants from US Department of Veterans Affairs and the National Institutes of Health (NIDA, NIAAA, NIMH) during the conduct of the study; and did paid editorial work for the journal Complex Psychiatry.

Karin J. H. Verweij is supported by the Foundation Volksbond Rotterdam and by a grant from Amsterdam Neuroscience.

Sandra Sanchez-Roige was supported by the Frontiers of Innovation Scholars Program (FISP; #3-P3029), the Interdisciplinary Research Fellowship in NeuroAIDS (IRFN; MH081482), a NARSAD Young Investigator Award from the Brain and Behavior Foundation under grant number 27676; funds from the California Tobacco-Related Disease Research Program (TRDRP) under grant numbers 28IR-0070 and T29KT0526, and a pilot award from P50DA037844.

#### 8.2 Cohort acknowledgments

##### *8.2.1 23andMe*

We would like to thank the research participants and employees of 23andMe for making this work possible. 23andMe research participants provided informed consent and participated in the research online, under a protocol approved by the AAHRPP-accredited institutional review board, Ethical and Independent Review Services (E&I Review). Participant data are shared according to community standards that have been developed to protect against breaches of privacy. Currently, these standards allow for the sharing of summary statistics for at most 10,000 SNPs. The full set of externalizing GWAS summary statistics can be made available to qualified investigators who enter into an agreement with 23andMe that protects participant confidentiality. Once the request has been approved by 23andMe, a representative of the Externalizing

Consortium can share the full set of summary statistics. Summary statistics from the *EXT* analyses can be obtained by following the procedures detailed at:

https://externalizing.org/request-data/

##### *8.2.2 Add Health*

This research uses data from Add Health, a program project directed by Kathleen Mullan Harris and designed by J. Richard Udry, Peter S. Bearman, and Kathleen Mullan Harris at the University of North Carolina at Chapel Hill, and funded by grant P01 HD31921 from *Eunice Kennedy Shriver* National Institute of Child Health and Human Development (NICHD), with cooperative funding from 23 other federal agencies and foundations. Add Health GWAS data were funded by NICHD Grants R01 HD073342 (Harris) and R01 HD060726 (Harris, Boardman, and McQueen).

##### *8.2.3 Vanderbilt University Medical Center's BioVU*

We would like to thank Lea Davis for providing access to the Vanderbilt University Medical Center Biobank (BioVU). The dataset(s) used for the analyses described were obtained from BioVU which is supported by numerous sources: institutional funding, private agencies, and federal grants. These include the NIH funded Shared Instrumentation Grant S10RR025141; and CTSA grants UL1TR002243, UL1TR000445, and UL1RR024975. Genomic data are also supported by investigator-led projects that include U01HG004798, R01NS032830, RC2GM092618, P50GM115305, U01HG006378, U19HL065962, R01HD074711; and additional funding sources listed at https://victr.vumc.org/biovu-funding/.

##### *8.2.4 COGA*

COGA: The Collaborative Study on the Genetics of Alcoholism (COGA), Principal Investigators B. Porjesz, V. Hesselbrock, T. Foroud; Scientific Director, A. Agrawal; Translational Director, D. Dick, includes eleven different centers: University of Connecticut (V. Hesselbrock); Indiana University (H.J. Edenberg, T. Foroud, J. Nurnberger Jr., Y. Liu); University of Iowa (S. Kuperman, J. Kramer); SUNY Downstate (B. Porjesz, J. Meyers, C. Kamarajan, A. Pandey); Washington University in St. Louis (L. Bierut, J. Rice, K. Bucholz, A. Agrawal); University of California at San Diego (M. Schuckit); Rutgers University (J. Tischfield, A. Brooks, R. Hart); The Children's Hospital of Philadelphia, University of Pennsylvania (L. Almasy); Virginia Commonwealth University (D. Dick, J. Salvatore); Icahn School of Medicine at Mount Sinai (A. Goate, M. Kapoor, P. Slesinger); and Howard University (D. Scott). Other COGA collaborators include: L. Bauer (University of Connecticut); L. Wetherill, X. Xuei, D. Lai, S. O'Connor, M. Plawecki, S. Lourens (Indiana University); L. Acion (University of Iowa); G. Chan (University of Iowa; University of Connecticut); D.B. Chorlian, J. Zhang, S. Kinreich, G. Pandey (SUNY Downstate); M. Chao (Icahn School of Medicine at Mount Sinai); A. Anokhin, V. McCutcheon, S. Saccone (Washington University); F. Aliev, P. Barr (Virginia Commonwealth University); H. Chin and A. Parsian are the NIAAA Staff Collaborators.

We continue to be inspired by our memories of Henri Begleiter and Theodore Reich, founding PI and Co-PI of COGA, and also owe a debt of gratitude to other past organizers of COGA, including Ting-Kai Li, P. Michael Conneally, Raymond Crowe, and Wendy Reich, for their critical contributions. This national collaborative study is supported by NIH Grant U10AA008401 from the National Institute on Alcohol Abuse and Alcoholism (NIAAA) and the National Institute on Drug Abuse (NIDA).

##### *8.2.5 The Externalizing Consortium*

The Externalizing Consortium gratefully acknowledges Aysu Okbay and the SSGAC. Initial analyses by the Externalizing Consortium were funded by the National Institute of Alcohol Abuse and Alcoholism through an administrative supplement to R01AA015146 (DMD). Additional funding for investigator effort has been provided by K02AA018755, U10AA008401 (COGA), and P50AA022537 (to DMD), a European Research Council Consolidator Grant (647648 EdGe, to PK). The study was classified as secondary research of de-identified subjects, and the study was awarded ethical approval by the internal review board (IRB) of Virginia Commonwealth University (VCU), with reference number HM20019386.

##### *8.2.6 The Psychiatric Genomics Consortium's Substance Use Disorders (PGC-SUD) working group*

The Psychiatric Genomics Consortium's Substance Use Disorders (PGC-SUD) working group is supported by MH109532 with funding from NIMH and NIDA. We gratefully acknowledge prior support from NIAAA and thank all our contributing investigators and study participants who make this research possible.

##### *8.2.7 UK10K Consortium*

This study makes use of data generated by the UK10K Consortium, derived from samples from the UK10K ALSPAC Cohort (EGAD00001000740) and the UK10K TwinsUK Cohort (EGAD00001000741). A full list of the investigators who contributed to the generation of the data is available from www.UK10K.org. Funding for UK10K was provided by the Wellcome Trust under award WT091310.

##### *8.2.8 UK Biobank (UKB)*

This research has been conducted using the UK Biobank Resource under Application Number 40830 and 11425. Informed consent was obtained from UK Biobank subjects.

##### *8.2.9 Philadelphia Neurodevelopmental Cohort (PNC)*

This study used data from PNC, acquired through dbGaP (accession number phs000607.v3.p2). Support for the collection of the data for PNC was provided by grant RC2MH089983 awarded to Raquel Gur and RC2MH089924 awarded to Hakon Hakonarson. Subjects were recruited and genotyped through the Center for Applied Genomics (CAG) at The Children's Hospital in Philadelphia (CHOP). Phenotypic data collection occurred at the CAG/CHOP and at the Brain Behavior Laboratory, University of Pennsylvania.
