## Supplementary Figures for "Multivariate genomic analysis of 1.5 million people identifies genes related to addiction, antisocial behavior, and health"

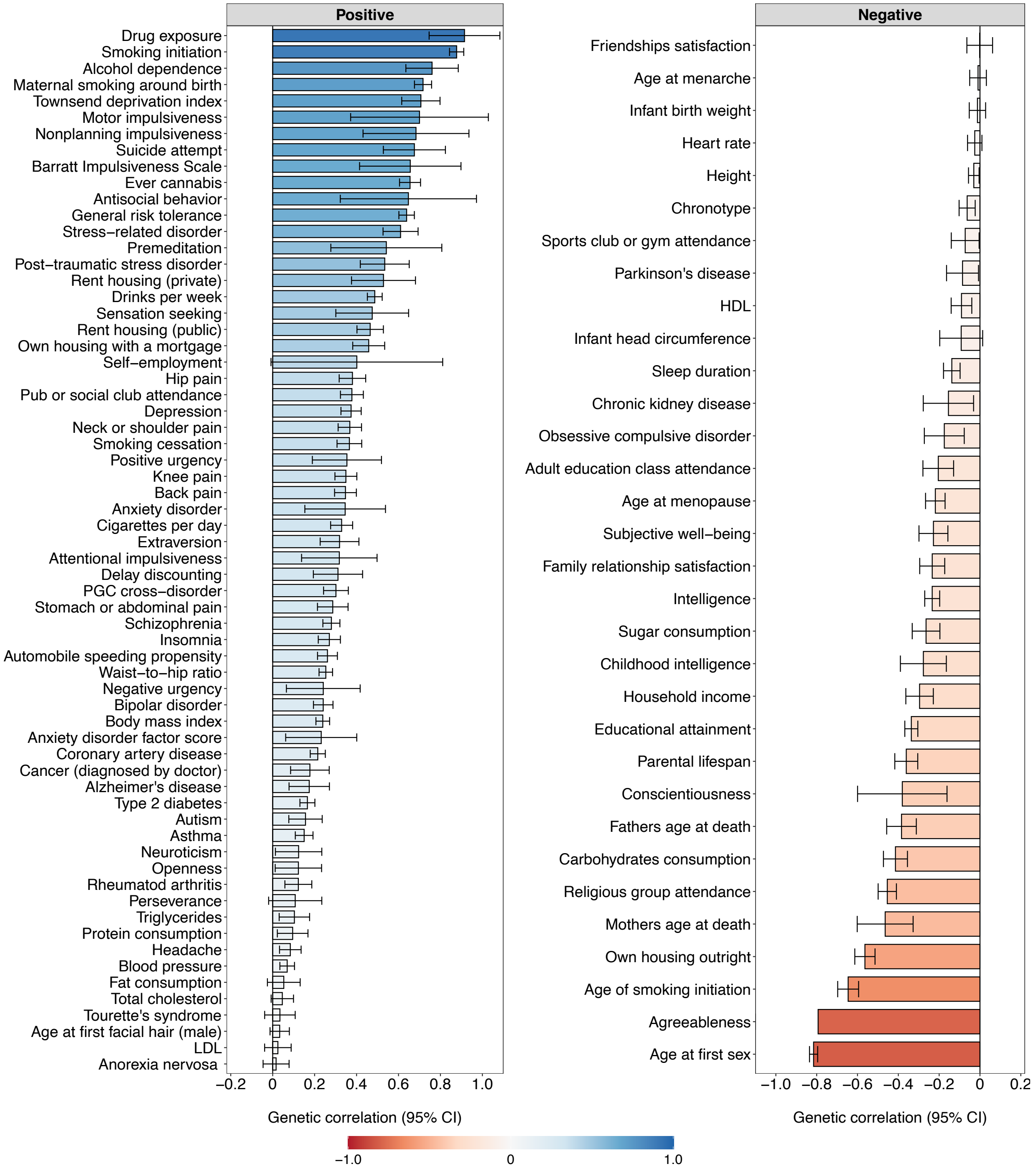


**Extended Data Figure 1 | Genetic correlations.** Bar plot of genetic correlations ($r_g$) estimated with Genomic SEM between the latent genetic externalizing factor with 92 other complex traits (**Supplementary Information section 3**). Error bars are 95% confidence intervals (omitted for Agreeableness). The estimates are also reported in **Supplementary Table 8**.

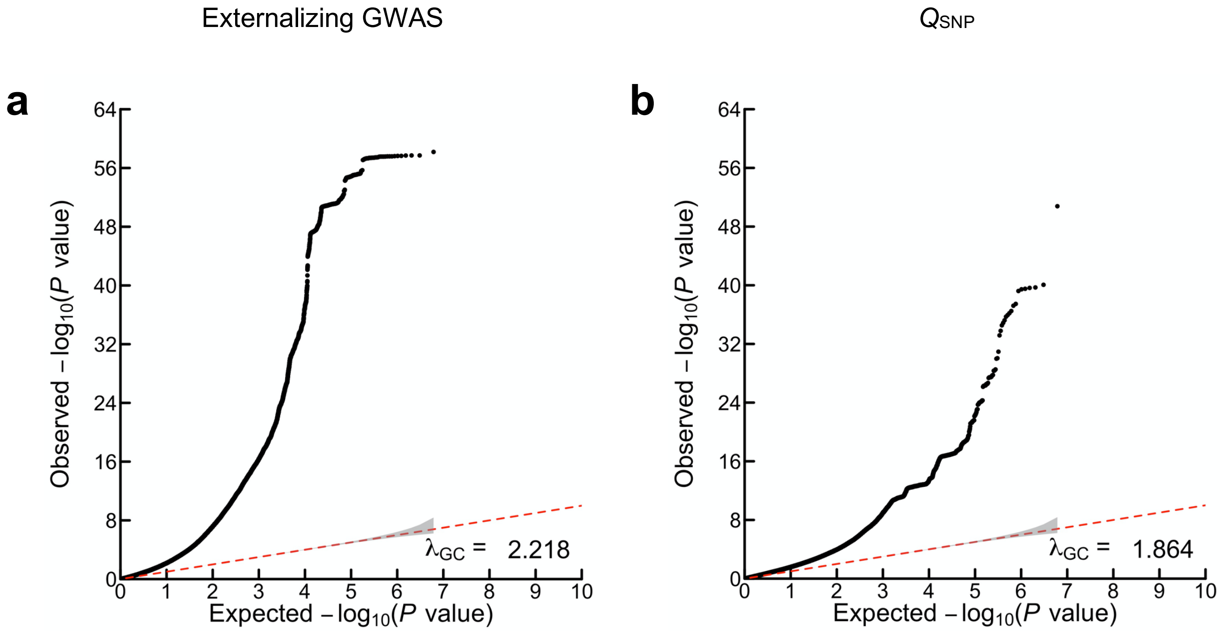


**Extended Data Figure 2 | Quantile-quantile (Q-Q) plots of the externalizing GWAS and $Q_{SNP}$ results.** The panels display Q-Q plots for (**a**) the externalizing GWAS ($N_{eff}$ = 1,492,085) and (**b**) SNP-level tests of heterogeneity ($Q_{SNP}$) with respect to the SNP-effects estimated in the externalizing GWAS (for more details see **Supplementary Information section 3**). The y-axis is the observed association *P* value on the $-\log_{10}$ scale (based on a two-tailed hypothesis test). The gray shaded areas represent 95% confidence intervals under the null hypothesis. The genomic inflation factors displayed here, $\lambda_{GC}$, is defined as the median $\chi^2$ statistic of the association test statistic divided by the expected median of the $\chi^2$ distribution with 1 degree of freedom, and were calculated with 6,132,068 and 6,107,583 SNPs for (**a**) and (**b**), respectively. Although there is a noticeable early "lift-off", the estimated LD Score regression intercepts of (**a**) 1.115 (*SE* = 0.019) and (**b**) 0.9556 (*SE* = 0.013) suggest that most of the inflation of the test statistics is attributable to polygenicity rather than bias from population stratification (*13*, *23*).

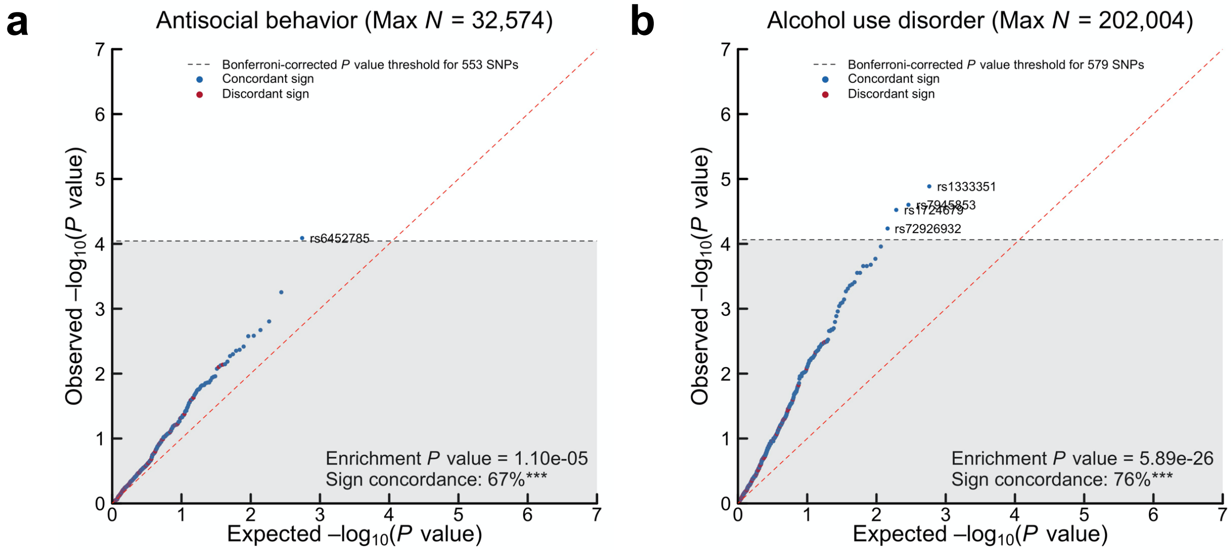


**Extended Data Figure 3 | Quantile-quantile (Q-Q) plots of the proxy-phenotypes analyses.** Panels (**a–b**) show –log10($P$) values for the 553 and 579 jointly associated lead SNPs (or such SNPs that could be proxied in case of missingness, $r^2 > 0.8$) that were looked up in independent, second-stage GWAS samples on (1) antisocial behavior ($N$ = 32,574) and (2) alcohol use disorder ($N$ = 202,400), respectively (**Supplementary Information section 4**). Dashed line denotes experiment-wide significance at $P < 0.05/553$ and 0.05/579 for (1) and (2), respectively. Enrichment $P$ value is the result of a one-tailed test of joint enrichment for association against an empirical null distribution of 138,250 and 144,750 near-independent ($r^2 < 0.1$) SNPs, matched on MAF, that were randomly selected from the GWAS on (1) and (2), respectively. Sign concordance is the proportion of looked-up SNPs with concordant direction of effect sizes across the externalizing GWAS and the second-stage GWAS. *** denotes $P < 0.001$.

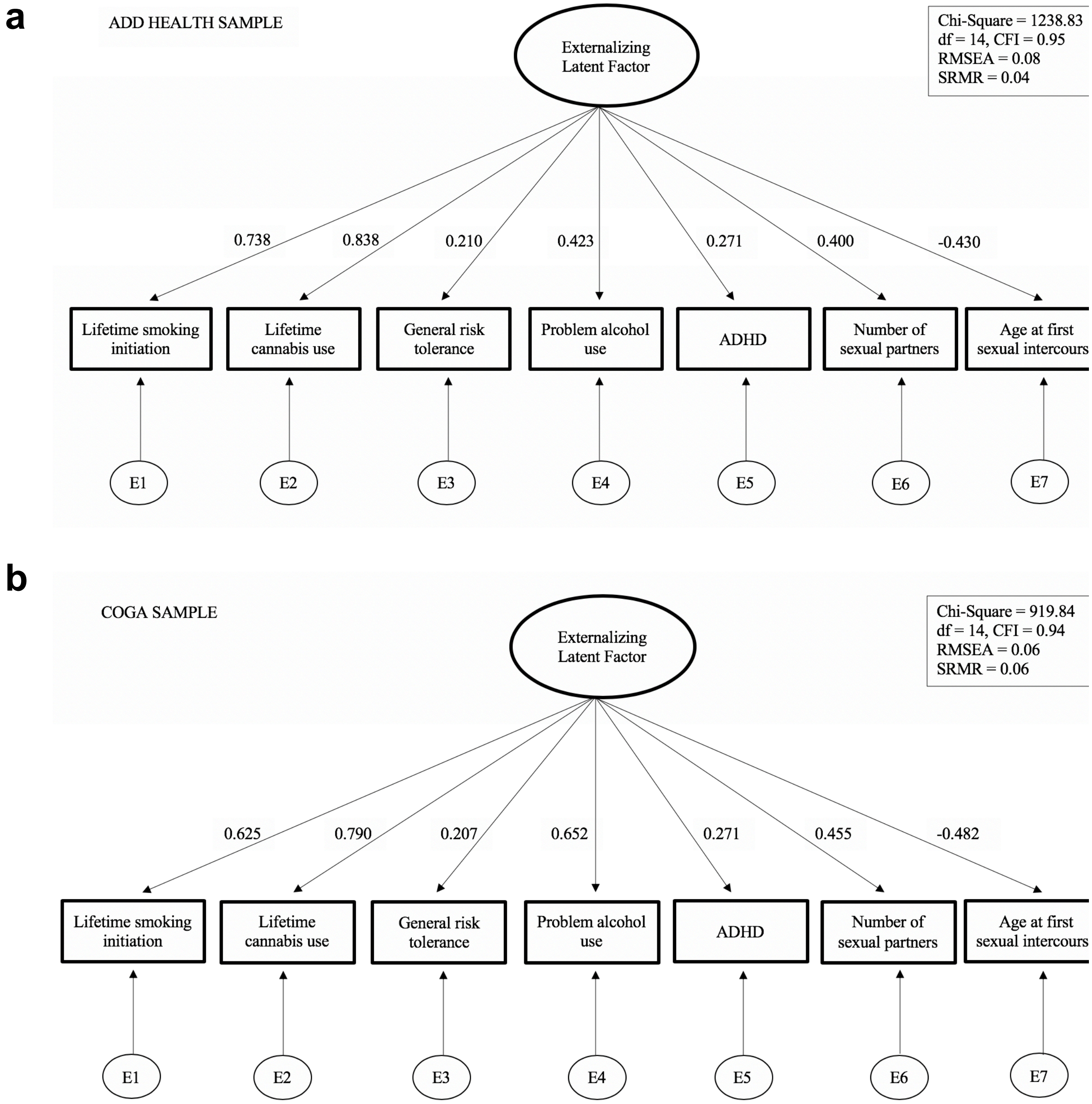


**Extended Data Figure 4 | Confirmatory factor analysis in Add Health and COGA.** Results of the confirmatory factor analysis (CFA) models in (**a**) Add Health and (**b**) COGA, in $N$ = 15,107 and 16,857 individuals, respectively (**Supplementary Information section 5**). Model fit statistics and fit indices are degrees of freedom (df), comparative fit index (CFI), root mean square error (RMSEA), standardized root mean squared residual (SRMR). Standardized factor loadings presented as numbers on the paths.

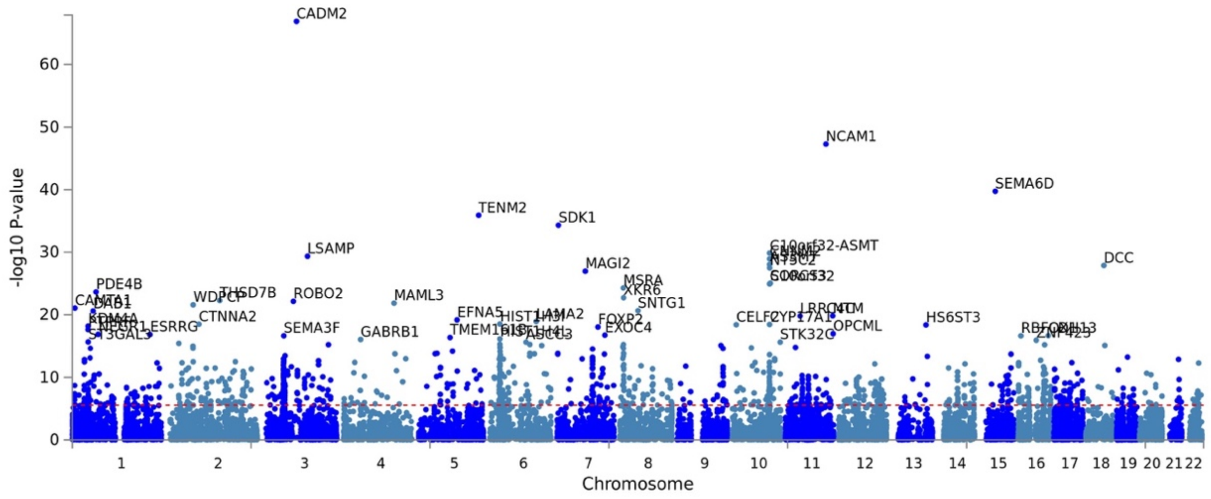


**Extended Data Figure 5 | MAGMA gene-based association analysis.** Manhattan plot of the –log10($P$) of 18,093 genes that were tested for association in the MAGMA gene-based association analysis with a one-tailed test (**Supplementary Information section 6**). The 50 most significant genes are labeled with gene names. Red dashed line represents Bonferroni-significance adjusted for the number of tested genes (one-tailed $P = 2.76{\times}10^{-6}$). 1,020 genes were found to be significant, of which 253 overlapped with the nearest genes annotated to the 579 jointly associated SNPs.

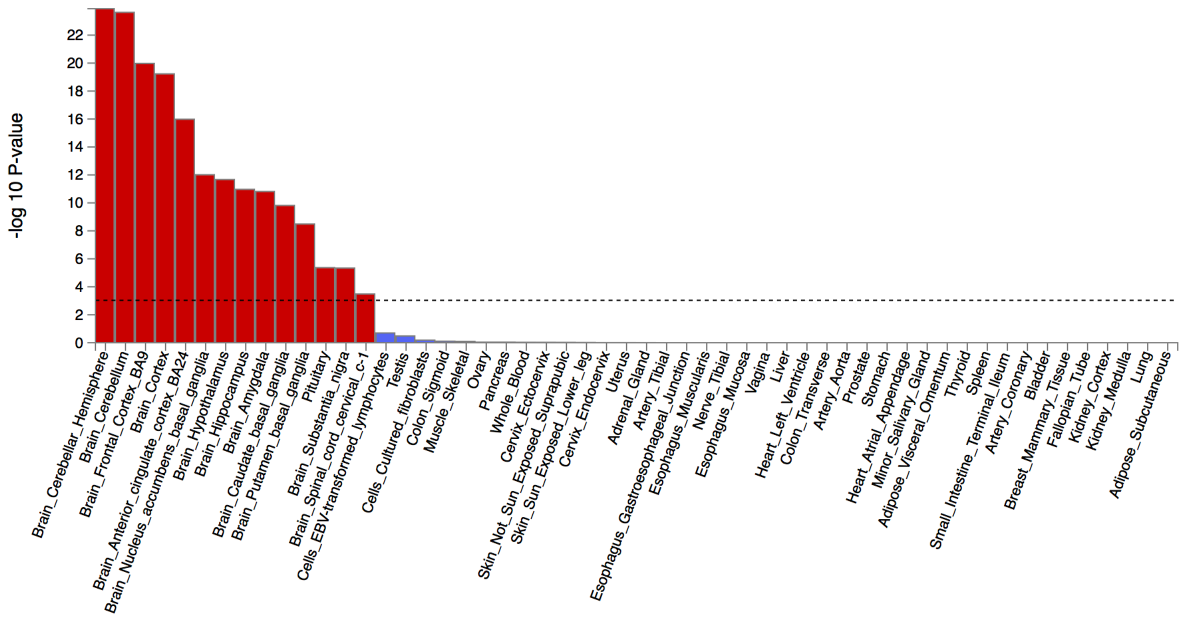


**Extended Data Figure 6 | MAGMA gene-property analysis.** The analysis identified that the externalizing GWAS is significantly enriched in brain and pituitary gland tissues (**Supplementary Information section 6**). Dashed line denotes Bonferroni-corrected significance, adjusted for testing 54 tissues (one-tailed $P < 9.26\times10^{-4}$). 14 tissues were significantly associated with the externalizing GWAS, including 13 brain related tissues and the pituitary tissue. The results are also report in **Supplementary Table 23**.

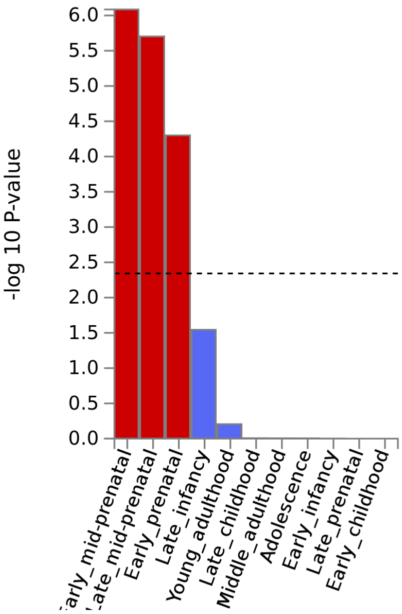


**Extended Data Figure 7 | MAGMA gene-property analysis of enrichment in brain tissues across 11 developmental stages (BrainSpan).** The analysis identified that the externalizing GWAS is significantly enriched in during prenatal developmental stages (**Supplementary Information section 6**). Dashed line denotes Bonferroni-corrected significance, adjusted for testing 54 tissues (one-tailed $P < 9.26{\times}10^{-4}$). The results are also report in **Supplementary Table 24**.

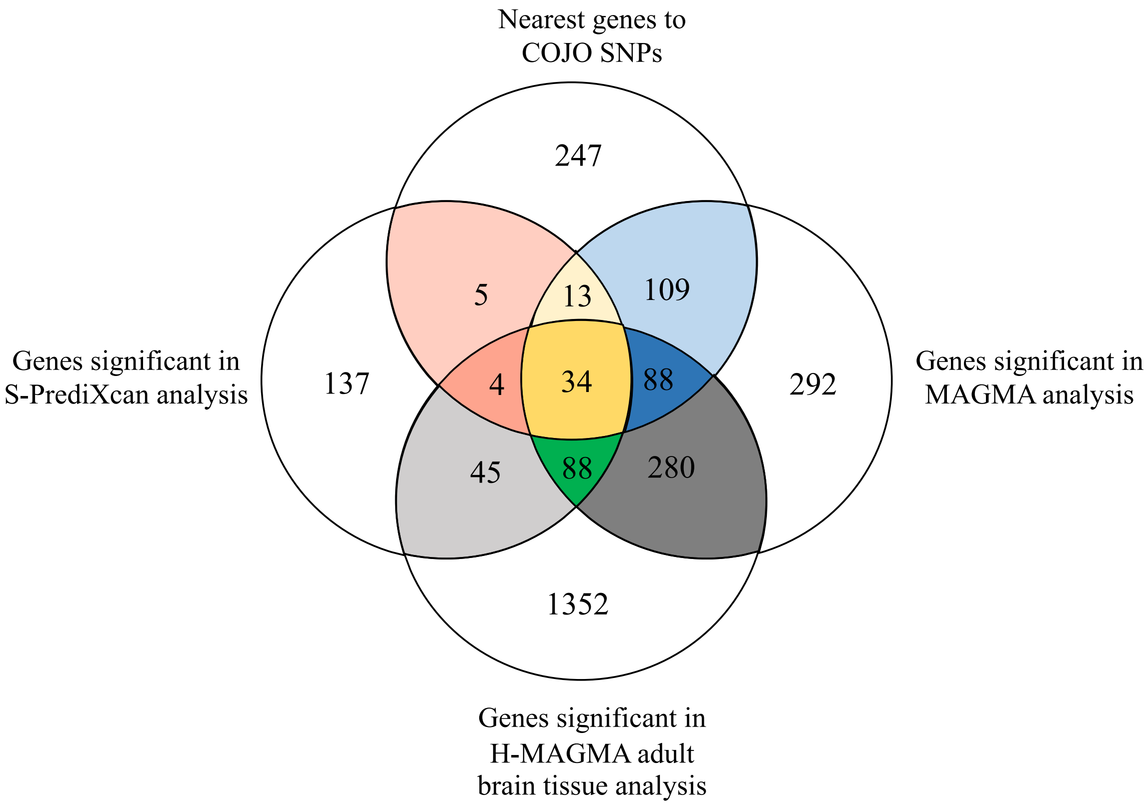


**Extended Data Figure 8 | Gene overlap across multiple gene-association methods.** This Venn diagram illustrates the overlap between the nearest genes to the 579 jointly associated lead SNPs (denoted as COJO SNPs), the genes significant in the MAGMA gene-based analysis (**Supplementary Table 21**), the genes significant in the H-MAGMA adult brain tissue analysis (**Supplementary Table 25**), and the genes significant in the S-PrediXcan analysis (**Supplementary Table 29**). Across the analyses, 34 genes were consistently implicated; these genes include *CADM2*, *PACSIN3*, *ZIC4*, *MAPT*, and *GABRA2*. Colored regions of this diagram correspond to the coloring shown in **Supplementary Table 30**, which lists all identified genes.
